# PINK1 stabilizes RMDN3-VAPB interaction without altering the proteomic landscape of mitochondria-ER contact sites

**DOI:** 10.64898/2026.07.21.739757

**Authors:** Alina Rühmkorf, Alexander Zorn, Jerome Duschek, Maria Georgina Herrera, Sophie Kratzert Levantovsky, Regina Feederle, Fereniki Moschogiannaki, Konstanze F. Winklhofer, Christian Behrends, Angelika Bettina Harbauer

## Abstract

Mitochondria-ER contact sites (MERCS) have emerged as central hubs of organellar biology, participating in the exchange of lipids, Ca^2+^ and even protein precursors. During mitochondrial dysfunction, the contact between the ER-resident protein VAPB and the outer mitochondrial membrane protein RMDN3 (PTPIP51) increases, yet the mechanism stabilizing this connection despite ongoing mitophagy remains to be determined. Here we develop a splitAPEX2 based proximity biotinylation tool to assess the interaction of VAPB and RMDN3. We determine that activation of the mitochondrial depolarization-induced kinase PINK1 leads to phosphorylation of RMDN3 at T160, which is necessary to increase the interaction with VAPB in the presence of mitochondrial depolarization. Mass Spectrometric analysis of the RMDN3-VAPB MERCS proteome reveals that these contacts also include peroxisomal proteins and that the proteomic composition remains stable during mitochondrial depolarization.

## Introduction

Contact sites between the outer mitochondrial membrane (OMM) and the endoplasmic reticulum (ER) form through specific interactions between distinct protein tether pairs^1^. Originally discovered as a single protein complex in yeast^2^, the repertoire of proteins tethering MERCS has diversified in mammals to include interactions between Mitofusin1/2 (Mfn1/2), VAPB-RMDN3 (PTPIP51), Fis1-Bap31, PDZD8-FKBP88, IP_3_R-VDAC, SYNJ2P-ESYT1 and SYNJ2BP-RRBP1^3–8^, with frequent new additions. The advent of proximity biotinylation has sped up the discovery of new interactors mediating MERCS in different approaches. For example, promiscuously biotinylating enzymes can be targeted to either the ER or the OMM, and the subsequent intersection of the derived local proteomes has revealed new interaction partners^8^. Further refinement of the specificity of this method can be achieved by combination with organellar isolation^9^, or by expressing a split version of TurboID that only activates upon close apposition of the two membranes^10,11^. Alternatively, the proximity of ER and mitochondria can reconstitute a split version of GFP tethered to the respective membranes, which is then subsequently detected by an intracellular nanobody coupled to the peroxidase APEX2^11^. As these approaches used generic OMM and ER transmembrane sequences, baseline proximity biotinylation was generally low and in many cases needed to be enhanced by inclusion of a drug-inducible heterodimerizer. While these approaches have given insights into the technical possibilities that enable the identification of new MERCS, they lack the specificity to selectively label one kind of MERCS over another. This is a critical limitation that prevents our understanding of the individual functions of MERCS that are not shared between tether pairs.

One functional difference between MERCS is the response to mitochondrial damage. Initial electron microscopic characterization of MERCS revealed that the distance between mitochondria and ER increased upon addition of the mitochondrial uncoupler CCCP^12^. Mechanistically, this was mediated by the stabilization of the depolarization-sensitive kinase PTEN-induced kinase 1 (PINK1), which phosphorylates the Mfn2 tether, initiating its ubiquitination-dependent extraction from the membrane and proteasomal degradation^12,13^. Similar ubiquitination and degradation mechanisms target other OMM tethering proteins, including SYNJ2BP and VDAC^14–16^, supporting the observed overall decrease in MERCS abundance upon mitochondrial depolarization and mitophagy induction. However, mitophagy is an extreme response to mitochondrial dysfunction. Thus, a temporary maintenance of organellar contact sites could be necessary to enable other mitigation attempts, including the shuttling of reactive oxygen species (ROS) to peroxisomes^17^ and the replacement of damaged mitochondrial proteins by increased protein translation of e.g. OXPHOS proteins in the vicinity of damaged organelles sensed by PINK1^18^. Especially in neurons, the long distances spanned by axons and dendrites provide an additional challenge for mitochondrial homeostasis^19^. We recently identified PINK1 as a substrate of the ER-associated import pathway that relies on transfer of preproteins between the two organelles at MERCS^20,21^. Thus, maintaining MERCS during the initial phase of mitochondrial dysfunction may provide a transient platform for local PINK1 accumulation and signaling, before damaged mitochondrial become committed to degradation. Indeed, PINK1 has been reported to relocalize to MERCS upon mitochondrial damage, where it promotes MERCS formation^22^. Intriguingly, it was recently reported that the contact site between VAPB and RMDN3 increases upon mitochondrial intoxication with inhibitors of the respiratory chain due to ROS^23^. This was dependent on phosphorylation of RMDN3 within its FFAT motif, which requires phosphorylation to enable efficient binding to the major sperm protein (MSP) domain of VAPB^24^. However, the kinase mediating this phosphorylation remained unidentified.

Here we develop a splitAPEX2-based proximity-labelling approach^25^ that reports specifically on the MERCS formed by RMDN3 and VAPB. We further identify PINK1 as the kinase mediating phosphorylation of RMDN3 in its FFAT motif to stabilize this contact site during mitophagy induction. Surprisingly, in spite of the ongoing degradation of OMM proteins upon mitophagy induction, the proteomic environment of RMDN3-VAPB interaction remains stable upon acute mitochondrial depolarization, suggesting it provides a safe haven of continued functionality during mitochondrial stress. These data expose an unexplored complexity of MERCS-specific responses to mitochondrial dysfunction.

## Results

### RMDN3-VAPB SplitAPEX2 in HEKs and neurons is localized to MERCS

To enable the specific proximity biotinylation of MERCS formed by RMDN3 and VAPB, we created expression constructs that included the entire open reading frame for RMDN3 and VAPB, and added the N-terminal and C-terminal halves of splitAPEX (termed AP and EX)^25^, respectively. To visualize expression and localization of our constructs, we also included GFP or mCherry and the small affinity tags HA and V5 to create RMDN3-GFP-V5-AP (abbreviated as RMDN3-AP) and EX-HA-mCherry-VAPB (abbreviated as EX-VAPB). Both constructs expressed well in HEK293 cells and displayed robust biotinylation of proteins (Fig. 1A) after treatment with biotin-phenol for 30 min, followed by a 1 min treatment with H_2_O_2_ to activate APEX2-mediated peroxidation of biotin-phenol. The resulting biotin-phenoxyl radical covalently binds to neighboring proteins during this short one-minute interval. This reaction is immediately followed by quenching to prevent spreading of the signal and H_2_O_2_-induced signaling^26^. The advantage of using APEX2 is the ability to also visualize the activity of the peroxidase in electron microscopy (EM) by treatment with DAB (3,3’-diaminobenzidine)^27^. This creates a DAB polymer that readily stains with electron-dense osmium tetroxide. HEK293 cells expressing RMDN3-AP and EX-VAPB displayed a strong DAB signal at MERCS (Fig. 1B, black arrows), indicating correct localization of the reconstituted split enzyme. Importantly, despite overexpression of both constructs, not all mitochondria nor the entire OMM were positive for DAB (Fig. 1B, white arrows), indicating that the diversity of MERCS remained intact. In order to validate expression and localization in a different cell type, we also transiently transfected cultured hippocampal neurons with RMDN3-AP and EX-VAPB, which we subjected to the same labelling protocol. Staining against the V5 and HA tags revealed faithful localization of RMDN3-AP and EX-VAPB to mitochondria and the ER, respectively. Anti-biotin staining detected biotinylated proteins in both, soma and neurites with signal enriched at areas of overlap between the two organelles (Fig. 1C). This was particularly visible in neurites due to the lower density of mitochondria in this area (Fig. 1C, second row). Omission of biotin-phenol only yielded background labelling due to endogenously biotinylated proteins, without the clear localization to mitochondria (Fig. 1C, lower panels). Similar to HEK cells, DAB-positive signal was detected specifically at MERCS also in cultured neurons (Fig. 1D, black arrows), but again not all MERCS were positive for DAB (Fig. 1D, white arrows). Thus, the addition of AP and EX to an existing MERCS tether yielded the expected localization of the biotinylation signal in several cellular models.

**Figure 1:**
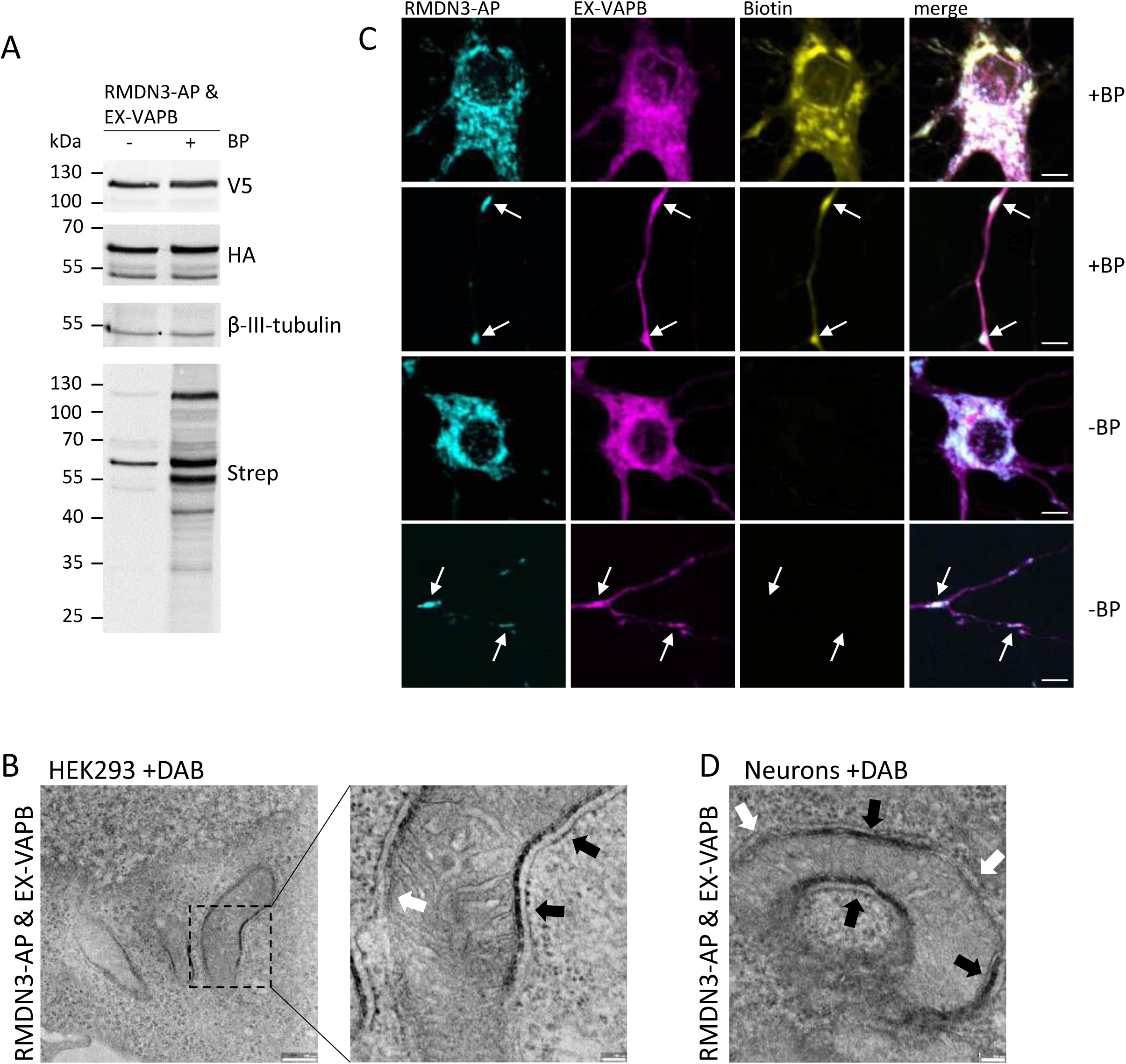
RMDN3-VAPB splitAPEX localizes to ER-mitochondria contact sites in HEK293 cells and primary neurons. A) Validation of RMDN3-VAPB splitAPEX activity in HEK293 cells. Cells expressing RMDN3-GFP-V5-AP and EX-HA-mCherry-VAPB were incubated in the presence or absence of biotin-phenol (BP). Biotinylated proteins were detected by streptavidin-staining. Expression of splitAPEX constructs was verified using antibodies against the V5 and HA tags. Scale bar 5 µm. B) Transmission electron microscopy of HEK293 cells expressing RMDN3-AP and EX-VAPB following DAB staining and osmium tetroxide processing. Electron-dense deposits are detected at subsets of mitochondria-ER contact sites (black arrows). White arrows indicate mitochondria-ER interfaces lacking detectable electron-dense staining. Scale bar 100 nm. C) Immunofluorescence analysis of primary hippocampal neurons expressing RMDN3-AP and EX-VAPB in the presence of absence of BP. RMDN3-AP is shown in cyan by staining for its V5 tag, EX-VAPB is shown in magenta by staining for its HA tag, and biotinylated proteins are shown in yellow by anti-biotin staining. Merged images are shown in the right panel. Arrows indicate regions of biotin labeling at sites of overlap between the two splitAPEX fragments. D) Transmission electron microscopy of primary hippocampal neurons expressing RMDN3-AP and EX-VAPB following DAB staining. Electron-dense deposits are observed at subsets of mitochondria-ER contact sites (black arrows), whereas other closely apposed mitochondrial and ER membranes remain unstained (white arrows). Scale bar 100 nm.

### Proteomic characterization of RMDN3-VAPB MERCS

To determine the proteomic composition of RMDN3-VAPB mediated MERCS, we isolated biotinylated proteins from both mock and biotin-phenol treated HEK293 cells and determined their identity by Mass Spectrometry after on-bead digestion with trypsin. Intriguingly, this approach highly enriched already established mitochondria-ER tether proteins (Fig. 2A). In total, we identified 787 significantly enriched proteins; subsequent cellular component enrichment analysis revealed a significant overrepresentation of mitochondria and ER proteins (Fig. 2B; Supplementary Table S1a). When compared to a previously published dataset using nanobody-targeted APEX^28^, 162 proteins were shared between both datasets, of which 50 proteins were ER-associated, and 78 mitochondria-associated (Fig. 2C, Supplementary Table S1b). This provides the specific subset of MERCS proteins present at MERCS specifically formed by RMDN3 and VAPB. Despite the cytosolic orientation of the reconstituted splitAPEX, we also recovered a substantial fraction of inner mitochondrial membrane proteins and ER luminal proteins (Fig. 2B). This is a commonly observed phenomenon when using APEX2 based detection^28^, which may be due to H_2_O_2_-induced membrane damage, allowing access of biotin-phenoxyl radicals to adjacent compartments.

**Figure 2:**
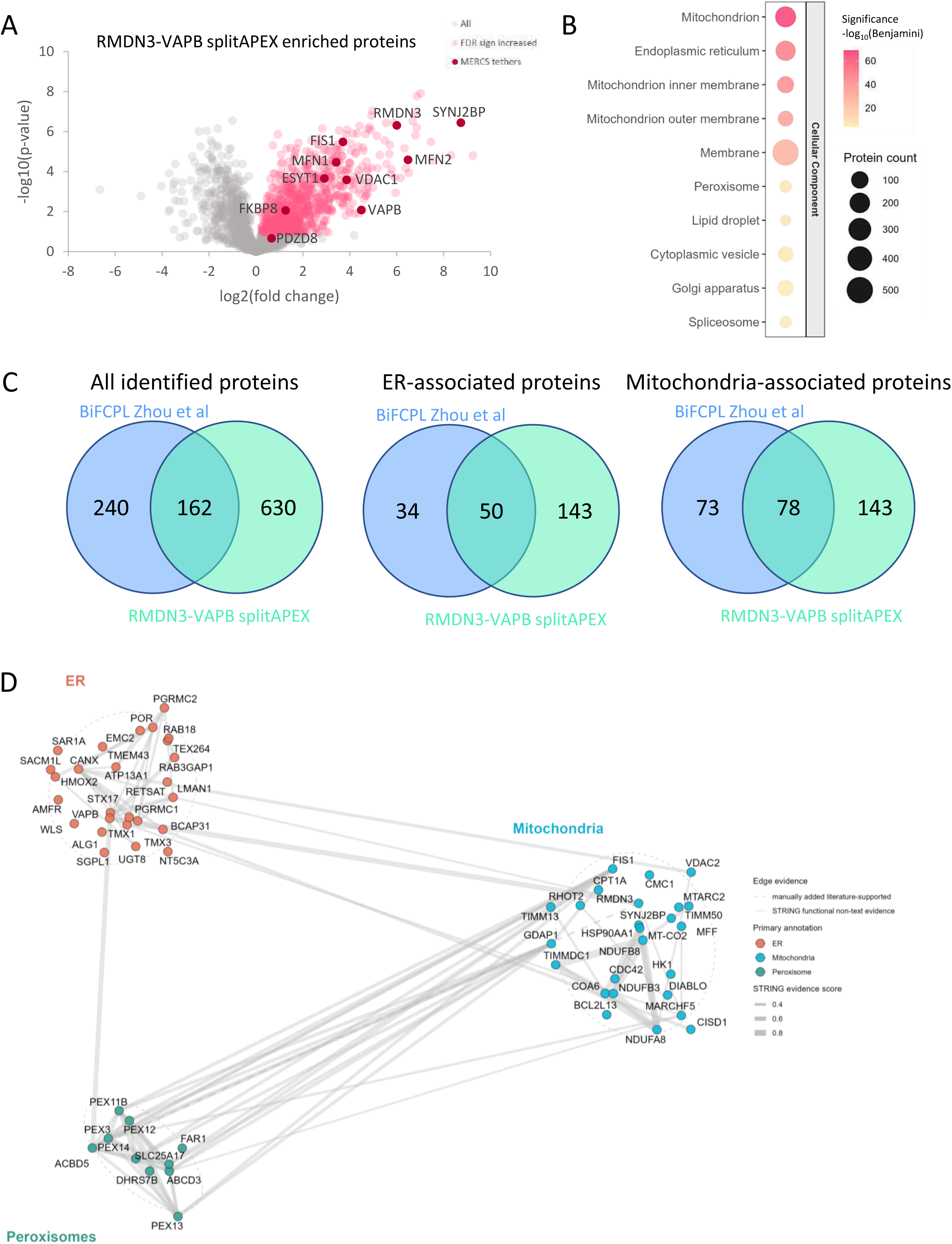
Proteomic characterization of the RMDN3-VAPB contact site. **A)** Volcano plot of proteins identified by RMDN3-VAPB splitAPEX proximity labeling. Proteins significantly enriched compared to controls are highlighted in red (FDR < 0.05). Known mitochondria-ER tethering proteins are indicated by dark red symbols. **B)** Bubbleplot showing significantly enriched GO-DAVID cellular component (CC) terms among proteins significantly enriched in the RMDN3-VAPB splitAPEX pulldown. Bubble size indicates the number of proteins assigned to each term, and color indicates statistical significance. **C)** Venn-diagrams showing the overlap between proteins found in the RMDN3-VAPB splitAPEX and proteins identified by bimolecular fluorescence complementation-based proximity labeling (BiFCPL)^28^. Comparisons were performed for all proteins identified, as well as ER-associated and mitochondria-associated proteins separately. **D)** STRING-based protein interaction network of selected organelle-associated proteins identified in the RMDN3-VAPB splitAPEX proteome. Up to 25 proteins with the highest log₂ fold enrichment were selected from each indicated organelle category. Nodes are colored according to their assigned cellular compartment. Edges represent protein associations retrieved from the STRING database. Dashed edges indicate manually added interactions that have been experimentally reported but were not represented in the STRING network^17^. Proteins assigned to more than one cellular compartment were categorized according to the prioritization scheme described in the methods.

Intriguingly, while ER and mitochondria represented the most significantly enriched cellular compartments in the RMDN3-VAPB splitAPEX proteome (Fig. 2B), we also detected proteins assigned to other ER-derived compartments such as peroxisomes and lipid droplets. STRING-based visualization of selected organelle-associated hits revealed tight links not only between the mitochondria and the ER as expected, but also pronounced interactions between mitochondria and peroxisomal proteins within our dataset (Fig. 2D). This included the interaction between RMDN3 and the peroxisomal protein ACBD5 which has recently been published to mediate transfer of reactive oxygen species (ROS) between mitochondria and peroxisomes^17^. ABCD5 also reportedly interacts with VAPB^29,30^. Thus, our data suggest that contacts between mitochondria and peroxisomes may overlap with ER-mitochondria contacts formed by RMDN3 and VAPB.

### RMDN3-VAPB interaction increases upon mitochondrial depolarization

Given the unique response of RMDN3-VAPB to mitochondrial depolarization^23^ and their connection to the detoxification of ROS via shuttling to peroxisomes, we sought to test the response of this contact site to mitochondrial dysfunction induced by inhibition of the respiratory chain. We compared the proximity of endogenous VAPB and RMDN3 in cultured neurons by proximity ligation assay (PLA), which uses oligonucleotide-coupled secondary antibodies that can be ligated and amplified only if their distance is less than 40 nm^31^. These proximity-dependent ligation events can then be detected and quantified as fluorescent puncta. PLA puncta were readily detected in cultured hippocampal neurons for both RMDN3-VAPB and another MERCS tether pair, SYNJ2BP and RRBP1 (Fig 3A). Interestingly, we observe a differential response: the abundance of SYNJ2BP-RRBP1 puncta in the soma decreased in response to complex III inhibition with Antimycin A (AA), compared with vehicle (EtOH) treated cells (Fig. 3B), whereas the number of PLA puncta for the RMDN3-VAPB interaction increased (Fig. 3C). To understand which response is the predominant effect on MERCS in neurons, we expressed a FRET-based indicator of ER-mitochondria proximity (FEMP)^32^ (Fig. 3D). We detected a reduced fluorescent life time of the donor fluorophore in the soma, which translates to an increased proximity between the two proteins (Fig. 3E). Finally, we tested whether also the splitAPEX based biotinylation response of RMDN3-AP and EX-VAPB was sensitive to mitochondrial depolarization. Indeed, we observed an increased level of biotinylated proteins in HEK cells expressing RMDN3-AP and EX-VAPB upon co-treatment with AA and Oligomycin A (OA) to prevent reverse proton transport by complex V in non-neuronal cells (Fig. 3F,G). This physiological response further supports the dynamic nature of the splitAPEX reporter.

**Figure 3:**
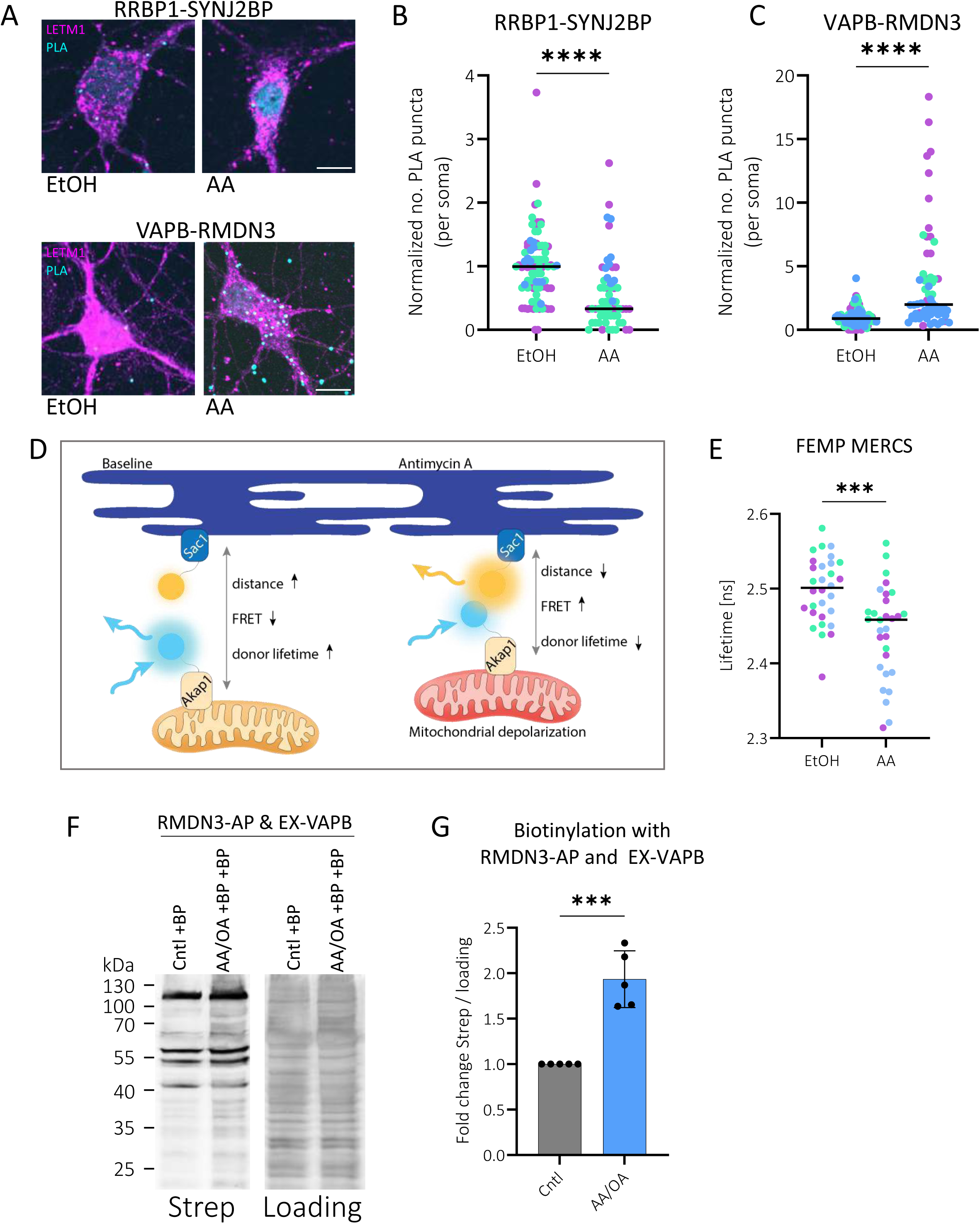
Mitochondrial damage differentially remodels MERCS. **A)** Representative images of PLA performed in primary hippocampal neurons detecting RRBP1-SYNJ2BP (top) or VAPB-RMDN3 (bottom) interactions following vehicle (EtOH) or Antimycin A (AA) treatment. Mitochondria were visualized by LETM1 immunostaining (magenta). PLA puncta are shown in cyan. Scalebar is 10 µm. **B, C)** Quantification of RRBP1-SYNJ2BP (B) and VAPB-RMDN3 (C) PLA puncta in vehicle- and AA-treated neurons. Values were normalized to the mean of the respective vehicle-treated control within each experimental batch. Colors indicate independent experimental batches (n=3), and black lines indicate the mean. Statistical analysis was performed using an unpaired t-test. **D)** Schematic representation of FEMP, a FRET-based mitochondrion-ER proximity sensor. mCerulean is targeted to mitochondria via the AKAP1 targeting sequence, while mVenus is targeted to the ER via the SAC1 targeting sequence. Close apposition of ER and mitochondria enables FRET from mCerulean to mVenus, resulting in reduced mCerulean donor lifetime. Thus, shorter donor lifetime indicates closer ER-mitochondria proximity. **E)** Fluorescence lifetime analysis of FEMP in cultured neurons, following vehicle or AA treatment. Three independent biological replicates were measured, with 10 cells in each replicate. Statistical analysis was performed using an unpaired t-test. **F)** Representative streptavidin blot of splitAPEX labeling in HEK293 cells expressing RMDN3-AP and EX-VAPB following treatment with vehicle (+BP) or Oligomycin A/Antimycin A (OA/AA +BP). Total protein staining is shown as a loading control. **G)** Quantification of the streptavidin signal shown in E after normalization to total protein staining. Statistical analysis was performed using an unpaired t-test.

### RMDN3-VAPB MERCS retain a stable proteome during mitochondrial depolarization

Prompted by the increase in RMDN3-VAPB interaction upon mitochondrial depolarization, and to determine whether depolarization also alters the molecular composition of RMDN-VAPB MERCS, we generated a new dataset comparing the biotinylated proteins upon RMDN3-AP and EX-VAPB expression in HEK293 cells that were subjected to AA/OA co-treatment for 1 h to control vehicle-treated cells. Upon AA/OA treatment, cells showed an increased phospho-ubiquitin signal in Western blot, confirming activation of mitophagy (Fig. 4A,B). Mass spectrometric analysis revealed a surprisingly stabile RMDN3-VAPB surrounding proteome, with only five hits increasing and seven hits decreasing in response to AA/OA treatment after false detection rate (FDR) correction (Fig. 4C, Supplementary Table S2). Among the significantly enriched proteins, oxidative stress sensor ACO2^33^ and the ER chaperone HSPA5 (BiP)^34^, as well as the metabolic enzymes ECHS1, DLST, and GCDH^35–37^. Proteins showing reduced RMDN3-VAPB proximity upon mitochondrial depolarization included the oxidoreductase TMX3^38^, the ER Ca^2+^ pump SERCA2 (ATP2A2)^39^, the apoptotic regulator BAX^40^, the heme biosynthesis associated enzyme CPOX^41^, the mitochondrial phosphate carrier SLC25A3, the translation elongation factor EEF1D^42,43^, and the spliceosome associated protein TFIP11^44,45^.

**Figure 4:**
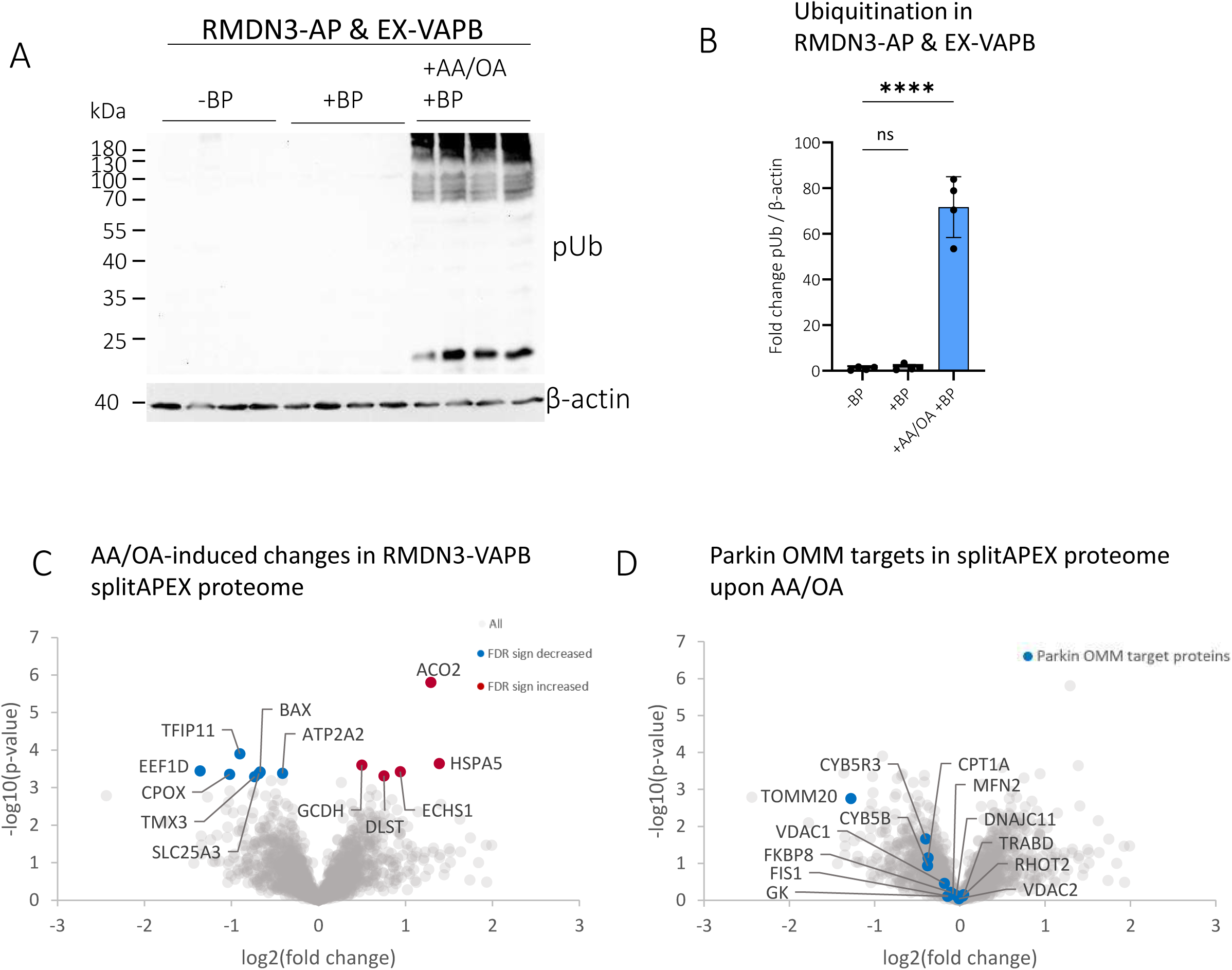
Effects of mitochondrial depolarization on the RMDN3-VAPB splitAPEX proteome. **A)** Representative western blot showing phospho-ubiquitin (pUb) levels in HEK293 cells expressing RMDN3-AP and EX-VAPB, following vehicle treatment and biotinylation (+BP) or combined OA/AA treatment with biotinylation (OA/AA +BP). Samples treated with vehicle and without biotin phenol (-BP) were included as controls. β-actin was included as a loading control. **B)** Quantification of phospho-ubiquitin (pUb) following AA/OA and biotin-phenol treatment from (A), normalized to β-actin. Values were normalized to the mean pUb/β-actin ratio of -BP condition. Statistical significance was assessed using one-way ANOVA followed by Dunnett’s multiple comparison test. **C)** Volcano plot comparing proteins identified by RMDN3-VAPB splitAPEX following OA/AA treatment and vehicle-treated +BP controls. Significantly increased or decreased proteins are shown in red or blue, respectively (FDR < 0.05). **C)** Same volcano plot as in B, with selected OMM Parkin-substrates highlighted and labeled. Selection of mitochondrial outer membrane proteins was based on increased ubiquitination in Parkin-expressing HeLa cells (WT-parkin_2h-values > 0.5), that are absent in inactive C431A-Parkin-expressing cells (C431A-parkin_2h-values < 0.5) as reported in Zittlau et al.^14^

To assess whether acute mitochondrial damage affected proteins linked to the PINK1/Parkin-pathway, we marked previously reported Parkin targets of the OMM in our dataset (Fig. 4D). While not reaching significance, the abundance of these proteins showed a trend towards a depletion upon induction of mitophagy. This may be due to the short treatment time that did not allow for extended degradation of the OMM proteome, which peaks at two to six hours after depolarization^14^. Overall, this data indicate that mitochondrial depolarization does not induce extensive remodeling of the RMDN3-VAPB proteome after 1 h, but rather results in selective changes affecting only a small subset of proteins.

### PINK1-mediated phosphorylation of RMDN3 at T160 enhances VAPB binding

Another way to stabilize the specific interaction between RMDN3 and VAPB during mitochondrial depolarization would be to promote their interaction by increased phosphorylation of RMDN3 in its phospho-FFAT motif at residue T160 (Fig. 5A)^23,24^. Phosphorylation at this site introduces a negative charge which can be mimicked by replacing threonine with the negatively charged amino acid aspartate. We confirmed by co-immunoprecipitation that the interaction between VAPB and RMDN3 was increased in T160D mutants, whereas mutations to alanine (T160A) or the longer, negatively charged amino acid glutamate (T160E) further reduced the interaction in HEK293 cells (Fig. 5B,C). In contrast, deletion of the coiled coil domain (RMDN3(ΔCC)), previously suggested to mediate the interaction^46^, had no effect on the interaction between RMDN3 and VAPB (Fig. 5B,C).

**Figure 5:**
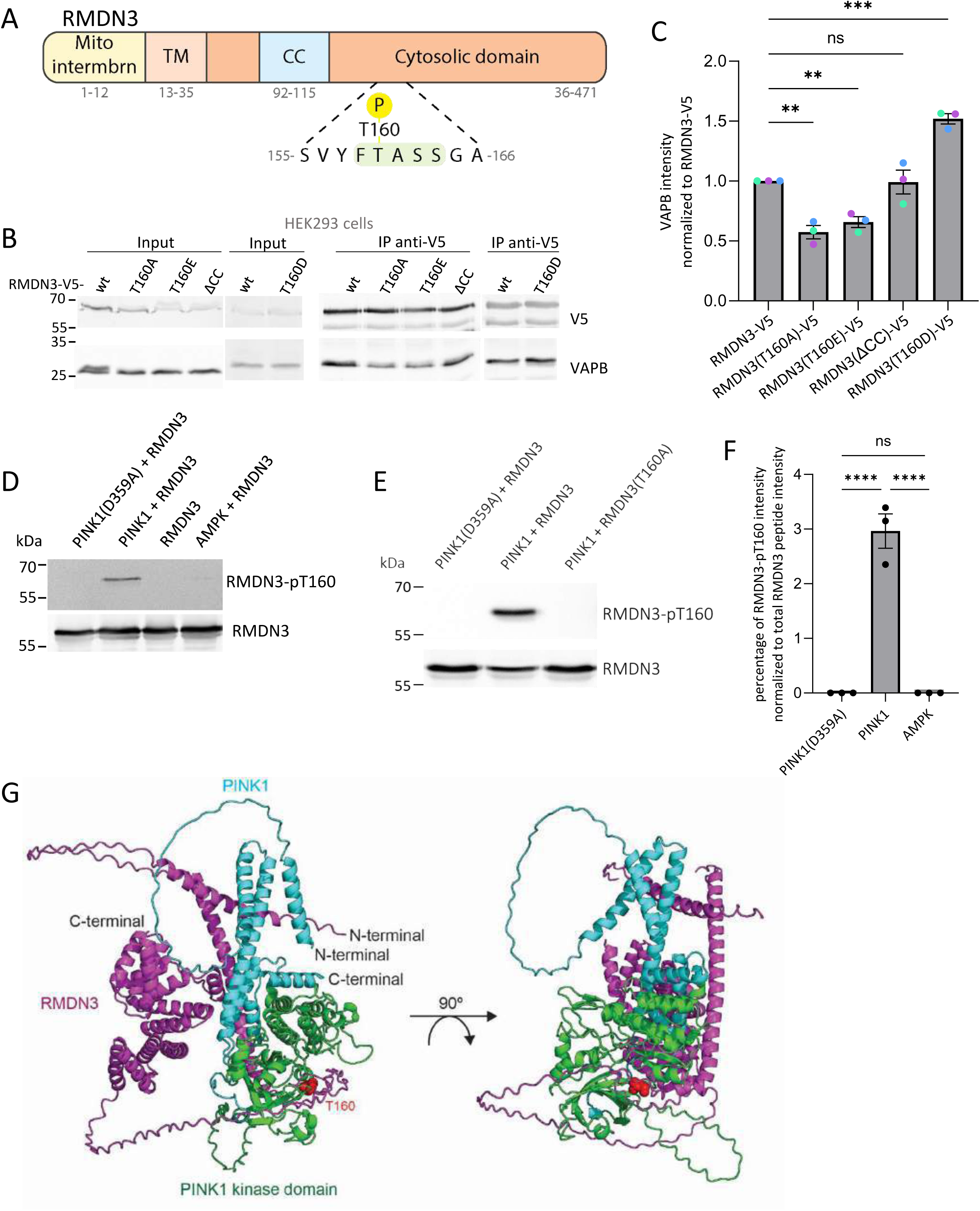
PINK1-mediated phosphorylation of RMDN3 at T160 enhances VAPB binding. **A)** Schematic representation of RMDN3 domain structure, including the mitochondrial intermembrane space-facing N-terminus (mito intermbrn), transmembrane domain (TM), coiled coil domain (CC) within the cytosolic domain that contains also the phospho-FFAT motif. **B)** Co-Immunoprecipitation (Co-IP) analysis of VAPB binding to V5-tagged RMDN3-T160 mutant variants. RMDN3-V5 was immunoprecipitated using anti V5-antibody and co-precipitated VAPB. RMDN3-T160 mutant variants included the phospho-ablative T160A, the phospho-mimetic T160E and T160D mutants, and an RMDN3 construct lacking the coiled coil domain(ΔCC). **C)** Quantification of Co-IP shown in (B). Co-immunoprecipitated VAPB signal was normalized to immunoprecipitated RMDN3-V5. Statistical analysis was performed using unpaired T-tests comparing each RMDN3-T160 mutant to wild-type RMDN3. **D)** *In vitro* kinase assay using recombinant RMDN3 incubated with active PINK1, kinase-dead PINK1 or active AMPK. Phosphorylation of RMDN3 at T160 was detected with a T160-phospho-specific antibody. Total RMDN3 is shown as loading control. **E)** *In vitro* kinase assay using recombinant WT or T160A mutant RMDN3 incubated with active or kinase-dead PINK1. RMDN3-T160 phosphorylation was detected with a T160-phospho-specific antibody. **F)** Phospho-proteomic analysis of recombinant RMDN3 following incubation with active or kinase-dead PINK1 or AMPK, showing relative phosphorylation of the T160-containing peptide exclusively following incubation with wild-type PINK1. Statistical analysis was performed using one-way ANOVA. **G)** Structure of the complex formed by human RMDN3 and human PINK1 predicted by AlphaFold3^49^. The analysis suggests that T160 from RMDN3 is in the vicinity of the PINK1 kinase domain when the complex forms. The predicted structure of RMDN3 is shown in magenta, and T160 is depicted as red spheres. The PINK1 kinase domain (residues 156–513) is shown in green, and the remainder of PINK1 is displayed in cyan.

One kinase known to be activated upon mitochondrial depolarization is PINK1. To test the hypothesis that PINK1 is the kinase responsible for RMDN3 phosphorylation upon mitochondrial depolarization, we raised a phospho-specific antibody against a phosphorylated peptide corresponding to T160 in RMDN3. We then subjected the purified recombinant cytosolic domain of RMDN3 to an *in vitro* kinase assay using recombinant PINK1 from the species *Tribolium castaneum* which retains activity *in vitro*^47^. As controls, we included a kinase dead version of PINK1 (D359A), as well as recombinant AMP-dependent kinase (AMPK), another kinase that gets activated during mitochondrial dysfunction^48^. Intriguingly, only active PINK1 was able to elicit a signal by the phospho-specific antibody, which was abolished when T160 was mutated to alanine (Fig. 5D,E). The specificity of the phosphorylation at this residue was also corroborated by mass spectrometry (Fig. 5F). Structural modeling using AlphaFold3^49^, predicts that the residue T160 is positioned in close proximity to the PINK1 kinase domain (Fig. 5G). This further supports a direct phosphorylation of RMDN3 by PINK1.

### PINK1 mediates damage-induced increase of the RMDN3-VAPB interaction

To validate the importance of PINK1 for the regulation of RMDN3-VAPB interaction upon mitochondrial depolarization, we performed PLA between VAPB and RMDN3 in cultured neurons in the presence of non-targeting or PINK1-targeting shRNAs. We co-transfected a mitochondrial marker to selectively image transfected neurons, taking advantage of the high co-transfection efficiency in neurons. As expected, treatment with AA increased the number of PLA puncta in control shRNA transfected neurons compared to control treatment. However, this increase was absent upon PINK1 knockdown (Fig. 6A,B). Co-immunoprecipitation experiments in N2A cells showed a similar effect of PINK1 knock-down. Intriguingly, not only the AA-induced increase in RMDN3-VAPB interaction was prevented in PINK1 knockdown cells, but also the increase of RMDN3 T160 phosphorylation as detected by the phospho-specific antibody. Together with the *in vitro* evidence for direct phosphorylation of RMDN3 by PINK1 (Fig. 5D,E), this supports a model in which phosphorylation of RMDN3 by PINK1 at T160 mediates the increased interaction between RMDN3 and VAPB during mitochondrial depolarization.

**Figure 6:**
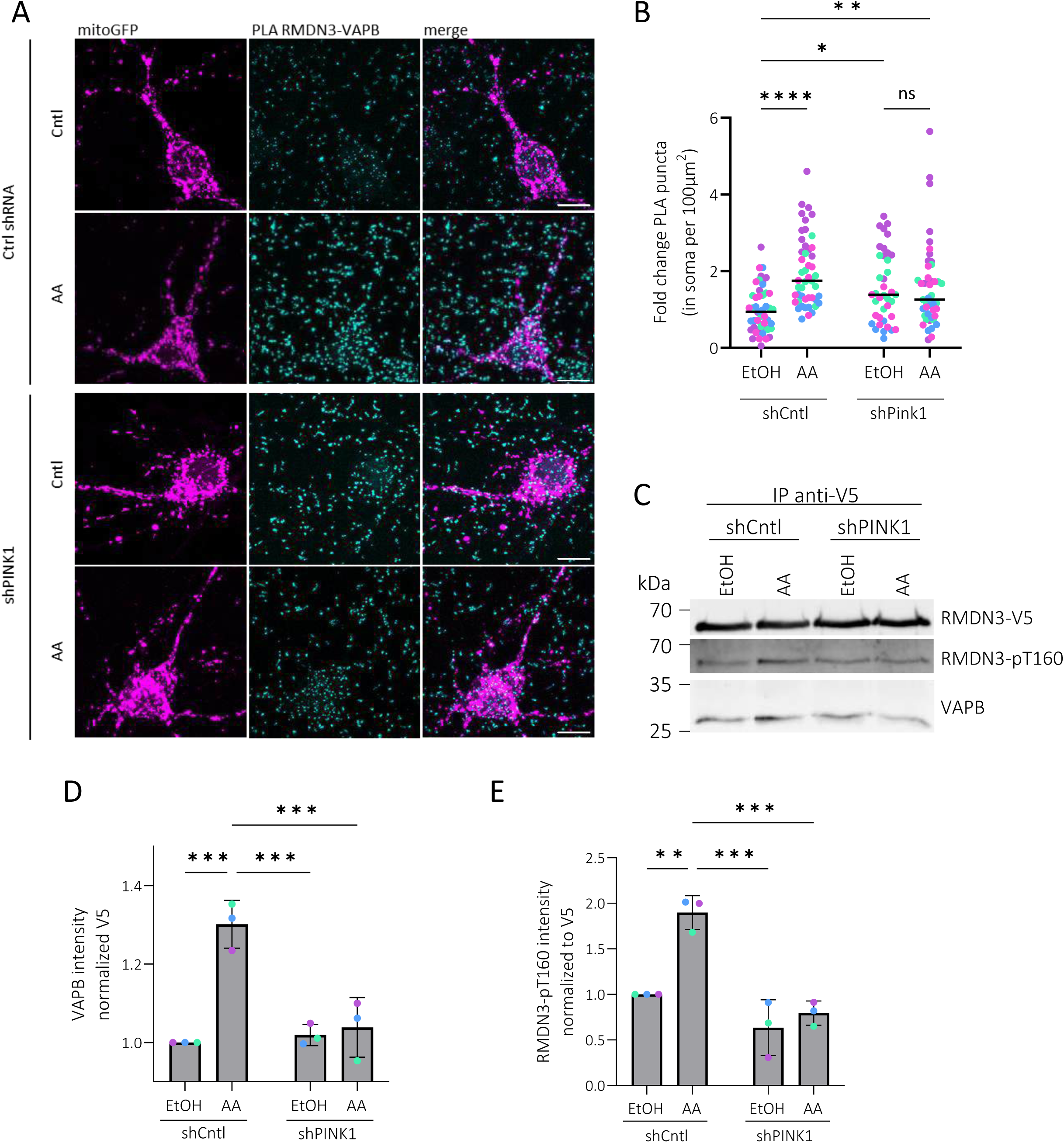
PINK1 is required for the damage-induced increase in RMDN3-VAPB interaction. **A)** Representative images of PLA analysis assessing RMDN3-VAPB proximity in control (shCntl) and PINK1 (shPINK1) shRNA expressing neurons under vehicle or AA treatment. MitoGFP-positive transfected cells are shown in magenta, PLA puncta corresponding to RMDN3-VAPB proximity are shown in cyan. The right column shows the merged signal. Scalebar is 10 µm. **B)** Quantification of PLA puncta from (A), normalized to the mean of the shCntl/vehicle condition of each experimental batch. Colors indicate independent experimental batches. Black lines indicate the mean with SEM. Statistical analysis was performed using two-way ANOVA. **C)** Co-IP analysis of VAPB binding to V5-tagged RMDN3 and assessing pT160 levels in N2A cells transfected with shCntl or shPINK1. **D, E)** Quantification of Co-IP shown in (C). Co-immunoprecipitated VAPB signal was normalized to the immunoprecipitated RMDN3-V5 of shCntl/vehicle condition (D). pT160 signal was normalized to RMDN3-V5 shCntl/vehicle condition. Statistical analysis was performed using two-way ANOVA.

## Discussion

During mitochondrial dysfunction, fundamental rearrangements of MERCS need to occur to allow the formation of the autophagosome around mitochondria. In cultured neurons, we observe a general increase in proximity between mitochondria and the ER in the soma upon induction of mitochondrial depolarization by AA treatment (Fig. 3E). However, the general increase in MERCS is not mirrored by all tether pairs alike. While the interaction between VAPB and RMDN3 increases as a response to mitochondrial depolarization in both, neurons and HEK293 cells, the contact site between RRBP1 and SYNJ2BP decreases in neurons (Fig. 3). The increased interaction between RMDN3 and VAPB depends on PINK1, a kinase activated during mitochondrial depolarization (Fig. 6A,B) which has previously been reported to localize to MERCS and promote contact between mitochondria and ER^22^. PINK1 directly phosphorylates RMDN3 in its FFAT motif (Fig. 5F), thereby enhancing its interaction with VAPB (Fig. 5B). Importantly, this stabilization will have differential consequences in early vs. late stage mitophagy. During early stages, the contact may ensure the continued delivery of the PINK1 precursor in a positive feedback loop to further stabilize the contact and amplify the PINK1 signal. This allows PINK1 to switch on repair mechanisms including the local translation of OXPHOS proteins^18^, as well as start the signaling cascade that will recruit the autophagic machinery to the damaged organelle. Together with the ubiquitin ligase Parkin, PINK1 activity initiates the phospho-ubiquitination of several outer membrane proteins, targeting them for proteasomal destruction. Among the OMM proteins degraded in a Parkin-dependent manner is also RMDN3^14,50^. Therefore, the increase in RMDN3-VAPB interaction mediated by PINK1 phosphorylation will be terminated by degradation of RMDN3, allowing the final engulfment of the damaged mitochondrion in an autophagosome. Excessive MERCS have been shown to suppresses Parkin-mediated mitophagy to proceed, including overexpression of RMDN3 or expression of synthetic tethers^51,52^. On the flip side, loss of RMDN3 and VAPB reduces autophagosome formation, which is tied to their role as MERCS proteins that enable the shuttling of Ca^2+^ ions from the ER to mitochondria^5^. Thus, the maintenance of the contact between RMDN3 and VAPB during early stages of mitophagy also supports the general upregulation of autophagy signaling necessary for later degradation of the damaged organelle.

During the early stage of mitophagy, loss of mitochondrial membrane potential will also elicit the generation of ROS^53^. It was recently shown that a contact between mitochondria and peroxisomes mediates the detoxification of ROS generated by mitochondria via the peroxisomal antioxidant enzymes^17^. Intriguingly, this contact is mediated by an interaction between RMDN3 and ACBD5, which we also recover in our MERCS splitAPEX proteome (Fig. 2A). The same peroxisomal protein, ACBD5, was previously also shown to interact with VAPB^29,54^. It is thus conceivable that the three proteins and thus the three organelles may form a tripartite contact site. Such a three-organelle interaction has been observed in the liver and is highest during metabolic shifts^55^. We did not observe a change in the association of peroxisomal proteins with RMDN3-VAPB MERCS during mitochondrial depolarization (Fig. 4B), suggesting that these contacts remain stable in their proteomic composition, but increase in their abundance. One caveat here is that the short treatment with H_2_O_2_ necessary to activate splitAPEX-mediated biotinylation may have already caused a baseline increase in oxidative stress, thus eliciting the formation of peroxisome-MERCS contacts already under basal conditions. While we cannot fully exclude movement of peroxisomes towards MERCS during this 1 min pulse with H_2_O_2_, it seems inconsistent with the reported timeline of over 30 min after rotenone intoxication^17^. In support of the physiology of our baseline conditions, we observe a clear trend towards the expected downregulation of Parkin substrates upon mitochondrial depolarization (Fig. 4D).

The significantly reduced proteins in RMDN3-VAPB proximity upon 1 h of AA/OA treatment (Fig. 4C) are not established Parkin substrates. Instead, collectively, the reduced proteins point towards selective remodeling and fine-tuning of RMDN3-VAPB MERCS during the early response to mitophagy rather than their disassembly, consistent with our PLA and FEMP data (Fig. 3). Among the reduced proteins TMX3, a reported regulator of MERCS^38^, and the ER Ca^2+^ pump SERCA which regulates ER Ca^2+^ loading^56^, point towards local remodeling of ER functions at these contact sites.

Given the established role of RMDN3 and VAPB in ER-mitochondria Ca^2+^ signaling^5,57^, it will be interesting to determine whether the reduced proximity of SERCA reflects changes in local Ca^2+^ handling during mitochondrial stress. Reduced proximity was also observed for several proteins linked to mitochondrial quality control, including the apoptotic regulator BAX^40^ as well as CPOX and SLC25A3, which have recently been implicated in Parkin recruitment and transient mitochondrial depolarization, respectively^58,59^. Whether the reduced recovery of these proteins reflect protein degradation, redistribution, or selective exclusion from remodeling MERCS remains to be determined.

Among the proteins increasing in proximity of RMDN3-VAPB contact sites, ACO2 and HSPA5 suggest that both mitochondria and ER stress responses are represented at these MERCS following mitochondrial depolarization. Interestingly, three additionally enriched proteins, ECHS1, DLST, and GCDH are all mitochondrial metabolic enzymes involved in distinct pathways of energy metabolism including fatty acid oxidation, the TCA cycle and amino acid catabolism^35–37^. Given the established role of RMDN3-VAPB contact in lipid metabolism and lipid transfer between mitochondria and the ER^60^, these findings raise the possibility that mitochondrial stress is accompanied by a metabolic reorganization at MERCS.

VAPB is found mutated in hereditary forms of amyotrophic lateral sclerosis (ALS)^61^. As this disease/mutant primarily affects neurons, we also sought to validate our findings in cultured neurons. Indeed, we observe a similar increase in RMDN3-VAPB contacts upon mitochondrial depolarization in cultured neurons (Fig. 3A,C), which could be prevented by PINK1 knock-down (Fig. 6A,B). Its partner protein RMDN3 exhibits an intriguing tissue specific expression in the brain^62^, particularly within the cerebellum^63^. Together, they influence key neuronal functions including synaptic activity^64^. Unfortunately, the low expression levels in transfected neurons prevented us from investigating the splitAPEX proteome also in neurons. Furthermore, ALS-associated changes in neuronal signaling disrupt the interaction between RMDN3 and VAPB leading to a reduced function of this MERC in ALS^65,66^. Our results now suggest that a reduction in interaction between RMDN3 and VAPB impacts the maintenance of functional MERCS during mitochondrial depolarization. Further research will elucidate how RMDN3-VAPB MERCS contribute to differential vulnerability of neurons to mitochondrial insults and how their function is affected in disease.

## Acknowledgments

We thank J. Lindner for technical support and all the members of the Harbauer and Behrends laboratory for their support and many fruitful discussions. We are grateful to E. Laurell/C. Polisseni/D. Paynter and M. Spitaler/M. Oster/G. Cardone from the Imaging facilities of the MPIs for Biological Intelligence (RRID:SCR_026797) and Biochemistry (RRID:SCR_025739), respectively, for assistance with cell imaging. For protein purification, we like to thank Y. Xiao und J. Basquin from the protein production facility. We also thank B. Steigenberger from the Mass Spec facility for help with phospho-proteomics. At the SyNergy Nanosclae hub, we would like to thank Martina Schifferer and Cornelia Förster for their help with the acquisition of EM images.

Work in ABH’s lab is supported by the Max Planck Society, DFG (DFG, HA 7728/6-1 ID 576693647, EXC 2145 SyNergy – ID 390857198, TRR353 – ID 471011418, SPP2453 ID 541742535), the European Union (ERC StG Project 101077138 — MitoPIP) and Germany’s Excellence Strategy within the framework of the Munich Cluster for Systems Neurology (EXC 2145 SyNergy – ID 390857198) and the Schram foundation (T0287/46550/2025). Views and opinions expressed are however those of the author(s) only and do not necessarily reflect those of the European Union or the European Research Council. Neither the European Union nor the granting authority can be held responsible for them. CB was supported by the Deutsche Forschungsgemeinschaft (DFG, German Research Foundation) within the frameworks of the Munich Cluster for Systems Neurology (EXC 2145 SyNergy, Project-ID 390857198), the Collaborative Research Center 1177 (ID 259130777) and the project grant BE 4685/11-2 (ID 447112704). Work in RF’s lab is supported by Germany’s Excellence Strategy within the framework of the Munich Cluster for Systems Neurology (EXC 2145 SyNergy – ID 390857198) Work in the lab of KW is funded by the German Research Foundation (SPP 2453, project number 541210481; RTG 2862, project number 492434978, and Germany’s Excellence Strategy - EXC 2033 - 390677874 – RESOLV) and the Michael J. Fox Foundation (Grant ID: 021968).

## Author Contributions

ABH and CB conceived the project and wrote the manuscript together with AR and AZ, with additional input from JD, MGH, KFW, RF and FM. AR and AZ designed and conducted most of the experiments. RF was responsible for generation of the phospho-specific antibody. FM was responsible for EM analysis. SK conducted phosphoproteomics analysis. MGH and KFW were responsible for the production of recombinant PINK1 kinase. AR, JD and CB were responsible for Mass Spectrometric analysis. All authors read and approved the manuscript.

## Methods

### Data availability

The datasets used during the study are available from the corresponding author on reasonable request. The MS data have been deposited to the ProteomeXchange Consortium (http://proteomecentral. proteomexchange.org) via the PRIDE partner repository with the dataset identifiers PXD080602 (splitAPEX) and PXD081057 (Phospho-proteomics).

### Animal procedures

Hippocampal neurons were isolated from E16.5 mouse embryos as previously described in Hees et al.^21^ All procedures involving mice in this study received approval from the Government of Upper Bavaria and were conducted observing all relevant guidelines and regulations. C57BL/6 WT mice were bred in the animal facility of the Max Planck Institute for Biological Intelligence, Martinsried, Germany. The animals were housed in a controlled environment including unrestricted access to food and water. Experiments involved embryos of both sexes.

Female Wistar rats aged 4 to 7 months were used for immunization in accordance with the German Animal Welfare Law and with the approval of the local authorities of Upper Bavaria, Germany (ROB-55.2Vet-2532.Vet_03-22-25).

### Neuronal cultures

In short, isolated hippocampi were collected in ice-cold dissociation medium (Ca^2+^-free Hank’s Balanced Salt Solution (Thermo Fisher Scientific) containing 10 mM MgCl2, 1 mM kynurenic acid (Sigma-Aldrich), and 10 mM HEPES). Hippocampal tissue was dissociated with papain/L-cystein (50 units, Roche) for 5 min at 37 °C. Subsequently, trypsin inhibitor (Abnova) was added and the tissue was gently triturated by repeated pipetting. Primary neurons were cultured in Neurobasal medium (Gibco) supplemented with B27 (2%, Gibco) and penicillin/ streptomycin/ glutamin (1%, Gibco).

Neurons were seeded at a density of 1*10^5^ on acid-washed coverslips coated with poly-L-lysine (Sigma-Aldrich) and laminin (ThermoFisher Scientific) in 24-well plates for immunofluorescence staining and PLA analyses. For electron microscopy, primary neurons were seeded at a density of 2*10^5^ cells on aclar in 24-well plates. Cells were transfected on DIV5 with Lipofectamine 2000 (Thermo Fisher Scientific). Unless differently specified, cells were treated and fixed on DIV8, PINK1-knockdown experiments were conducted at DIV9.

### Cell line cell culture

Human Embryonic Kidney (HEK) 293T cells and Neuro-2A (N2A) cells were maintained in DMEM (Thermo Fisher Scientific) supplemented with 10% FBS, penicillin and streptomycin (1%, Gibco). Cells were seeded at a density of 6*10^6^ cells in a 10 cm dish for Western blot and CoIP analyses. HEK293T cells were transfected with Calcium-Phosphate, N2A cells were transfected on DIV1 for 1 hour using lipofectamine 2000 transfection reagent (Thermo Fisher Scientific) in Opti-MEM GlutaMAX reduced serum medium (Gibco). HEK293T and N2A cells were kept in culture for 48 h past transfection. For electron microscopy, HEK293 cells were seeded at a density of 5*10^4^ on aclar, transfected the next day and allowed 48 h for gene expression.DNA constructs.

For the splitAPEX constructs, RMDN3 was C-terminally fused to GFP and to V5-AP (FKBP-V5-AP-nes_pLX304 was a gift from Alice Ting (Addgene plasmid # 120912; http://n2t.net/addgene:120912; RRID:Addgene_120912)) into the pLJM1 backbone. EX-HA (EX-HA-FRB-Cb5_pLX304 was a gift from Alice Ting (Addgene plasmid # 120915; http://n2t.net/addgene:120915; RRID:Addgene_120915)) was coupled to mCherry fused N-terminally to VAPB. For knockdown experiments, shControl (Sigma, SHC016) and shPINK1 (Sigma, TRCN0000026743) were used. FEMP was a gift from Luca Scorrano (Addgene plasmid # 191973; http://n2t.net/addgene:191973; RRID:Addgene_191973). For recombinant RMDN3, the cytosolic domain of RMDN3 was cloned into pET29b, creating a C-terminally 6xHis tagged protein (pET29b-dTM-RMDN3-6His).

### Immunofluorescence staining

Cells were fixed for 15 mins with 4% PFA pre-warmed to 37 °C. Subsequently, cells were permeabilized with 0.3% Triton x-100 in PBS for 12 min and blocked with 3% BSA in PBS for 1h at room temperature. Primary antibody staining was conducted at 4 °C overnight. Primary antibodies were used at the following dilutions in 3% BSA in PBS: anti-HA: Cell signaling, cat. no. 3724S, 1:500; anti-V5: Invitrogen, cat.no. R960-25, 1:1000, anti-biotin: Invitrogen, cat. no. 31852, 1:1000).

Secondary antibodies coupled to Alexa fluorophores were purchased from Invitrogen and used 1:500.

### Transmission electron microscopy imaging

Cells expressing split APEX RMDN3-GFP-V5-AP (RMDN3-AP) and EX-HA-mCherry-VAPB (EX-VAPB) fusion constructs were grown on aclar sheets (Science Services). Neurons were treated with 40 µM AA for 30 min before fixation. HEK293 cells were treated with 10 µM OA and 40 µM AA for 1 h. Cells were fixed in 2.5% glutaraldehyde (EM-grade, Science Services) in 0.1 M sodium cacodylate buffer (pH 7.4) for 30 min on ice. After washes in buffer, endogenous peroxidases were blocked in 20 mM glycine (Sigma) in buffer for 5 min and cells washed in buffer. The 1x diaminobenzidine (DAB) solution was prepared in buffer with 2 mM calcium chloride from a 10x DAB stock (Sigma) in hydrochloric acid (Sigma) and added to the cells for 5 min without and for another 40 min with 10 mM H_2_O_2_ (Sigma). After washes in buffer, cells were postfixed in osmium (0.5% osmium tetroxide, Science Services) for 5 min, washed in buffer and water and incubated for 5 min in 0.5% aqueous uranyl acetate (Science Services). Dehydration was accomplished using a graded series of ice-cold ethanol. Cell monolayers were infiltrated in LX112 (Ladd research) and cured for 48 h at 60 °C. Cells were ultrathin sectioned at 50 nm onto formvar-coated copper grids (Plano). Transmission electron micrographs were acquired on a JEM 1400plus (JEOL) equipped with a XF416 camera (TVIPS) and the EM-Menu software (TVIPS). Images were processed in ImageJ.

### Proximity Ligation Assay (PLA)

After permeabilization with 0.3% triton-x for 12 min, PLA in farRed (DUO92013) was conducted according to the manufacturer’s instructions. Cells were incubated overnight at 4 °C with primary antibodies RMDN3/PTPIP51 (Bioss, bs-5719R, 1:100) and VAPB (Proteintech 66191-1-Ig, 1:50), or RRBP1 (abcam, ab95983, 1:600) and SYNJ2BP (Sigma, SAB1400613, 1:50). On the following day, cells were incubated with the PLA probes Anti-Mouse PLUS (DUO92001) and anti-Rabbit MINUS (DUO92005).

For endogenous PLA staining of RRBP1-SYNJ2BP and VAPB-RMDN3, mitochondria and neurites were additionally visualized with an immunofluorescence staining. All following steps described were conducted in the dark. Following the completion of the PLA protocol, cells were carefully fixed with 4% PFA prewarmed to 37 °C and washed 3x with PBS. Subsequently, cells were incubated with LETM1 (Invitrogen, cat. no. PA5-22233, 1:500) and Smi31 (Biolegend, cat. no. 801602, 1:250) in PLA antibody diluent overnight at 4 °C. On the following day, cells were washed 3x with PBS, and incubated with species-specific secondary antibodies (Invitrogen, 1:500) for 1.5 h at room temperature and washed off with 2x PLA wash buffer B. Lastly, coverslips were briefly rinsed in 1:100 diluted PLA wash buffer B and mounted for imaging.

### SplitAPEX biotinylation

For biotinylation, HEK293 cells were incubated with either vehicle control (EtOH and DMSO) without BP (-BP), vehicle control and 500 µM final concentration of BP (+BP) or 500 µM BP together with OA (10 µM), and AA (40 µM) (+BP +OA/AA) for 1 h. -BP samples were lysed directly, +BP and +OA/AA +BP samples were treated with 1 mM final H_2_O_2_ concentration in the conditioned media. Afterwards cells were incubated 2x 1min with quenching solution (10 mM sodium ascorbate, 5 mM TROLOX, 1 mM sodium azide in PBS) and afterwards washed 2x with PBS. Subsequently, cells were lysed as described in Western blot or Mass spectrometry analyses section.

### Western blot analysis

Lysates were prepared in RIPA buffer (Serva) supplemented with PhosSTOP (Roche), protease inhibitor (Roche) and benzonase (Invitrogen, 1:2000). Then 4x Laemmli was added to a final concentration of 1x and samples were boiled for 5 min. Proteins were separated in a 12% SDS-PAGE gel, and transferred for 90 min on a nitrocellulose membrane. To assess protein loading, membranes were stained with Pierce reversible stain (Thermo Fisher Scientific) for 5 min, washed with the destain-solution for 5 minutes, imaged at the iBright FL1000 Imaging System (Thermo Fisher Scientific), and subsequently destained with stain eraser-solution for 10 min. Membranes were rinsed with distilled water and blocked with 5% nonfat milk powder dissolved in Tris buffered saline containing 0.1% Tween-20 (TBS-T). Subsequently membranes were incubated in streptavidin or adequate antibody solutions overnight, shaking at 4 °C. Next day, membranes were washed 2x 5 min with TBS, incubated with secondary antibody coupled to ALEXA-plus fluorophores (Thermo Fisher Scientific, diluted 1:5000) for 1-1.5 h in 5% milk-TBS-T at room temperature shaking in the dark. Fluorescence signals were acquired using the iBright FL1000 Imaging System. Image processing and densitometry analyses were conducted in FIJI/ImageJ^67^. Primary antibodies were used at the following dilutions in 5% milk in TBS-T: HA (Cell signaling, cat. no. 3724S, 1:100), V5 (Invitrogen, cat. no. R960-25, 1:5000), β-III-tubulin (2G10) (Invitrogen, cat. no. MA1-118, 1:2000), β-actin (AC-74) (Sigma-Aldrich, cat. no. A5316, 1:1000), VAPB (Invitrogen, cat. no. PA5-53023, 0.4 µg/mL). Phospho-ubiquitin (Ser65) (Cell signaling, cat. no. 62802, 1:500) was diluted in 3% BSA TBS-T. Streptavidin was conjugated to DyLight488 (abcam, cat. no. ab134349, 1:1000) or Alexa Fluor647 (Invitrogen, cat. no. S21374, diluted 1 µg/mL) and diluted in 3% BSA in TBS-T.

For the phospho-specific antibody targeting T160 of RMDN3, a phosphorylated peptide corresponding to RMDN3 T160 (SSVYF**pT**ASSGA, aa155-165, Peps4LS) coupled to ovalbumin was used to immunize female Wistar rats with ovalbumin-coupled peptides (VFCSNSQASQPC; aa 1238-1249;). Boost injections were given five and thirteen weeks later, and spleen cells were fused with myeloma cell line P3 3 63-Ag8.653 (ATCC, American Type Culture Collection) by standard procedures. Hybridoma supernatants were screened in a flow cytometry assay (iQue, Intellicyt; Sartorius) for binding to biotinylated peptides coupled to streptavidin beads (PolyAN, Berlin) and negatively selected against the non-phosphorylated peptide. Positive clones were further validated in immunoblotting. Clone RMDP 23C11 (IgG2b/λ) was subcloned twice by limiting dilution to obtain a stable antibody-producing monoclonal cell line (RRID: AB_3751972).

### Streptavidin-Pulldown for mass spectrometry

Biotinylation for splitAPEX experiments was conducted as described in SplitAPEX biotinylation. Following quenching, cells were washed 3x with PBS, scraped off, washed twice with PBS in suspension, and lysed on ice in freshly prepared RIPA buffer supplemented with PhosSTOP and Protease inhibitor. Protein concentration was determined by BCA and input was adjusted accordingly. Tryptic in-solution digestion of enriched biotinylated fractions was performed as described by Lobingier et al^68^, followed by LC-MS-analysis. In brief, cleared cell lysates were incubated over night with pre-equilibrated streptavidin-agarose beads. Subsequently, the beads were washed twice with RIPA buffer and three times with freshly prepared 3 M Urea buffer (in 50 mM ammonium bicarbonate). Protein samples were reduced by adding TCEP (5 mM final; Sigma) while shaking at 55°C for 30 min. Samples were alkylated with iodoacetamide (10 mM final; Sigma) covered from light at room temperature for 20 min and quenched with DTT (20 mM final; Sigma). Samples were washed twice with 2 M Urea (in 50 mM ammonium bicarbonate) before overnight trypsin digestion with 1 µg trypsin (Promega) per sample at 37 °C. Peptides were recovered by supernatant collection, before washing the resin twice with 2 M Urea (in 50 mM ammonium bicarbonate) and pooling the fractions. Samples were acidified with 1% trifluoroacetic acid and concentrated by vacuum centrifugation. For mass spectrometric analysis, peptide samples were desalted on custom-made C18 stage tips^69^, and solved in 0.1% formic acid.

### MS data collection and analysis for proximity proteomics

Digested peptide mixtures were separated using an Easy nLC1200 liquid chromatograph (Thermo Scientific) followed by peptide detection on a Q Exactive HF mass spectrometer (Thermo Scientific). Samples separation was performed on a 75 µm × 15 cm custom-made fused silica capillary packed with C18AQ resin (Reprosil-PUR 120, 1.9 µm, Dr. Maisch), with a 35 min acetonitrile gradient in 0.1% formic acid at a flow rate of 400 nl/min (5–38% ACN gradient for 23 min, 38–60% ACN gradient for 3 min, 60–95% ACN gradient for 2 min). Peptides were ionized using a Nanospray Flex Ion Source (Thermo Scientific). Peptides were identified in full MS / dd MS² (Top15) mode, dynamic exclusion was enabled for 20 s and identifications with an unassigned charge or charges of one or >8 were rejected. MS1 resolution was set to 60,000 with a scan range of 300–1650 m/z, MS2 resolution to 15,000 and the AGC target1 was set to 3e6.

Raw mass spectrometric data were analyzed using MaxQuant’s (version v2.6.7.0)^70^ Andromeda search engine in reversed decoy mode based on human reference proteome (Uniprot-FASTA, UP000005640) with a false discovery rate (FDR) of 0.01 at both peptide and protein levels. Digestion parameters were set to specific digestion with trypsin with a maximum number of 2 missed cleavage sites and a minimum peptide length of 7. Oxidation of methionine and amino-terminal acetylation were set as variable and carbamidomethylation of cysteine as fixed modifications. The tolerance window was set to 20 ppm (first search) and to 4.5 ppm (main search). Label-free quantification (LFQ) was set to a minimum ratio count of 2, re-quantification and match-between-runs was selected and 4 biological replicates per condition were analyzed. The resulting protein group files of MaxQuant analyzed using Perseus (version 2.0.7.0)^71^ as described in Wu et al^72^. Briefly, common contaminants and reverse hits were excluded prior to downstream analyses. LFQ values were used for statistical analyses and differentially enriched proteins were identified using a two-sample Student’s t-test followed by FDR correction, with an FDR threshold of 0.05. DAVID^73,74^ was used for functional enrichment analyses of cellular component (CC).

For the annotation of previously reported Parkin ubiquitination targets, OMM proteins identified after 2 h of treatment with CCCP were extracted from Zittlau et al.^14^ Proteins with WT-Parkin_2h values > 0.5 and C431A-Parkin_2h values < 0.5 were considered Parkin-depnendent targets.

### Fluorescence lifetime imaging

To quantify Förster resonance energy transfer (FRET) of the FEMP biosensor, fluorescence lifetime imaging microscopy (FLIM) was used. Primary neuronal cultures were transfected on DIV5 with FEMP, and were treated and fixed on DIV7. Imaging was performed using the Leica SP8 Falcon confocal microscope equipped with an HCX PL APO 63x/1.2 motCORR CS water-immersion objective and LAS X software (version 3.5.7.23225) with the FLIM module (Leica) at 22 °C at the Imaging Facility of the Max Planck Institute for Biochemistry in Martinsried, Germany. The donor fluorophore was excited with a pulsed Sepia Picosecond Dioide Laser with 440 nm laser operating at a repetition rate of 20 MHz (PicoQuant). Donor fluorescence lifetimes were determined by phasor analyses as previously described^75^. For each cell, the center of the phasor distribution for the donor lifetime was determined from an ROI encompassing the soma, using 3×3 pixel binning.

### Protein expression and purification of RMDN3

Sequence verified pCoofy49 expression plasmids encoding RMDN3 WT or T160A were transformed by heat shock into Ca2+-competent E. coli Rosetta 1 (DE3) cells and plated on LB agar supplemented with kanamycin and chloramphenicol for selection. Single colonies were used to inoculate 50 ml overnight LB cultures. Fresh TB medium was inoculated with overnight cultures at an initial OD600 of 0.04. Cells were grown at 37 °C to an optical density at 600 nm (OD600) of approximately 0.6–0.8, and protein expression was induced by addition of 0.1 mM isopropyl β-D-1-thiogalactopyranoside (IPTG). After 24 h of expression at 18 °C, cells were harvested by centrifugation at 5,000 × g for 10 min at 4 °C, yielding cell pellets of 9 g (WT) and 40 g (T160A). Cells were grown to an optical density at 600 nm (OD600) of approximately 0.6–0.8. Protein expression was induced by addition of 0.1 mM isopropyl β-d-1-thiogalactopyranoside (IPTG). After 24 h of expression, cells were harvested by centrifugation at 5,000 × g for 10 min at 4 °C, yielding cell pellets of 9 g (WT) and 40 g (T160A).

For the purification of RMDN3 variants, bacterial pellets were resuspended in lysis buffer (either 20 mM Tris pH 8.0/500 mM NaCl/10% glycerol for WT or 50 mM sodium phosphate pH 7.5/250 mM NaCl/30 mM imidazole for T160A) supplemented with 1 mM AEBSF-HCl, 2 µg/ml aprotinin, 1 µg/ml leupeptin, 1 µg/ml pepstatin, 2.4 U/ml Benzonase, and 2 mM MgCl₂, then lysed via sonication and clarified by centrifugation at 62,000g for 30 min at 4 °C. The WT supernatant was purified using Strep-Tactin affinity chromatography and eluted with biotin, while the T160A variant was purified via Ni-NTA affinity chromatography and eluted with 800 mM imidazole. Pooled elution fractions for both proteins were incubated overnight with His-tagged HRV 3C protease for tag cleavage, subjected to a subtractive Ni-NTA step to remove the protease and uncleaved protein, and finally polished using size-exclusion chromatography on a HiLoad 16/60 Superdex 75 column equilibrated in storage buffer (20 mM Tris pH 7.2, 30 mM NaCl, 10% glycerol). Purified peak fractions were pooled, aliquoted, snap-frozen in liquid nitrogen, and stored at −80 °C.

### Recombinant PINK1 kinase preparation

Competent *E. coli* Rosetta 1 (DE3) were transformed with either wildtype pGEX-PINK1 WT or catalytically inactive pGEX-PINK1(D359A) by heat shock as described previously (PMID: 36398858). In brief, single colonies were used to inoculate 100 ml overnight LB cultures. LB medium was inoculated and grown at 37°C to an optical density at 600 nm (OD600) of approximately 0.6–0.8.

Protein expression was induced by the addition of 0.2 mM isopropyl β-D-1-thiogalactopyranoside (IPTG). After 24 h of expression at 18°C, cells were harvested by centrifugation at 4,000 × g for 20 min at 4°C.

For the purification, bacterial pellets were resuspended in lysis buffer 50 mM Tris-HCl, pH 8.0, 150 mM NaCl, 5% glycerol, 5mM DTT, 1mM MgCl_2_ supplemented with protease inhibitors, DNase, and RNase. The cells were lysed by French Press and clarified by centrifugation at 12000 × g for 30 min at 4°C. The supernatant was collected, and the proteins were purified using a GSTPrep™ FF 16/10 column (Cytiva) on an Äkta Pure 25. After the lysate was passed through the column, the column was washed with the lysis buffer. The elution was performed using the lysis buffer supplemented with 20 mM glutathione. The eluted fraction was dialyzed overnight in 50 mM Tris-HCl, 100 mM NaCl, 0.5mM TCEP. Finally, the protein was concentrated, centrifuged, and applied to size-exclusion chromatography using a HiLoad 16/60 Superdex 75 column equilibrated in 50 mM Tris-HCl, 100 mM NaCl, 0.5 mM TCEP. Purified peak fractions were analyzed by SDS-PAGE. The pure fractions were pooled, aliquoted, flash-frozen in liquid nitrogen, and stored at −80 °C.

### *In vitro* phosphorylation assay

Prior to phosphorylation, the recombinant cytosolic domain of RMDN3 (WT or T160A, 1 µg) was dephosphorylated 1 µL of calf intestinal phosphatase (NEB) in 1 x CutSmart buffer (NEB) and a total volume of 10 µL. The mixtures were incubated at 37 °C for 1 h, followed by the addition of 1 x phosphatase inhibitor (PhosStop, Roche) to terminate the reaction. The dephosphorylated recombinant RMDN3 (1 µg in 13 µL) was then subjected to in vitro phosphorylation with active recombinant *T. castaneum* PINK1 (1.3 µM), the kinase inactive mutant PINK1(D359A) (1.3 µM) or active recombinant AMPK (3.1 x 10^-12^ µM) in kinase reaction buffer (8 mM MOPS/NaOH, pH 7 and 200 µM EDTA) with 200 µM ATP (Serva) in 30 µL total volume. For the RMDN3 only control condition addition of kinase was omitted. The samples were incubated at 30 °C for 2 h (shaking, 300rpm). To terminate the phosphorylation, Laemmli sample buffer was added and samples were boiled at 95 °C for 5 min.

### Co-Immunoprecipitation

Lysates were prepared in RIPA buffer supplemented with protease inhibitor (Roche), phosphatase inhibitor (PhosStop, Roche), 200 nM PMSF, and Benzonase (1:2000). 6% of lysates were used as input samples, added with 4x Laemmli, boiled for 5 minutes and stored at −20 °C. The remains of the lysates were diluted 1:4 with IP-wash buffer (0.1% Triton X-100, 20 mM Tris, pH 8, 140 mM KCl, 5 mM MgCl_2_, 0.5 mM DTT, 200 nM PMSF) and incubated with V5-antibody (1:2000) for 1 h with overhead rotation at 4 °C. Subsequently, 100 µL of Protein A-Sepharose 4B Conjugate Beads (Thermo Fisher Scientific, 10-1141) were added and the solution was rotated for 4 h at 4 °C. Beads were washed 4x with IP wash buffer and subsequently boiled at 95 °C with 50µL of 1x Laemmli. The whole eluate was probed on Western blot.

N2A cells were treated for 30 min with 40 µM AA or vehicle control (ethanol) and were lysed in SDS-free IP lysis buffer (25 mM Tris/HCl, pH 7.4, 150 mM NaCl, 1% NP-40 and 1 mM EDTA freshly supplemented with 200 nM PMSF, protease inhibitor (Roche) and PhosSTOP (Roche). The following procedure was performed as described above with minor modifications. Wash buffer contained 150 mM NaCl instead of 140 mM KCl, lysates were incubated with V5-antibody for 2 h at 4 °C under rotation instead of 1h. Finally, proteins were eluted by boiling with 1x Laemmli for 4 min at 95 °C.

### Sample preparation for LC-MS/MS (phosphoproteomics)

Samples from recombinant protein assay (without addition of Laemmli sample buffer) were supplemented with SDC buffer containing 1% sodium deoxycholate (SDC; Sigma-Aldrich), 40 mM 2-chloroacetamide (CAA; Sigma-Aldrich), 10 mM tris(2-carboxyethyl)phosphine (TCEP; Thermo Fisher Scientific), and 100 mM Tris (pH 8.0) in a 1:1 ratio and cooked at 95 °C for 10 min.

Subsequently, samples were sonicated using a Bioruptor Plus (Diagenode) for 10 cycles of 30 s at high intensity, followed by another incubation at 37 °C shaking for 30 min. Protein digestion was performed overnight at 25 °C with rChymoselect (Promega) at 1:20 enzyme to substrate ratio. The resulting peptide mixtures were acidified to a final concentration of 1% trifluoroacetic acid (TFA; Merck) and centrifuged to precipitate SDC. Peptides were purified with SCX-plugged tips and were loaded on Evosep Pure tips (Evosep).

### LC-MS/MS (phosphoproteomics)

LC-MS analysis was conducted on an Evosep ENO HPLC system (Evosep) coupled to a SCIEX ZenoTOF 7600 equipped with the SCIEX OptiFlow 1-50 µL ion source (SCIEX). Peptides were eluted from Evosep Pure tips onto a C18 reversed phase PepSep column (8 cm x 150 µm x 1.5 µm) (Bruker Daltonics) and separated with a 60 samples per day (SPD) method of 22.4 min duration. Mobile phase consisted of buffer A (H₂O, 0.1% FA) and buffer B (99.9% ACN, 0.1% FA). The column compartment was maintained at 50 °C throughout the measurement and the multi sampler temperature was kept at 4 °C.

Ion source and gas parameters were set as follows: Ion source gas 1: 20 psi; ion source gas 2: 60 psi; curtain gas: 35 psi; CAD gas 7; and temperature: 250 °C. The spray voltage was set to 4500 V, with a 100 ms TOF MS accumulation time, a mass scan range set to 300–2000 m/z, a declustering potential of 80 V, and a collision energy of 10 V.

Data were acquired in positive ion mode in data-dependent acquisition (DDA). Max. 20 precursor ions (Top20) exceeding the intensity threshold of 100 cps and of charge state ranging from 2-8 were selected for fragmentation with 10 s dynamic exclusion after one occurrence.

As fragmentation method electron activated dissociation (EAD) was chosen with the following parameters: collision energy: 12 V; electron kinetic energy (KE): 7 eV, electron beam current: 6500 nA, reaction time: 10 ms; TOF MS/MS mass scan range: 100–3000 m/z and accumulation time: 25 ms. ETC was set to dynamic and Zeno pulsing was enabled with a 100,000-cps threshold.

### Data analysis (phosphoproteomics)

Raw data were analyzed with the PEAKS Studio software (version 13.1) using a customized databank. Precursor mass error tolerance was set to 10.00 ppm, fragment mass error tolerance to 0.02 Da, max. missed cleavage to 2 and peptide length range to 6-45. Carbamidomethylation (+57.02) was added as fixed modification and acetylation (N-term) (+42.01), oxidation (M) (+15.99) and phosphorylation (STY) (+79.97) as variable modifications. PSM FDR and protein group FDR were kept at 1.0% and min. of unique peptides per protein was set to ≥ 1. PTM site localization was considered reliable for an AScore ≥ 15 and a relative ion intensity of site-determining fragment ions of ≥ 1%.

### STRING network analysis

For STRING network visualization, proteins identified in the RMDN3-VAPB splitAPEX proteome were assigned to primary cellular compartments based on UniProt subcellular location annotations.

Proteins were filtered for ER-, mitochondria- and peroxisome-associated annotations. Up to 25 proteins were selected for each compartment, based on enrichment in the splitAPEX proteome and ranked by log₂ fold change. Protein association networks were retrieved from the STRING database using Homo sapiens as reference organism, functional network mode and no additional interaction score cutoff at the query step. For visualization, STRING associations were filtered to retain only edges supported by experimental or curated database annotations. Textmining-only, coexpression, neighborhood, fusion, and co-occurrence associations were excluded. Edge width is proportional to the STRING evidence score. The RMDN3-ACBD5 interaction was manually added based on literature support^17^ and displayed as a dashed edge.

### Quantification and statistical analyses

All statistical analyses were conducted with GraphPad Prism version 10.6.0 (890). Comparisons of two groups were analyzed using an unpaired two-tailed Student’s t-test. Experiments involving two independent variables were performed using a two-way ANOVA followed by a post hoc test.

Statistical significance was defined as p < 0.05 (*), p < 0.01 (**), p < 0.001 (***), p < 0.0001 (****). In the graphs, data points in different colors represent independent biological replicates.

---

Supplementary table 1

S1A) Proteomic dataset obtained from RMDN3-VAPB splitAPEX proximity labeling in the presence or absence of BP. LFQ intensities for four biological replicates per condition are shown together with protein annotations, peptide and sequence coverage information and statistical analysis comparing +BP to -BP. Significant enrichment was determined by a permutation-based Student’s t-test (FDR < 0.05). S1B) Comparison of all proteins identified by RMDN3-VAPB splitAPEX and BiFCPL^28^ as well as ER-associated and mitochondria-associated proteins separately.

Supplementary table 2

Proteomic dataset obtained from RMDN3-VAPB splitAPEX proximity labeling following treatment with AA/OA or vehicle in the presence of BP. LFQ intensities for four biological replicates per condition are shown together with protein annotations, peptide and sequence coverage information and statistical analysis comparing +BP +AA/OA to +BP and vehicle. Significant enrichment was determined by a permutation-based Student’s t-test (FDR < 0.05).

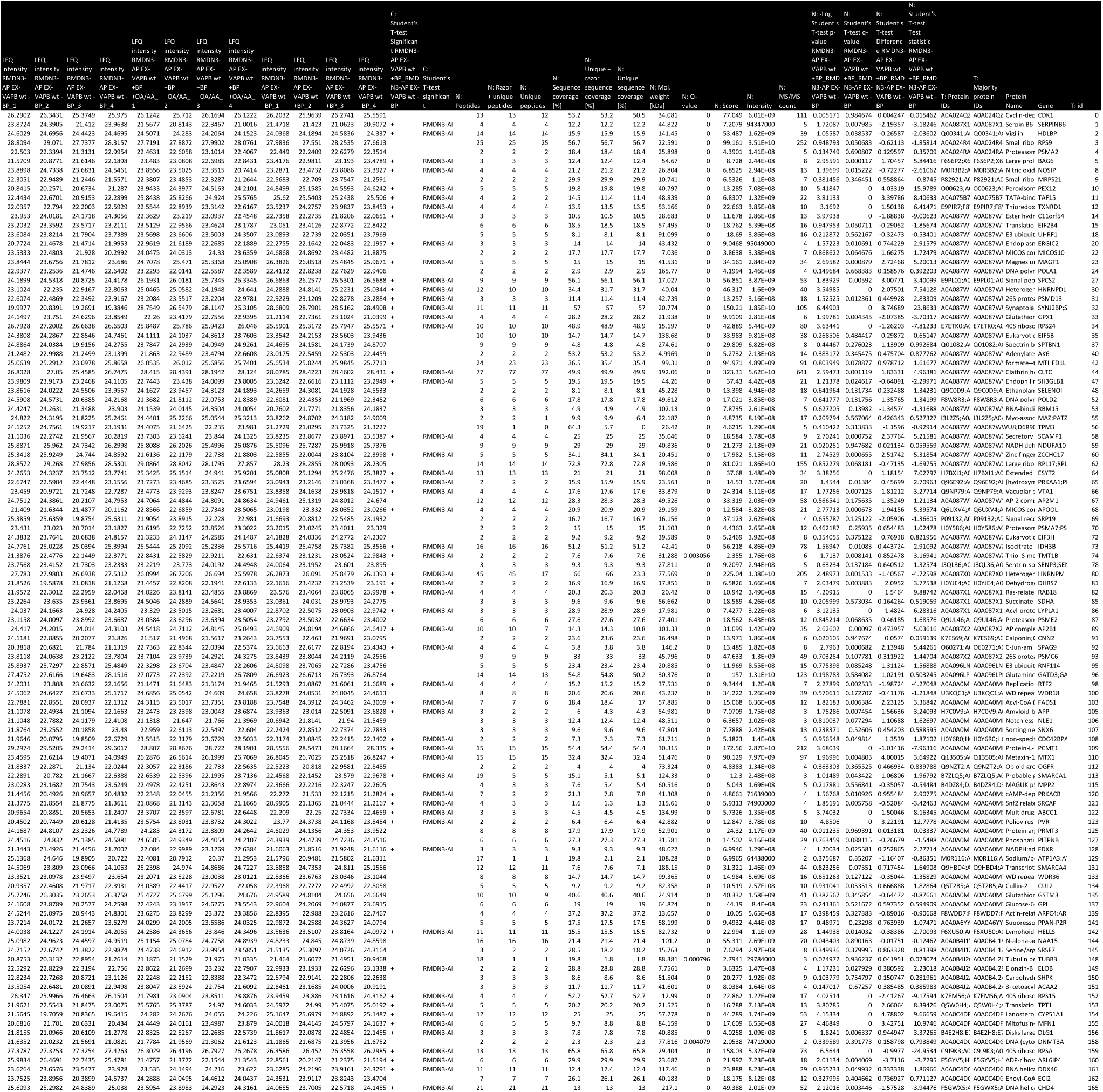
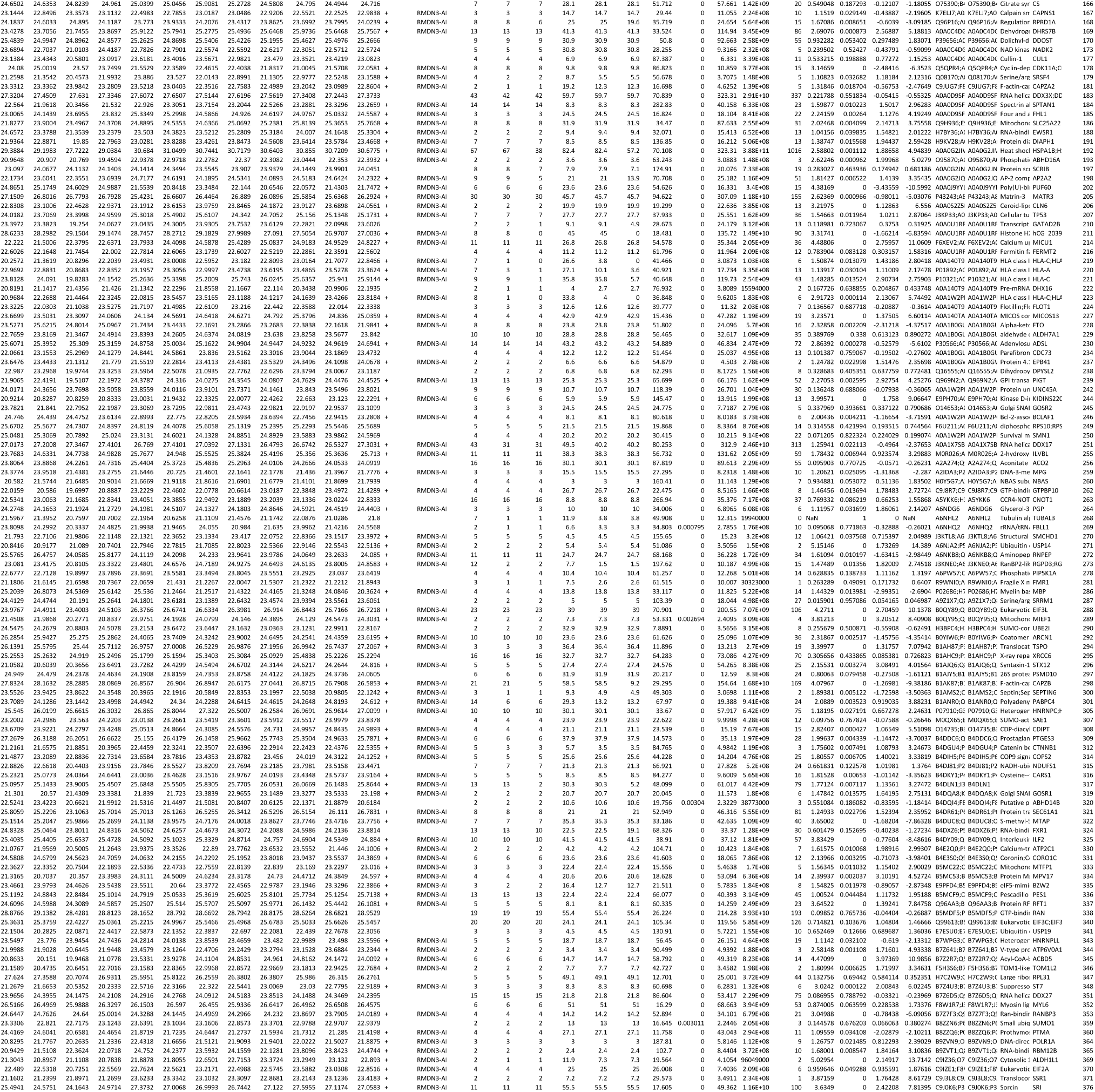
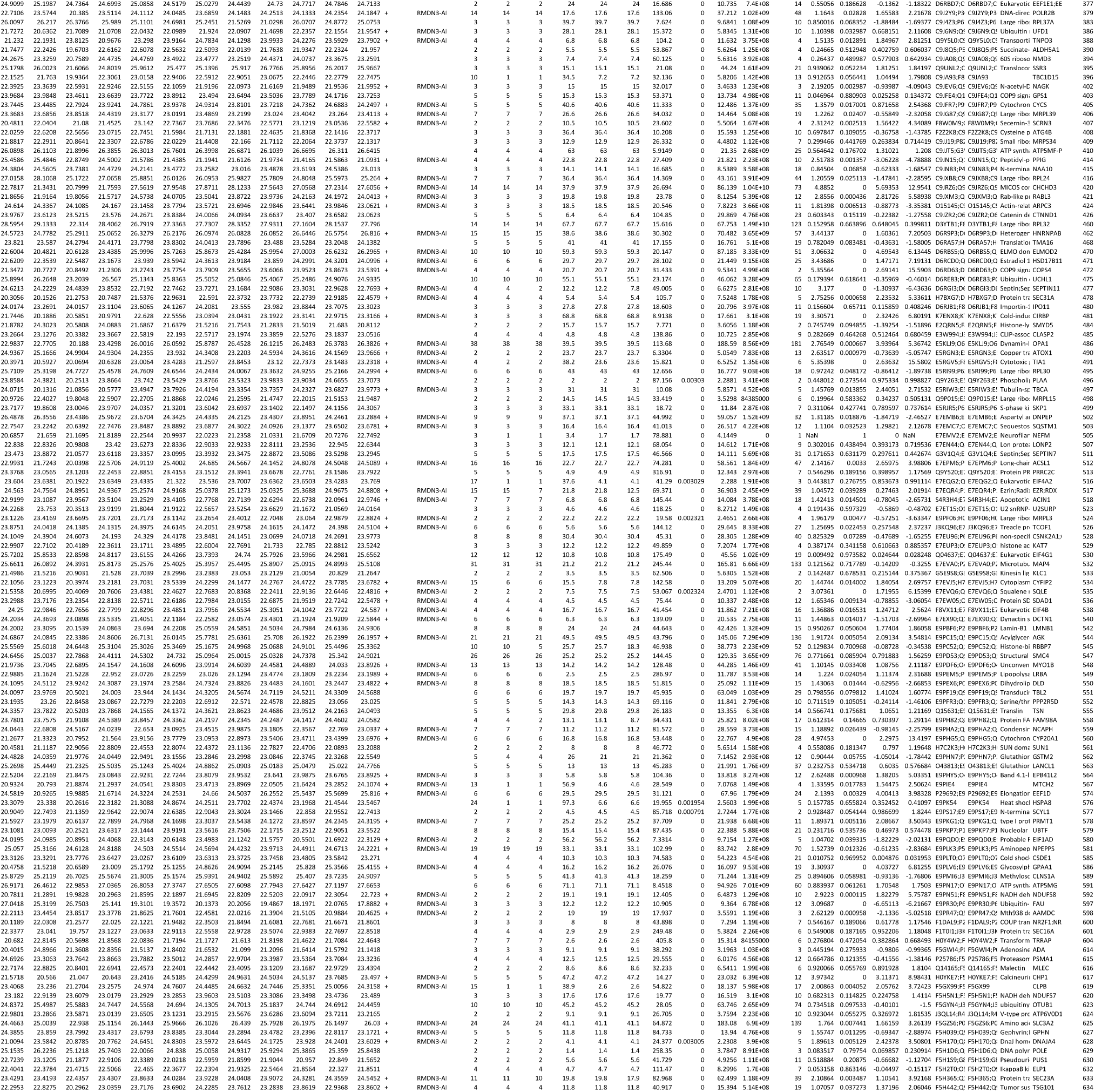
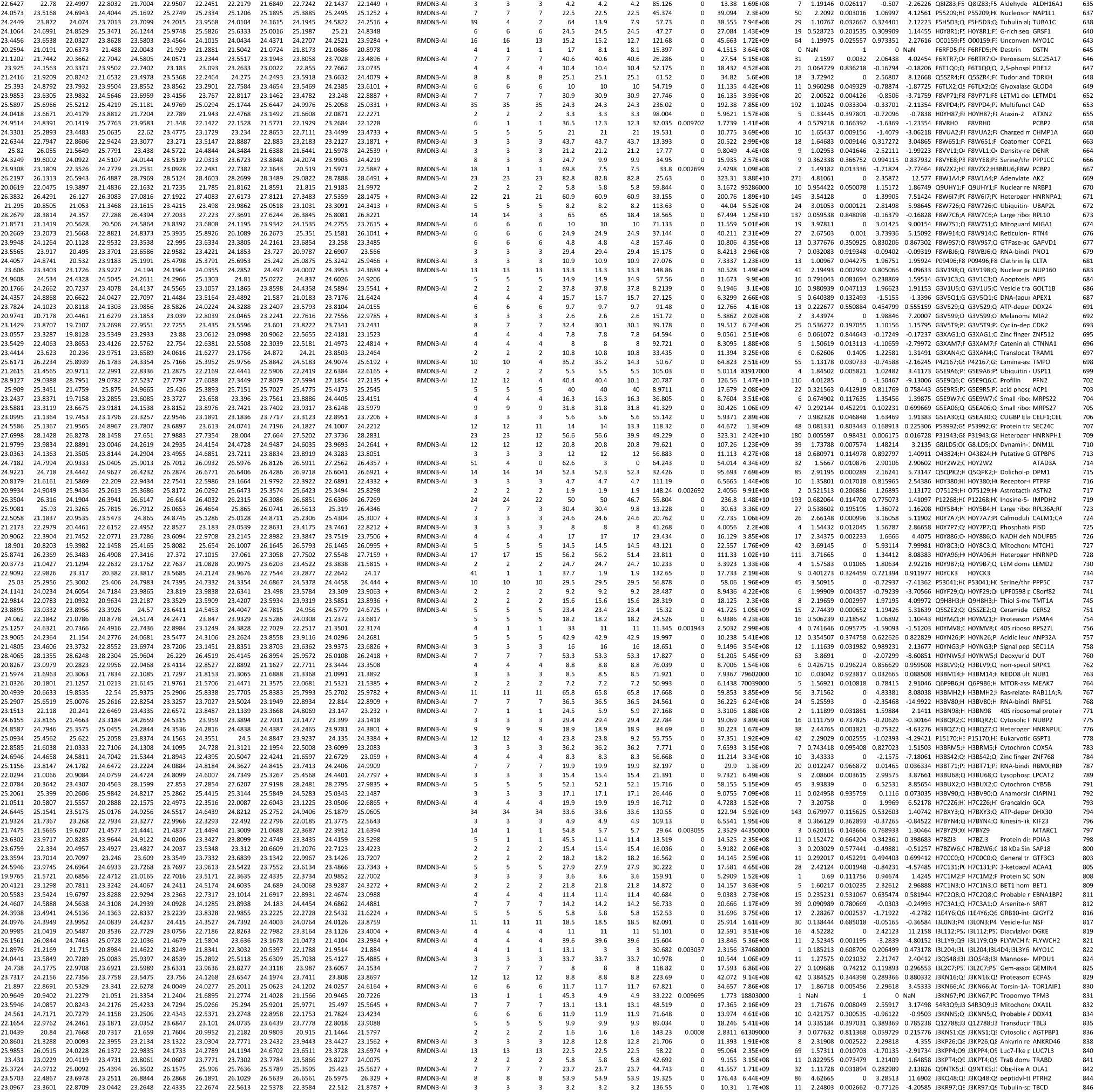
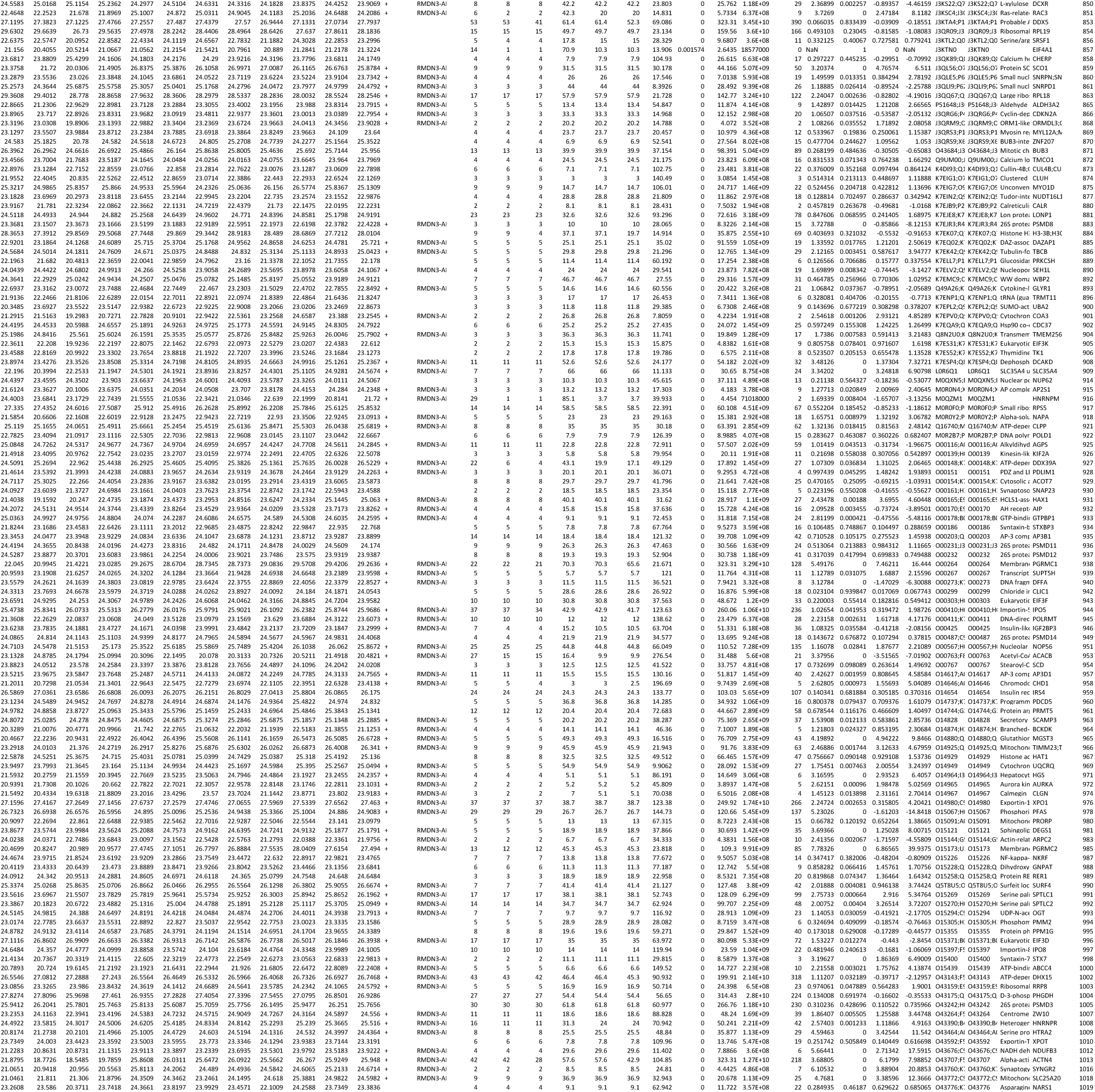
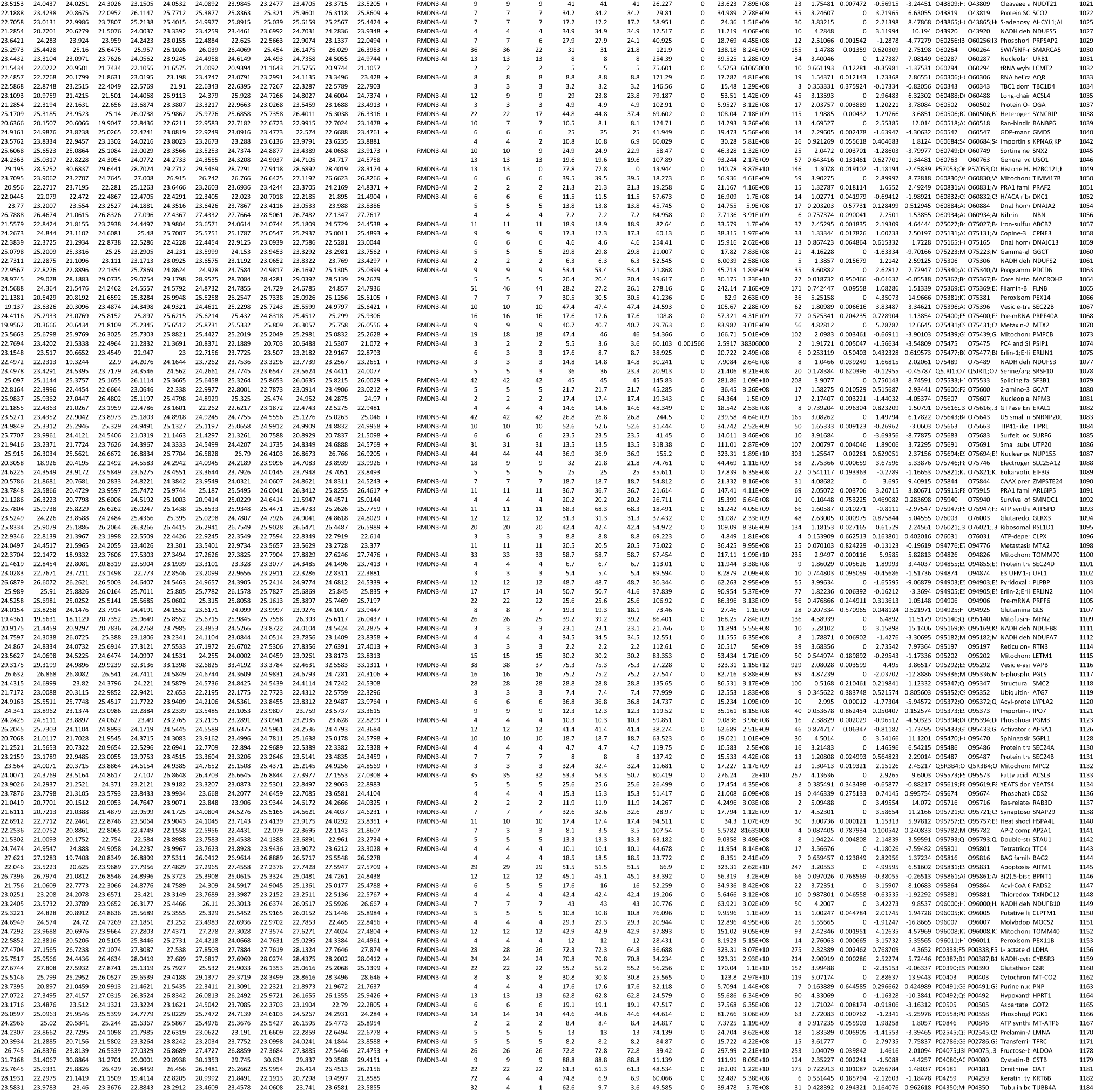
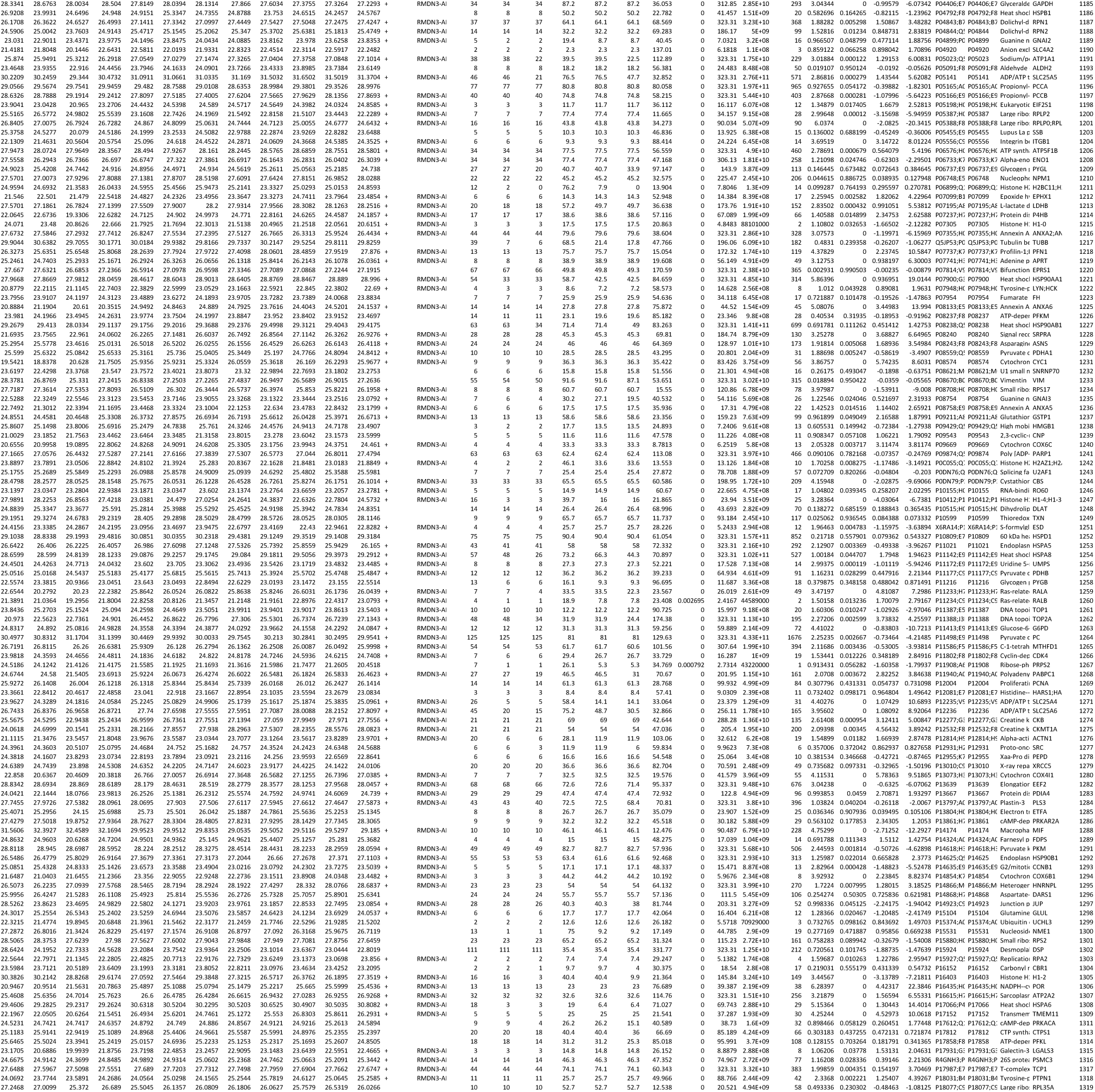
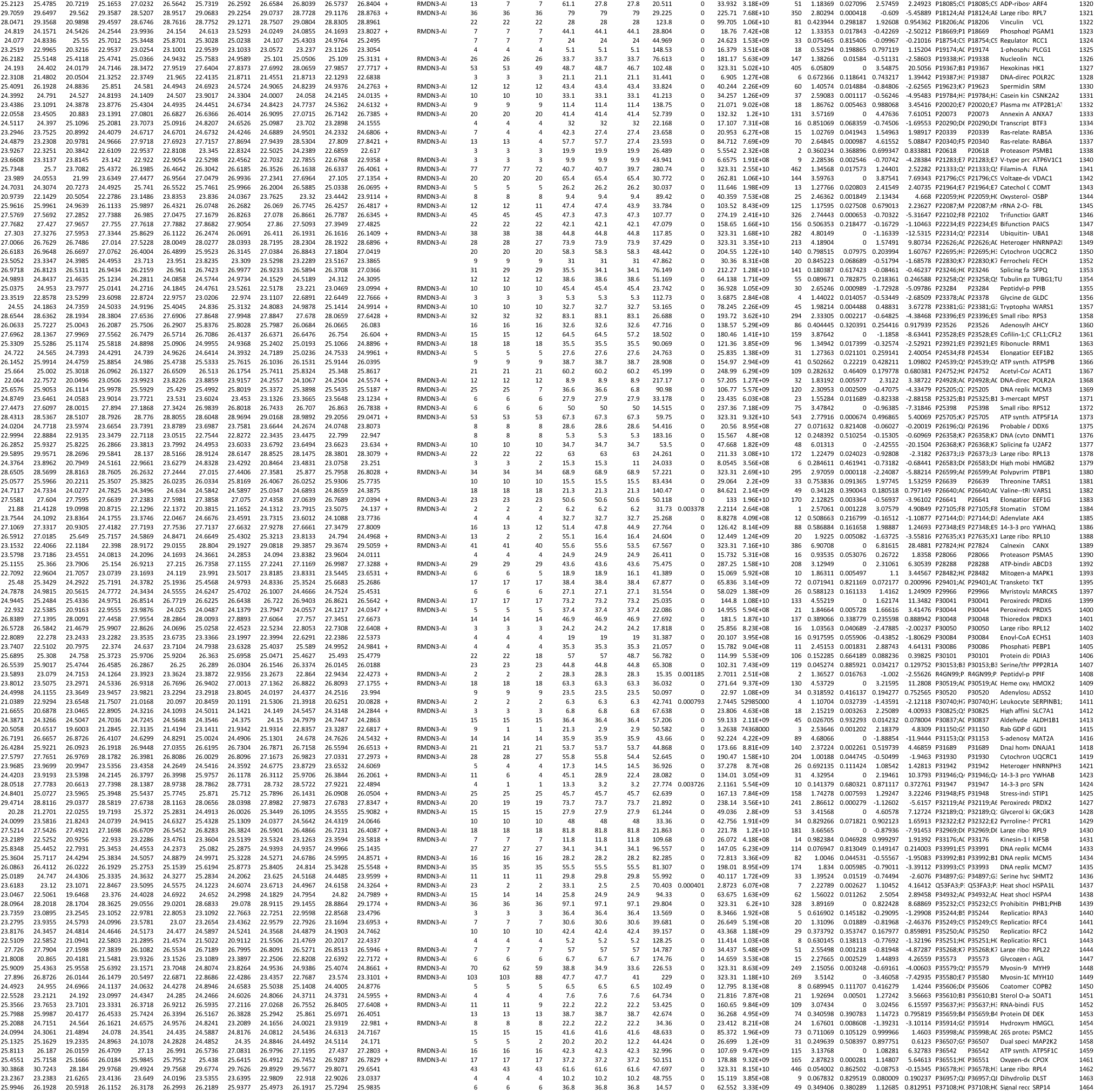
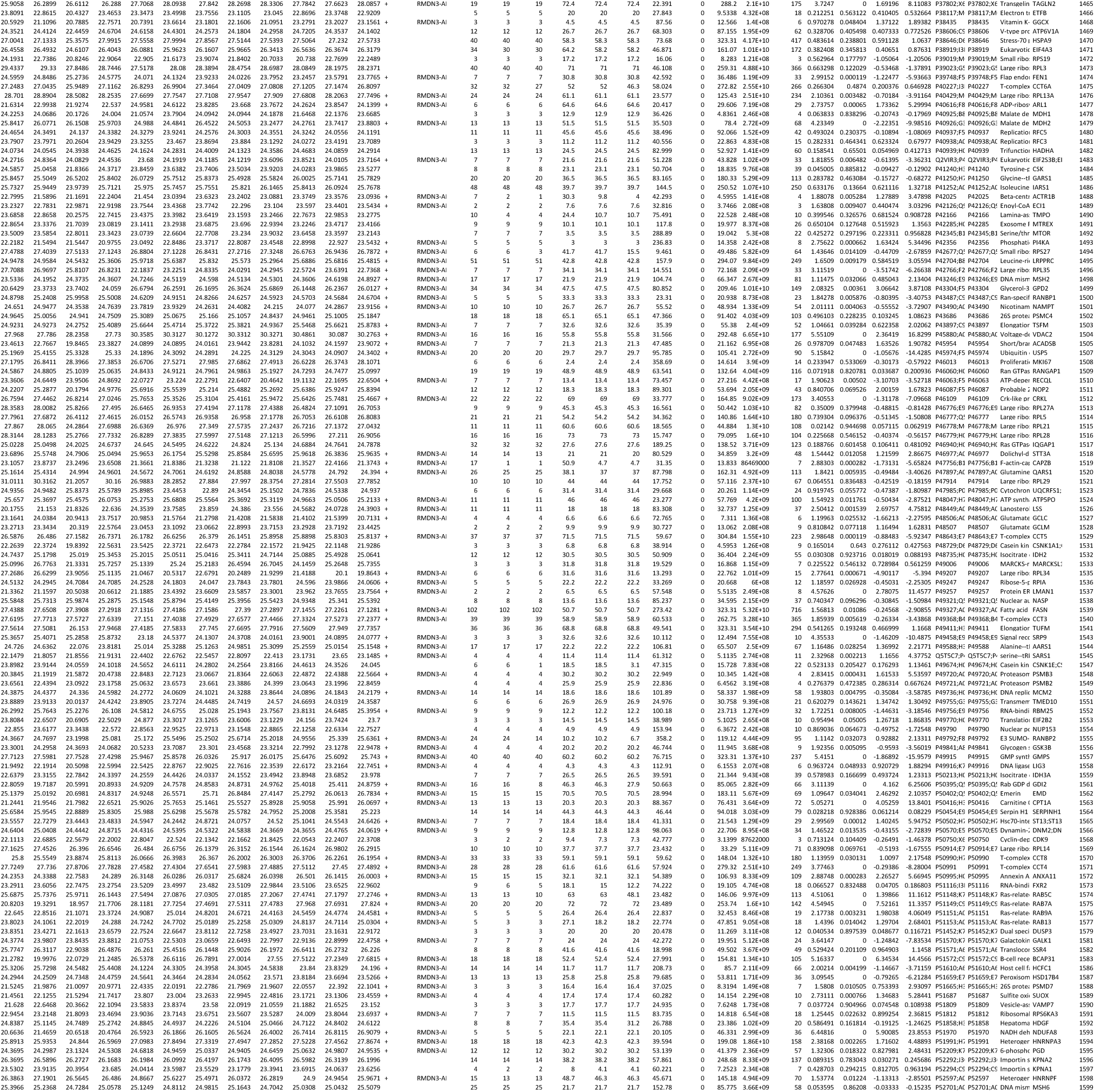
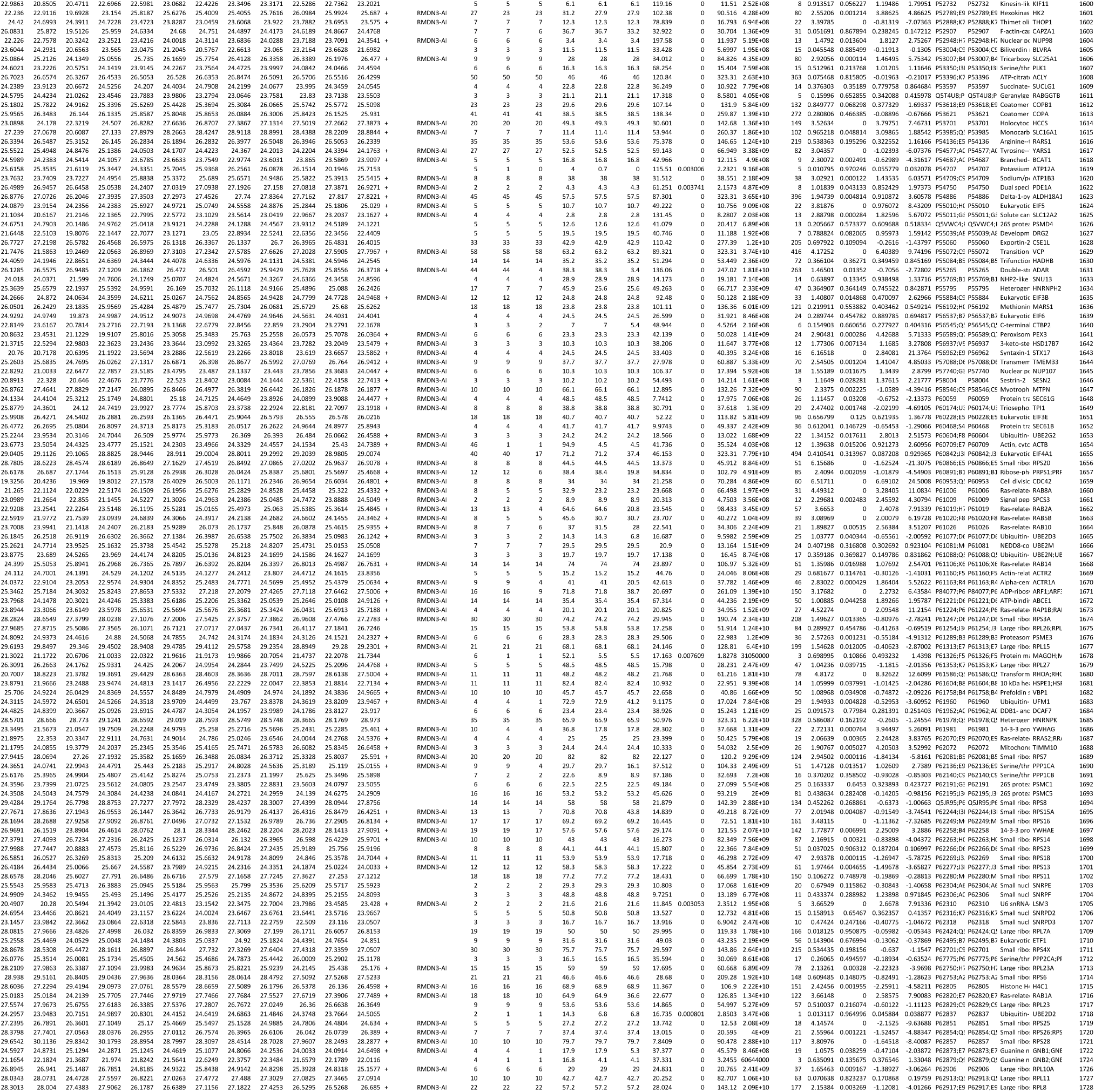
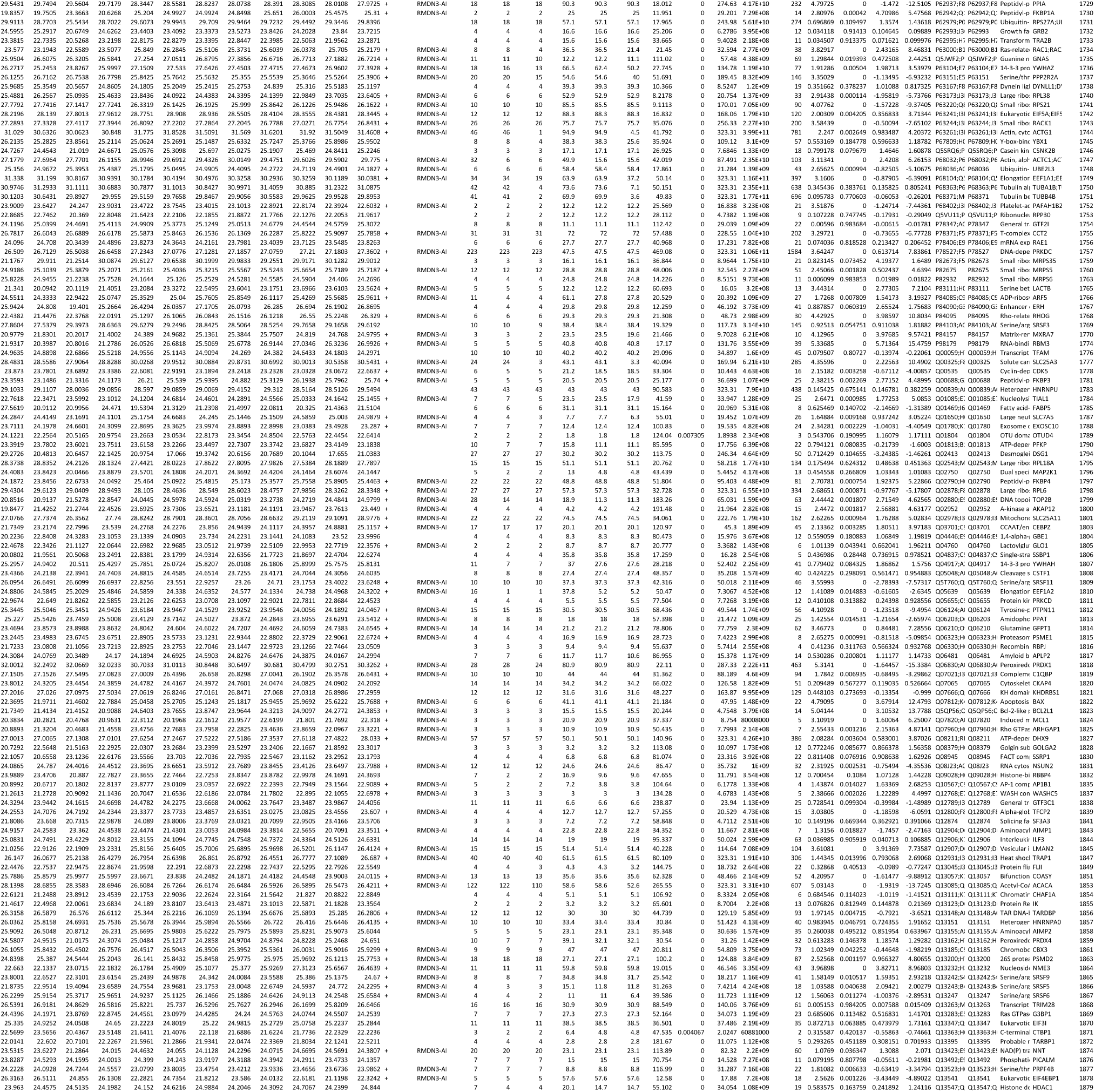
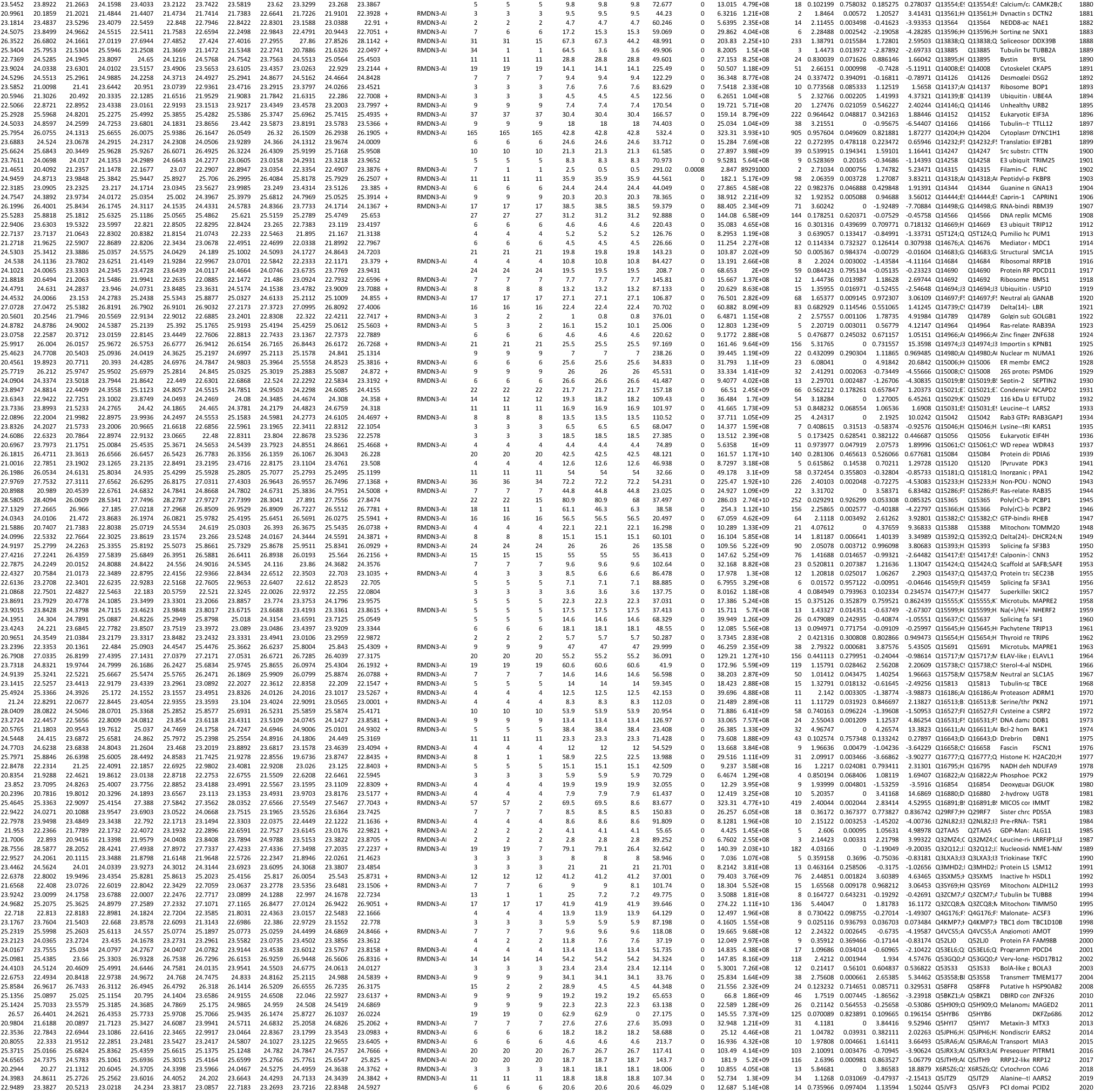
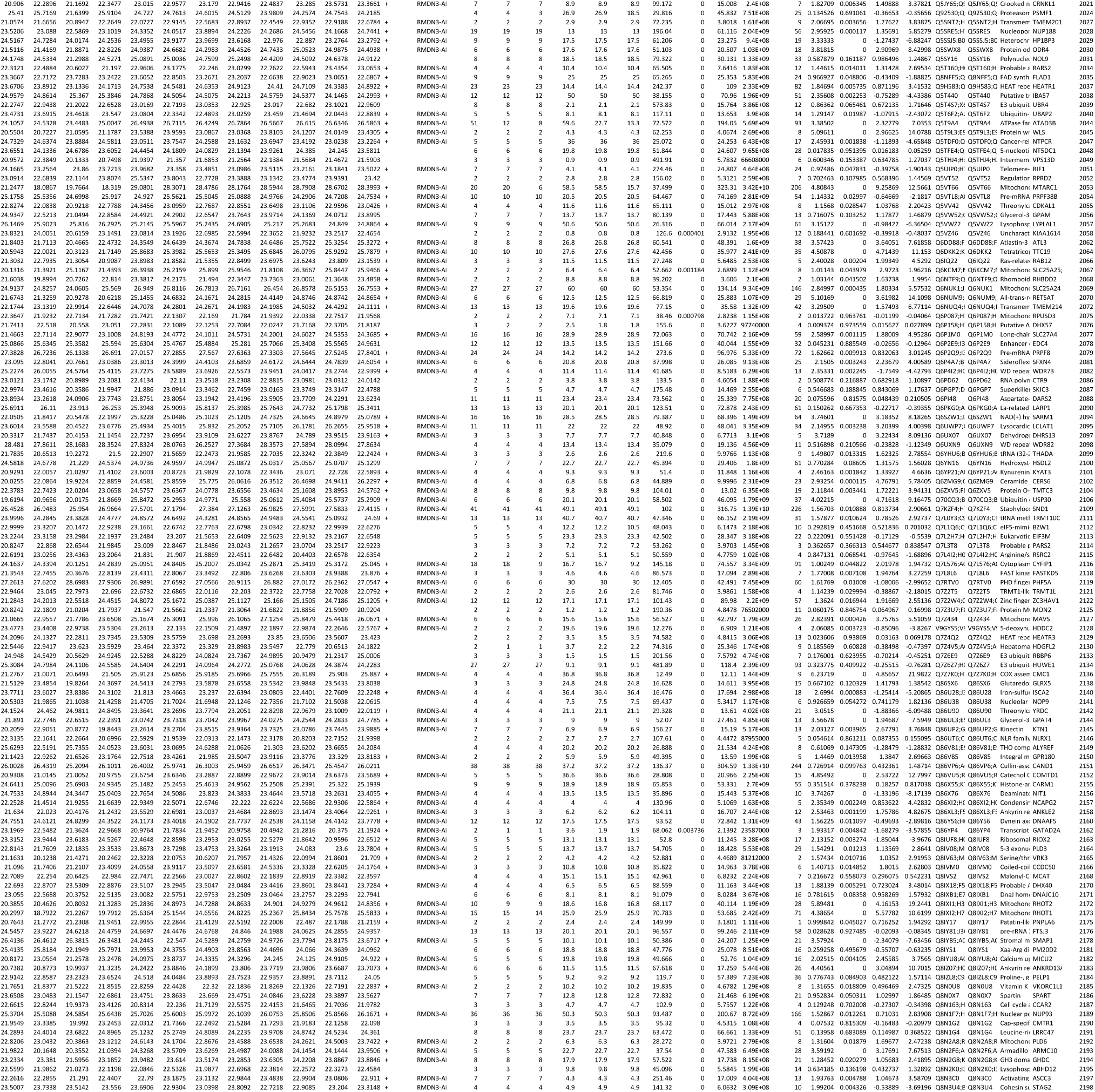
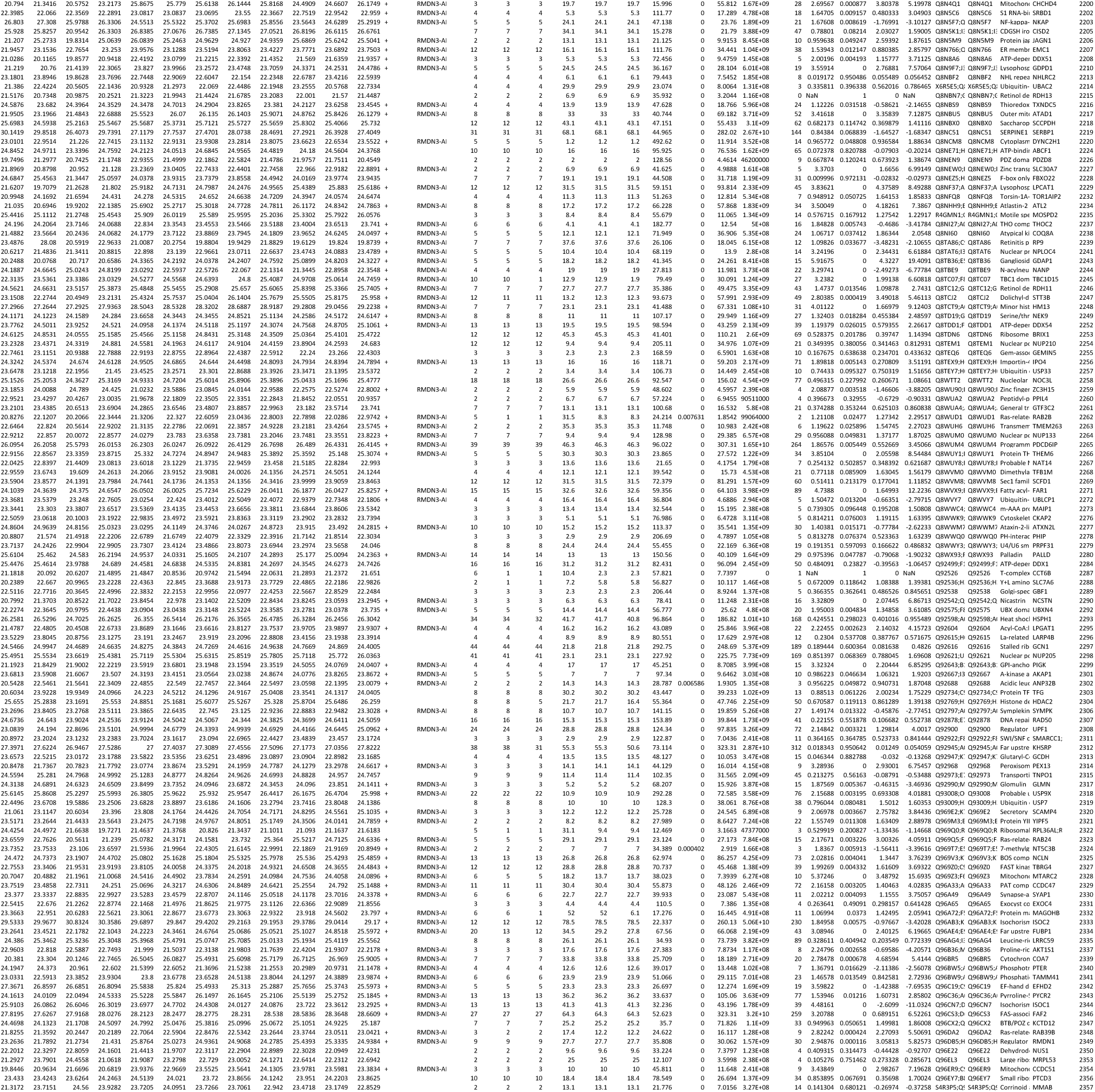
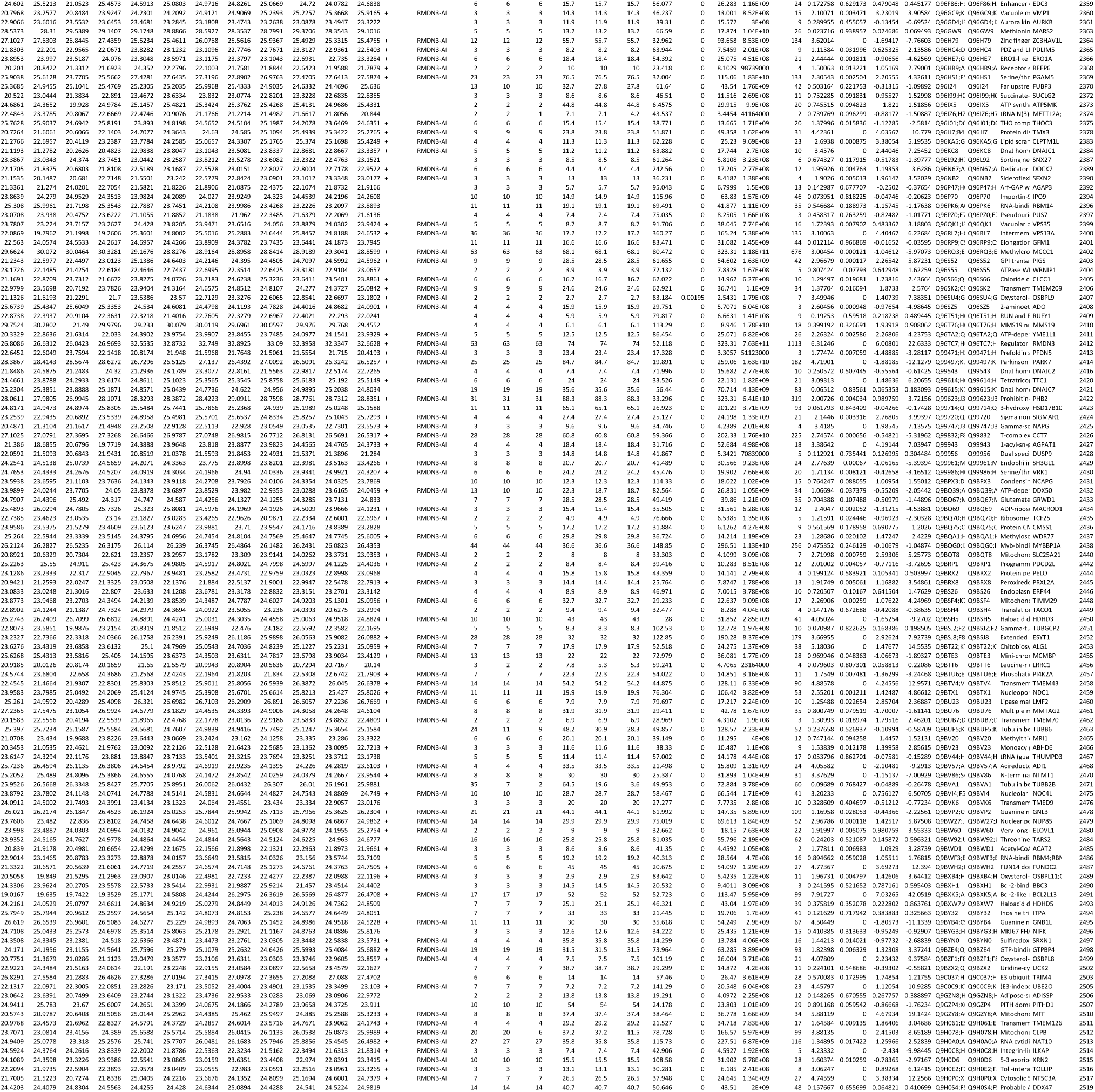
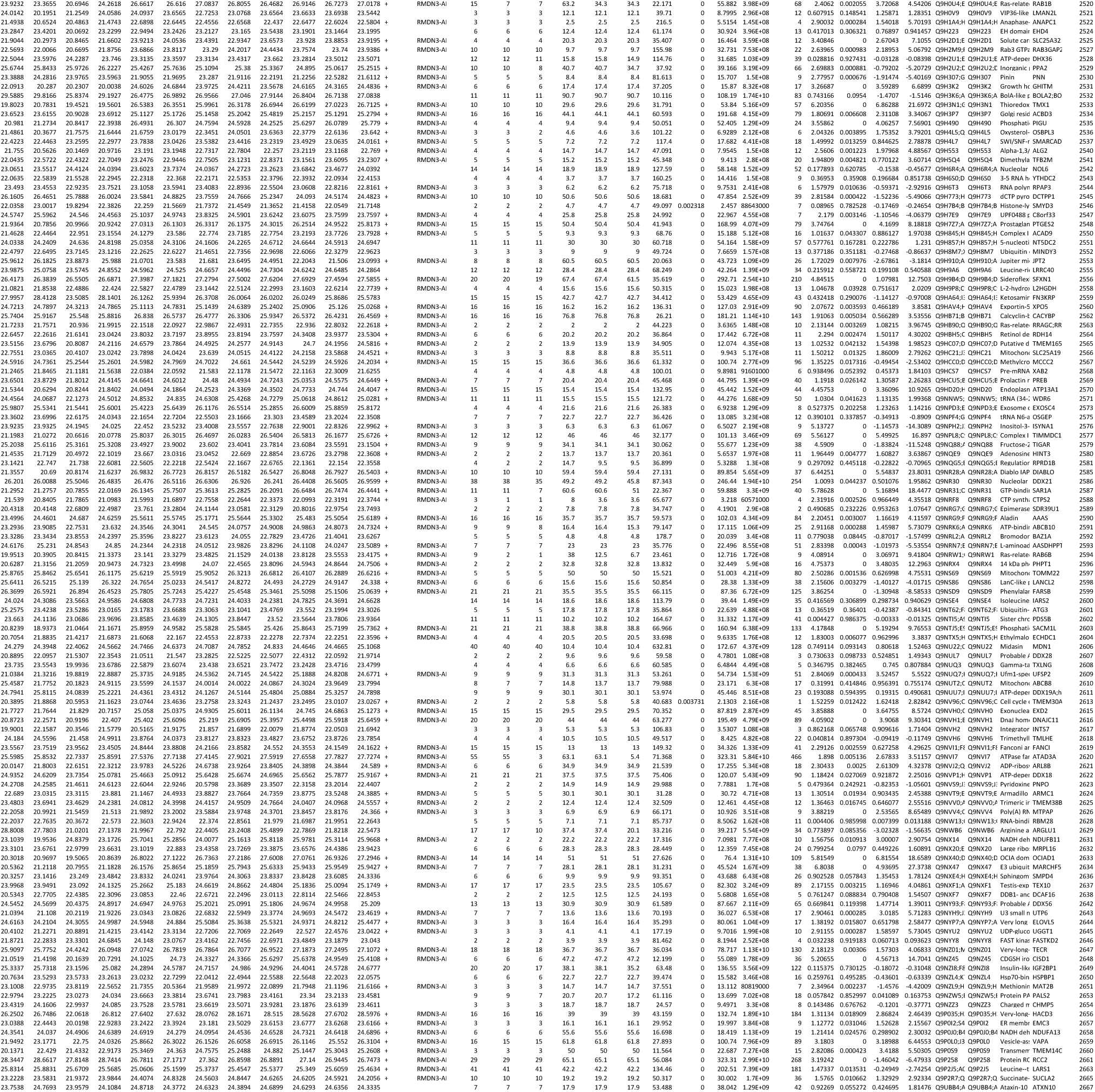
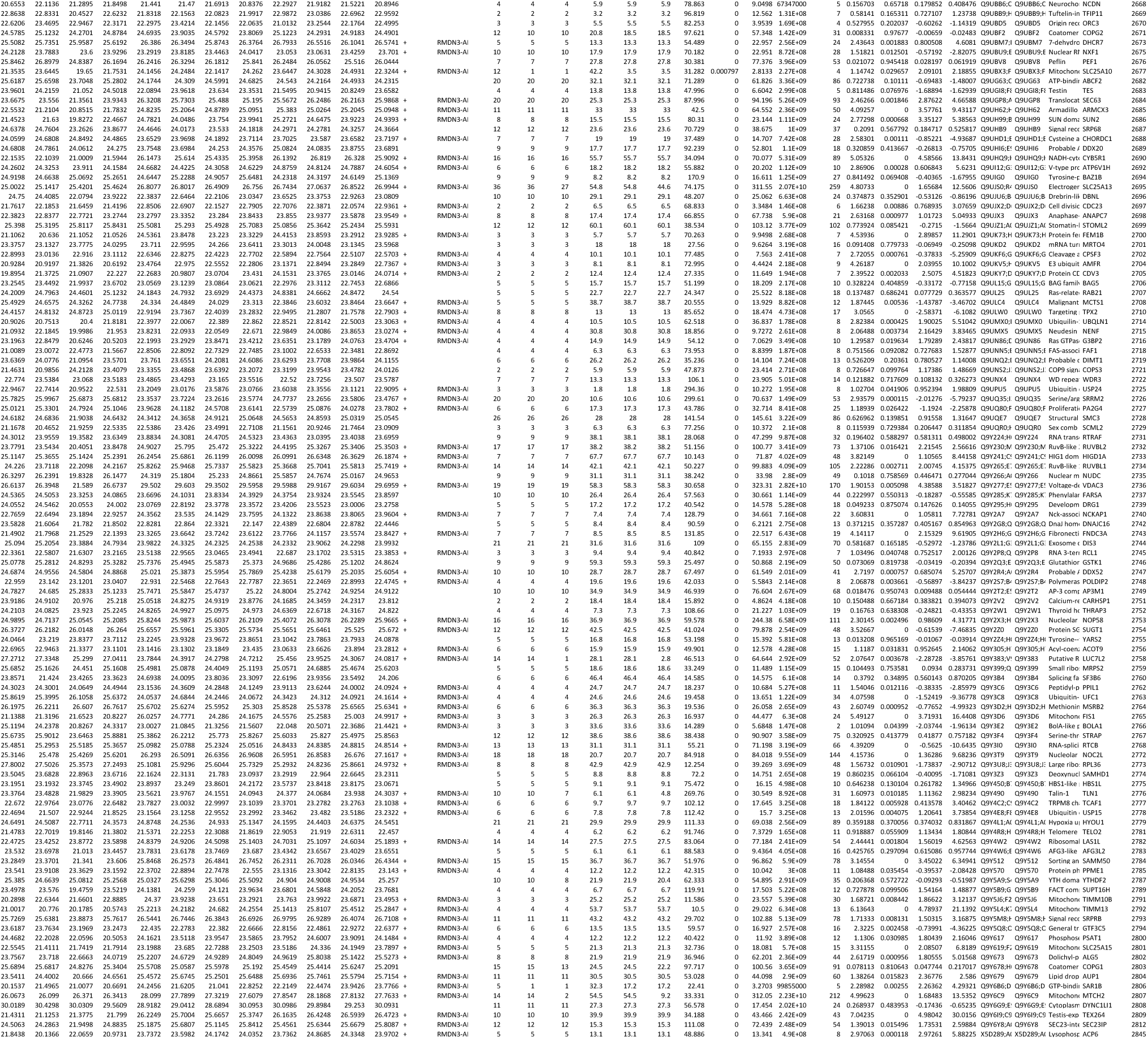
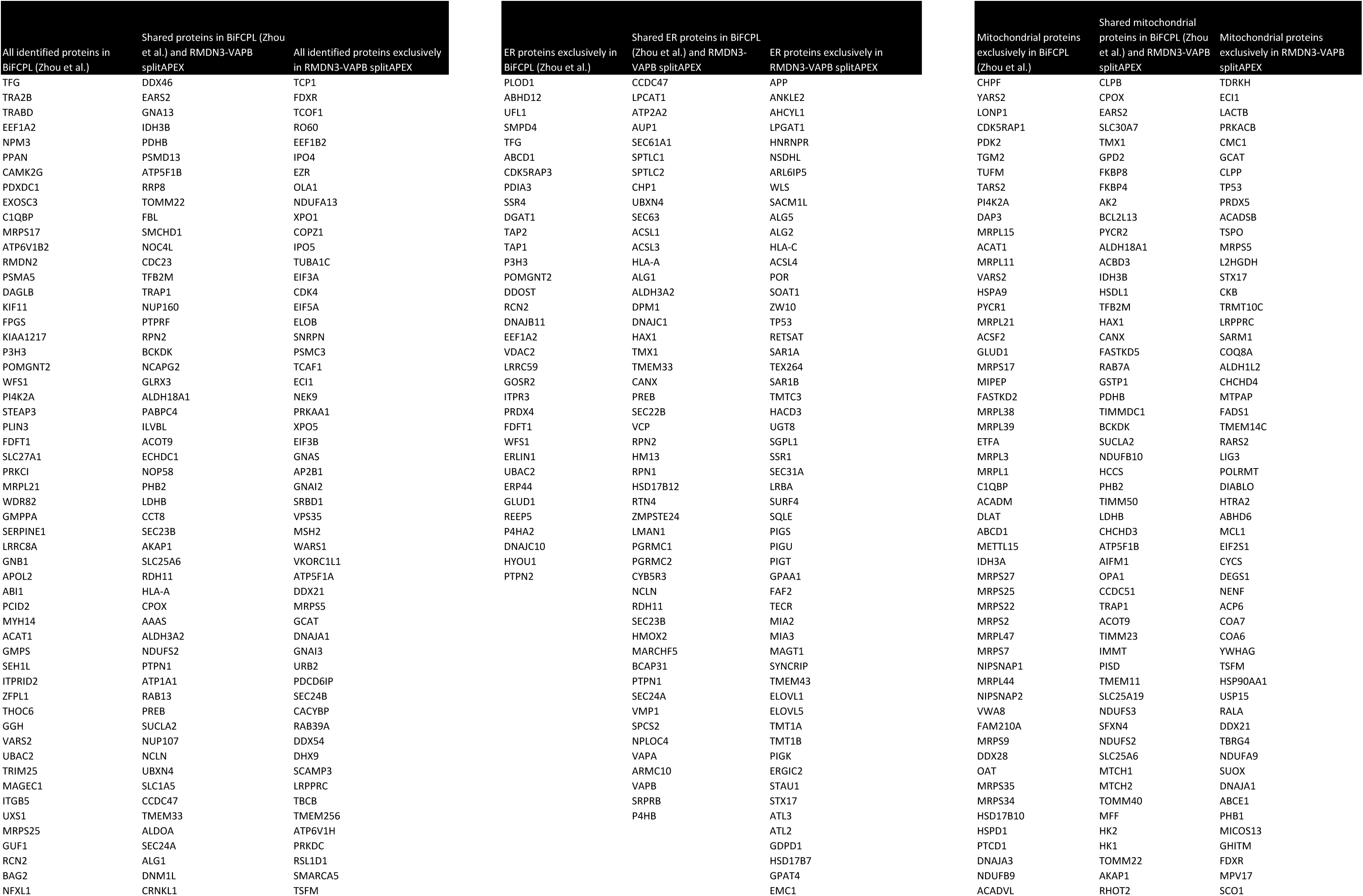
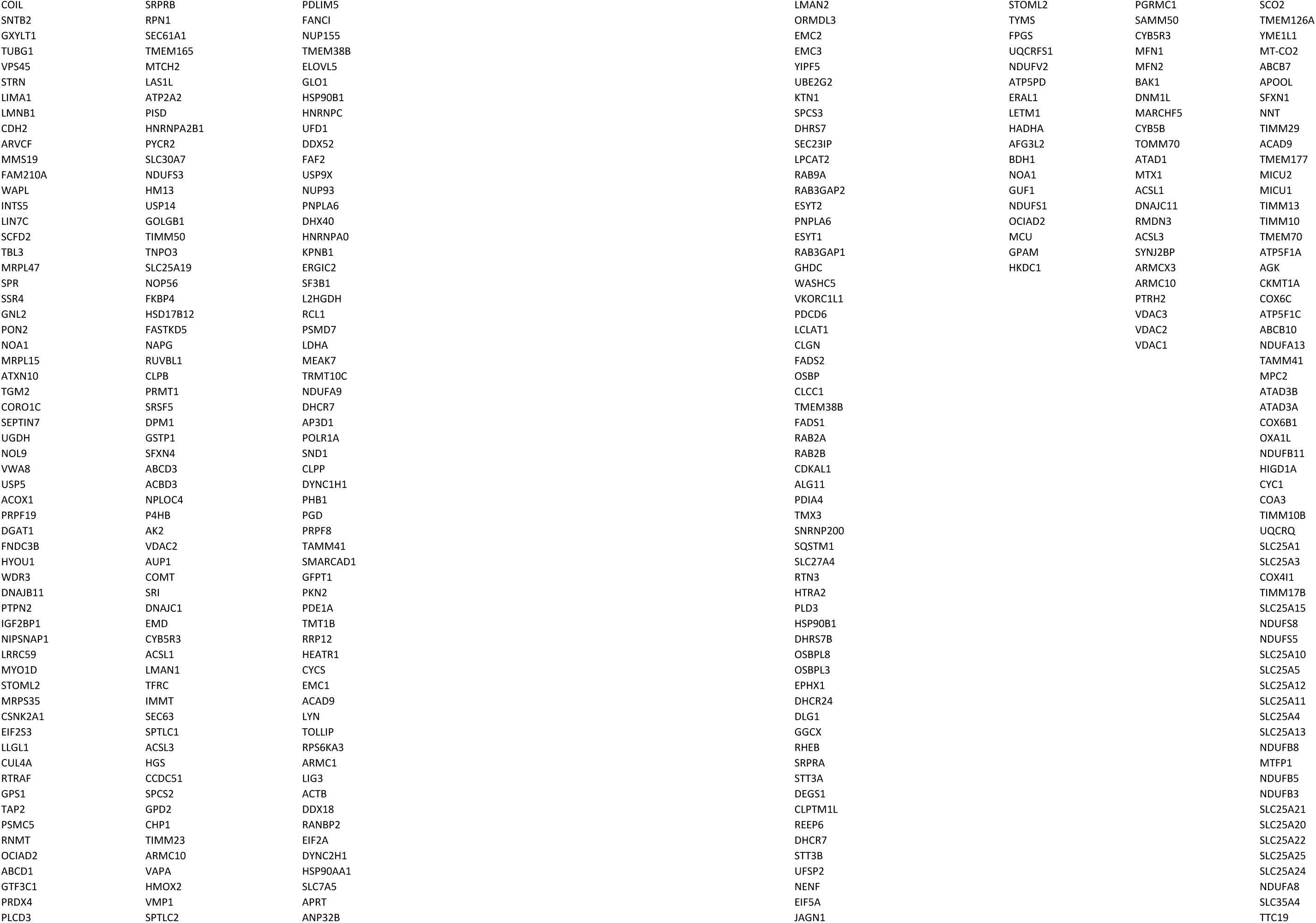
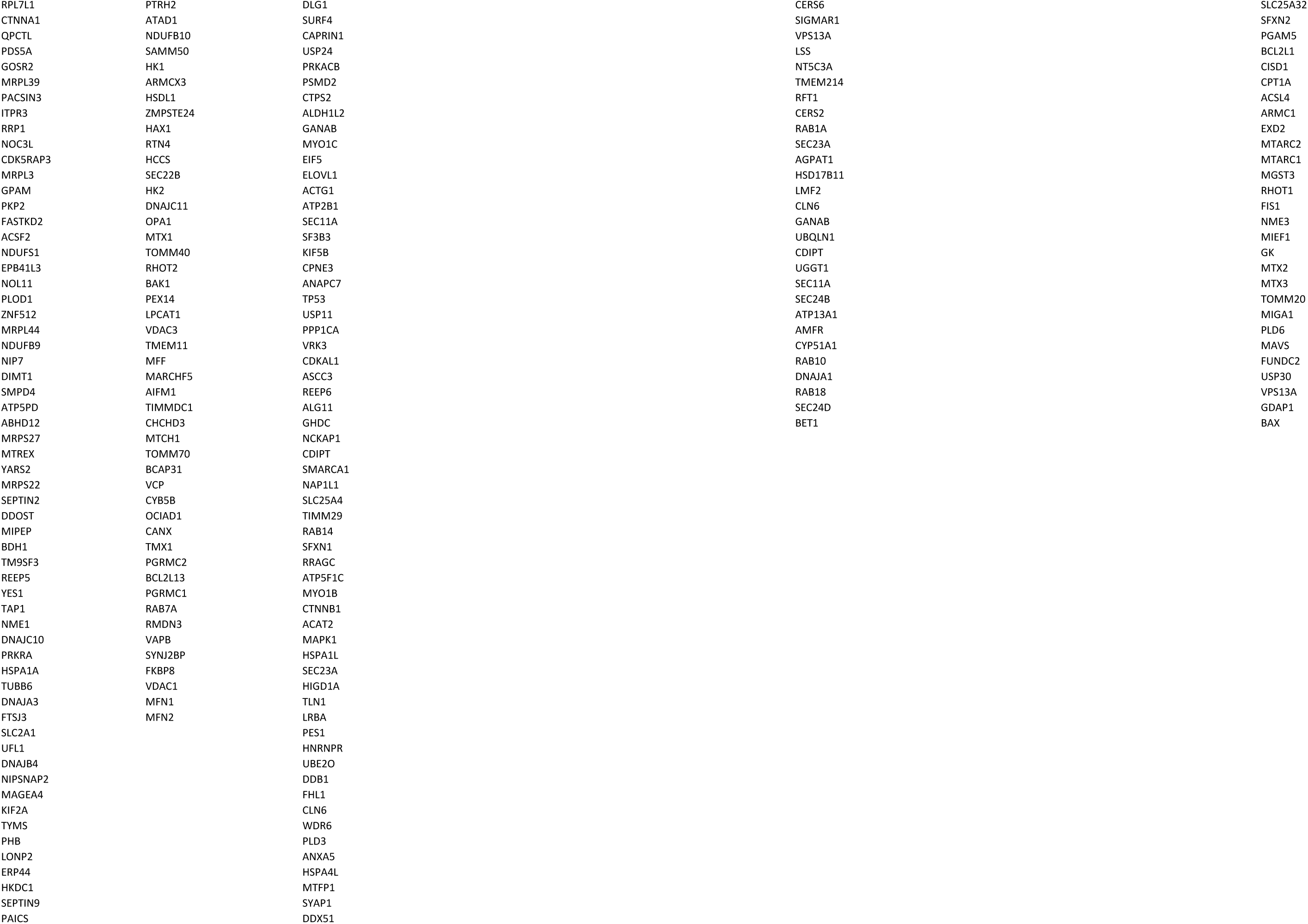
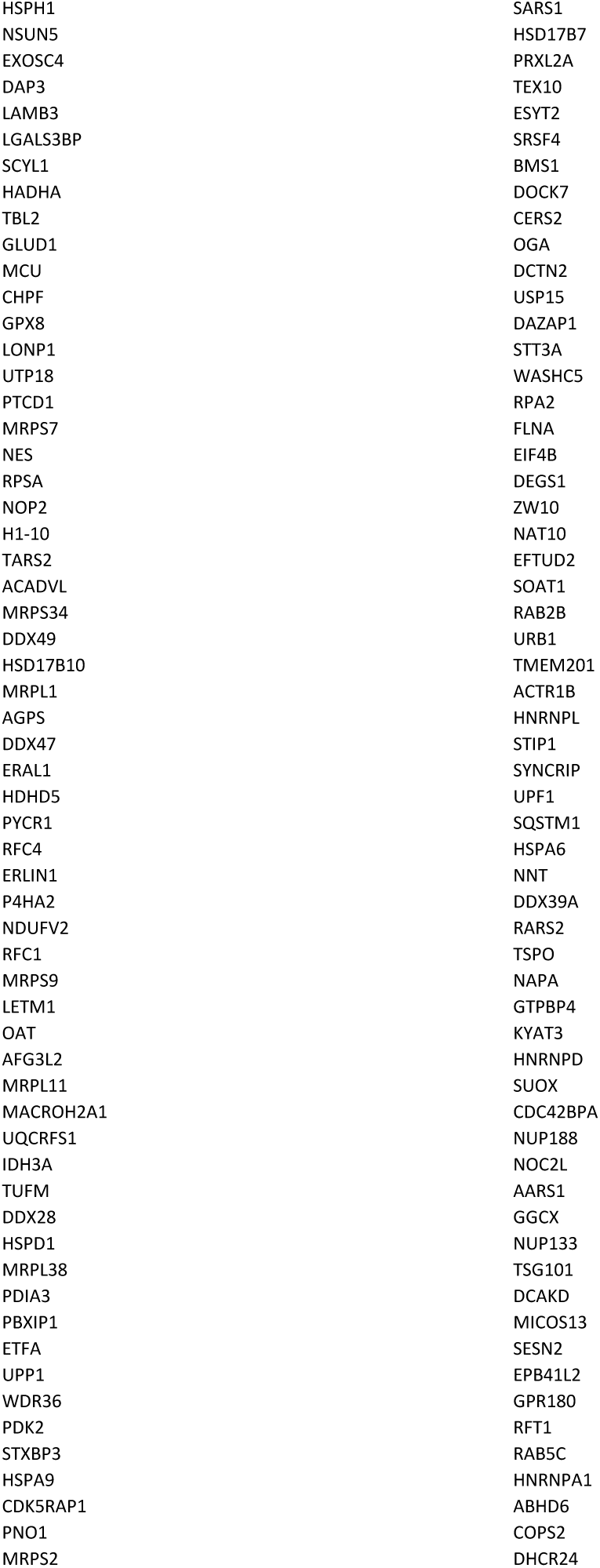
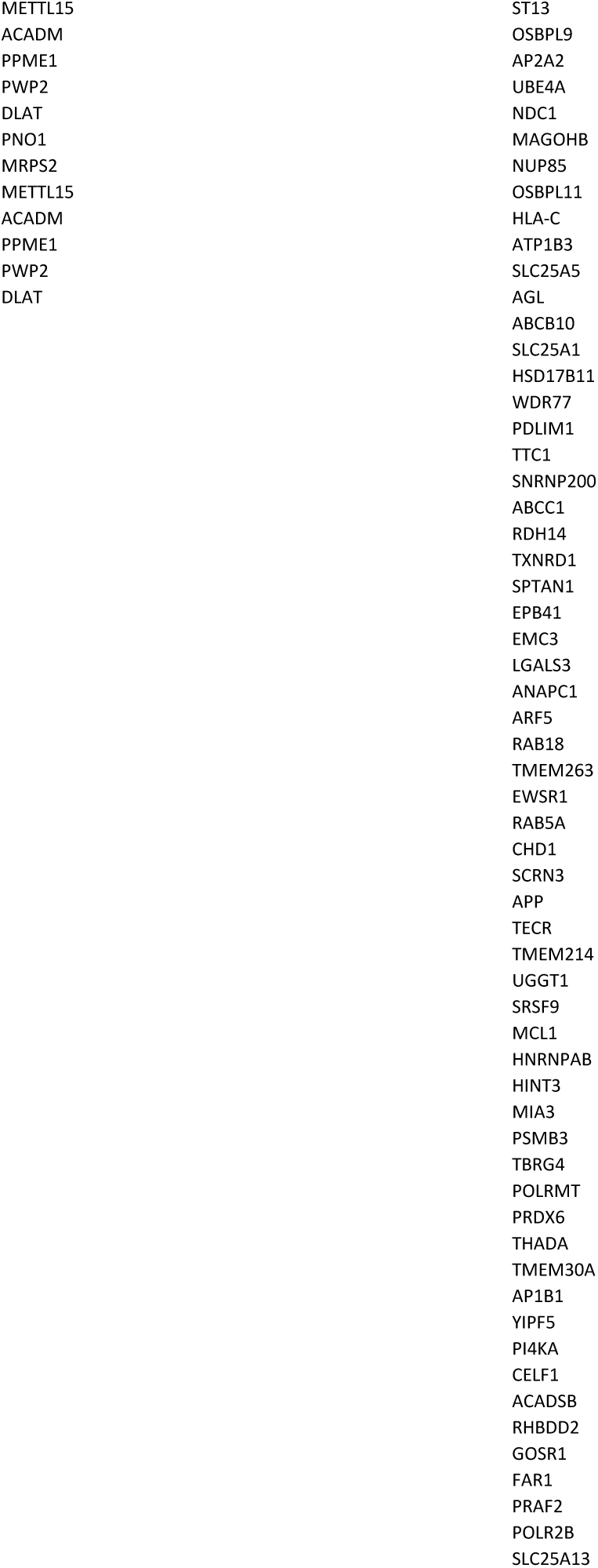
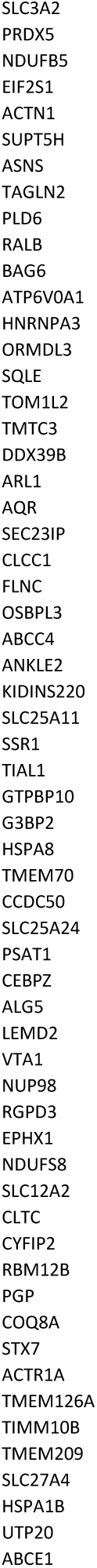
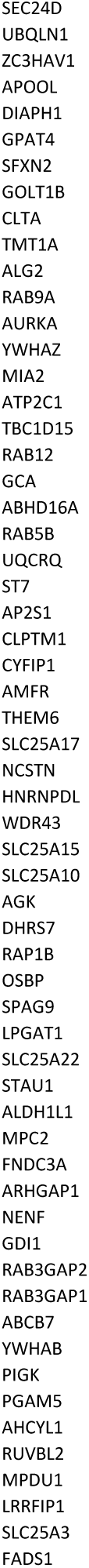
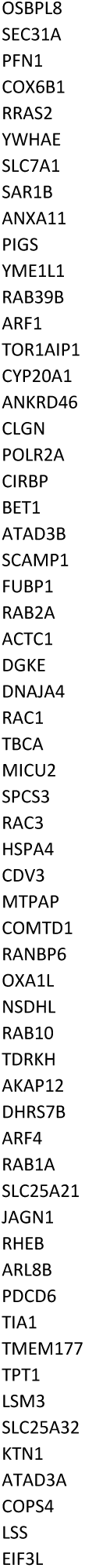
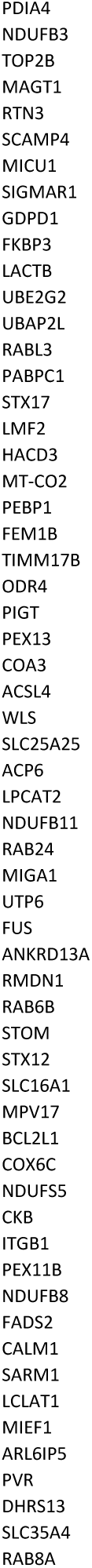
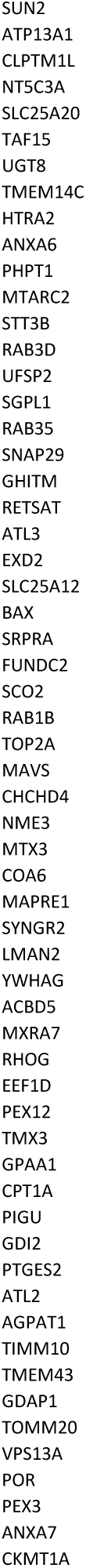
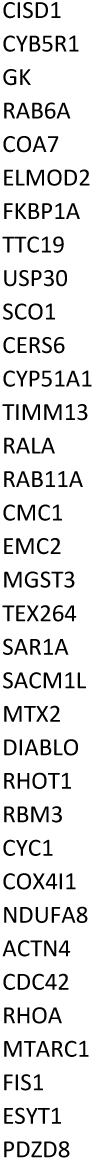
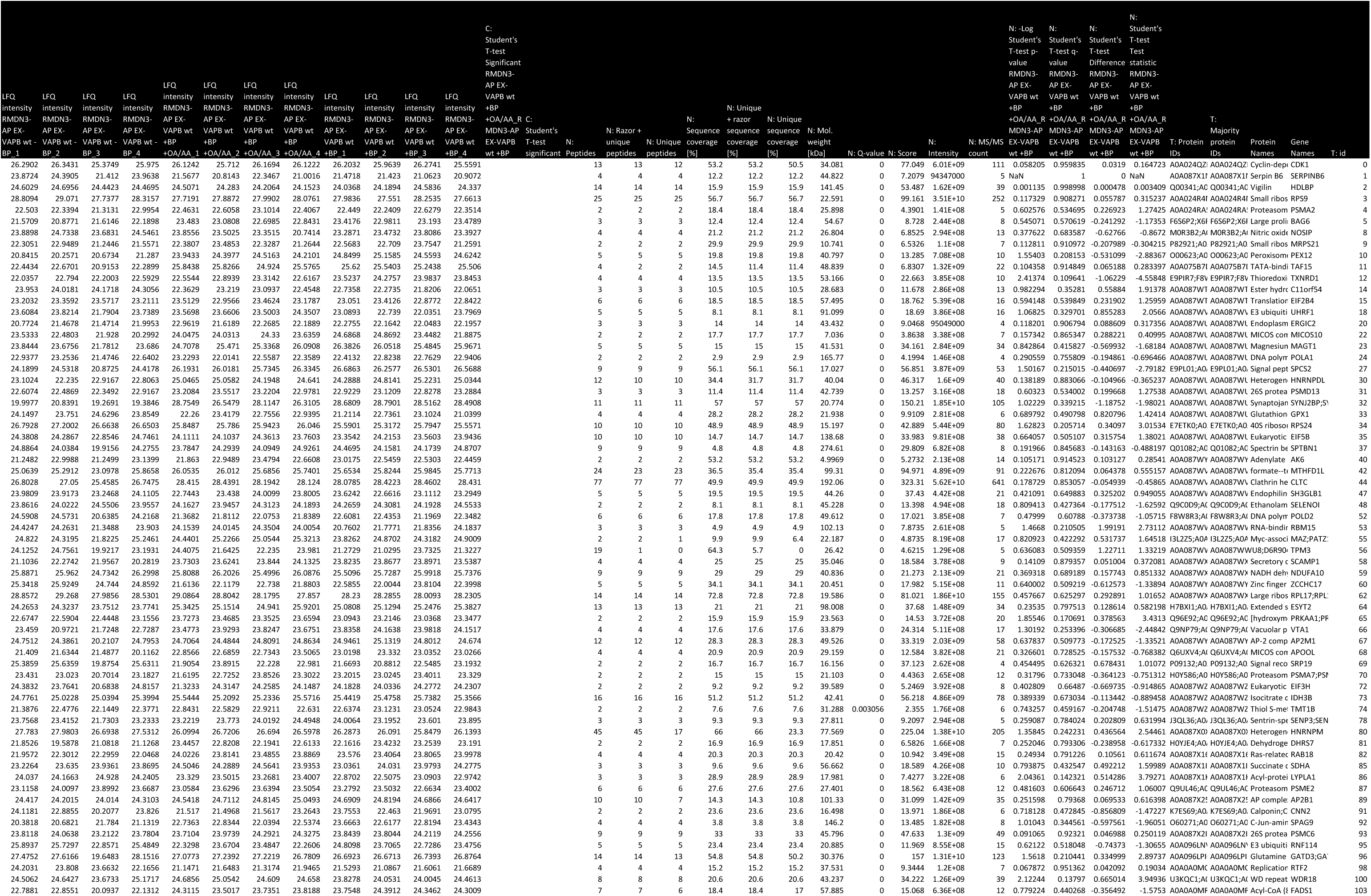
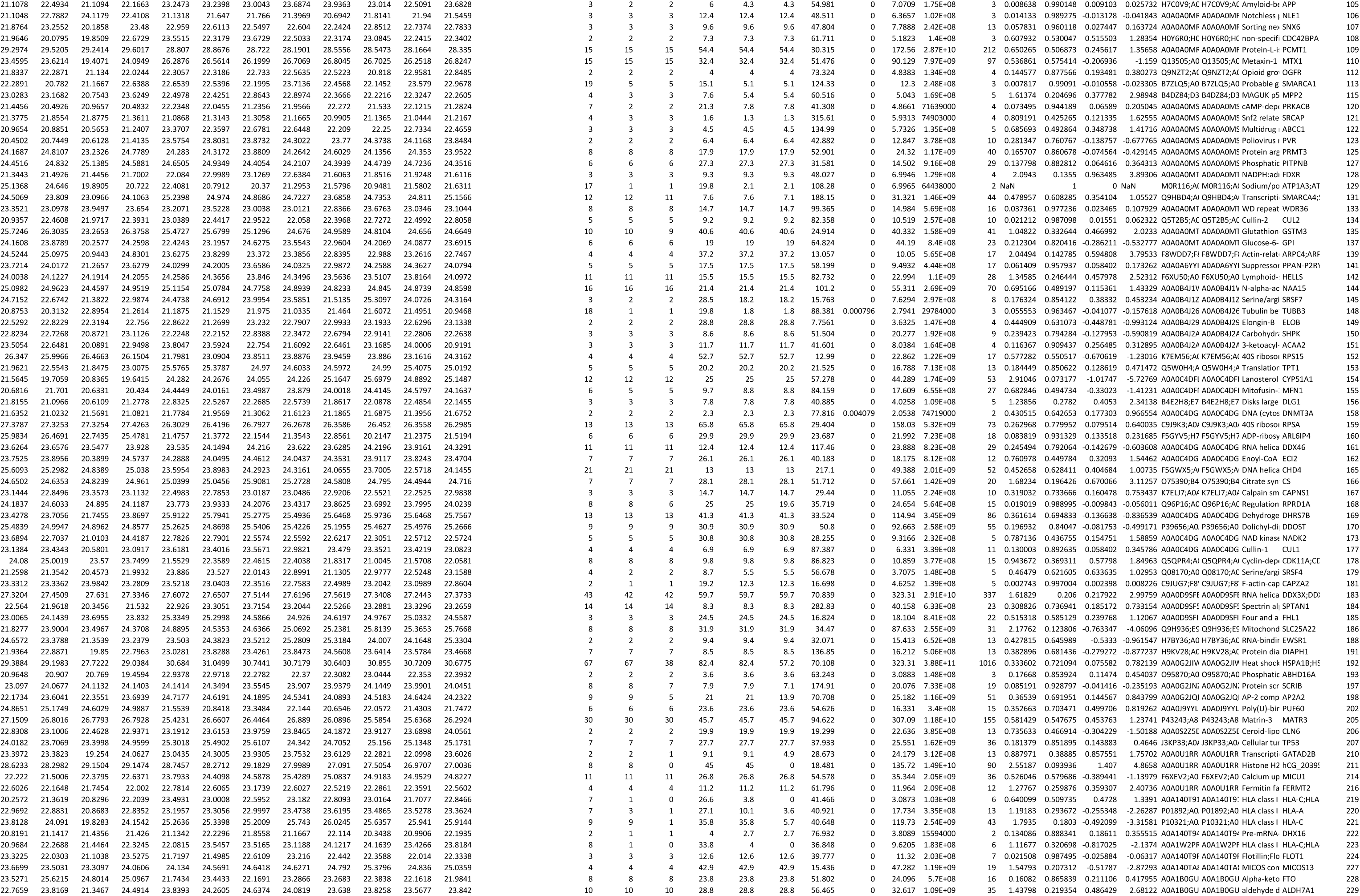
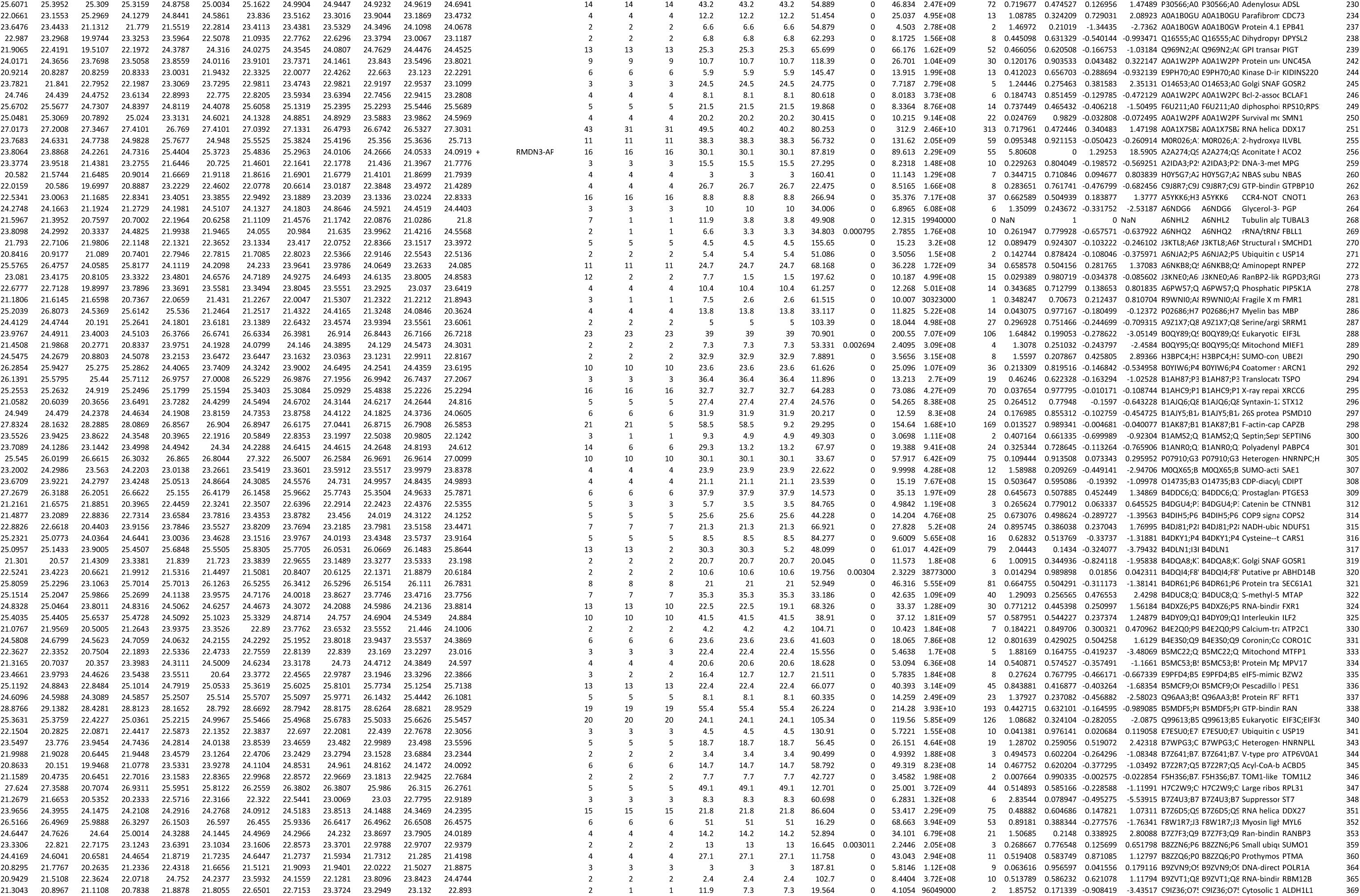
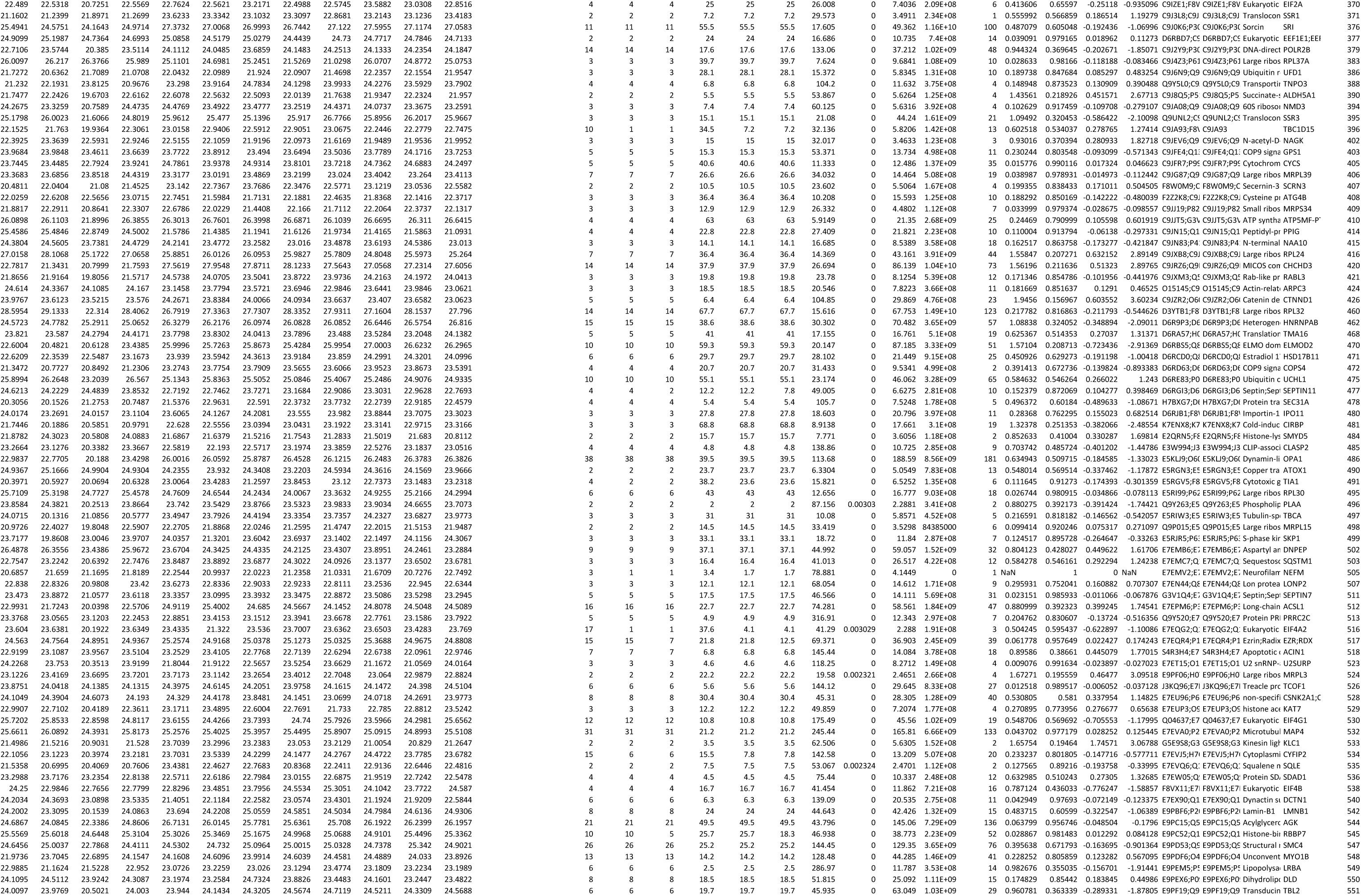
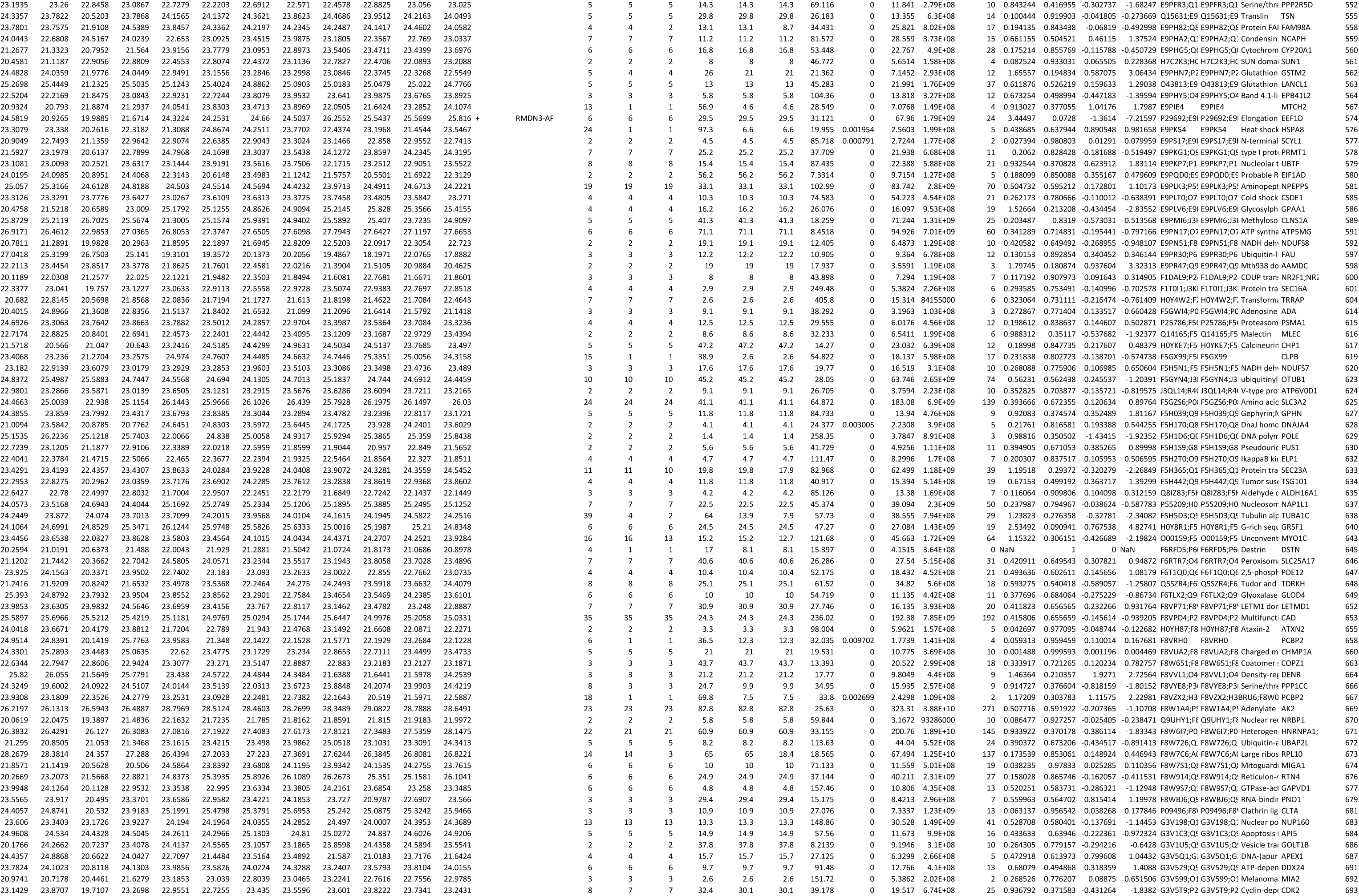
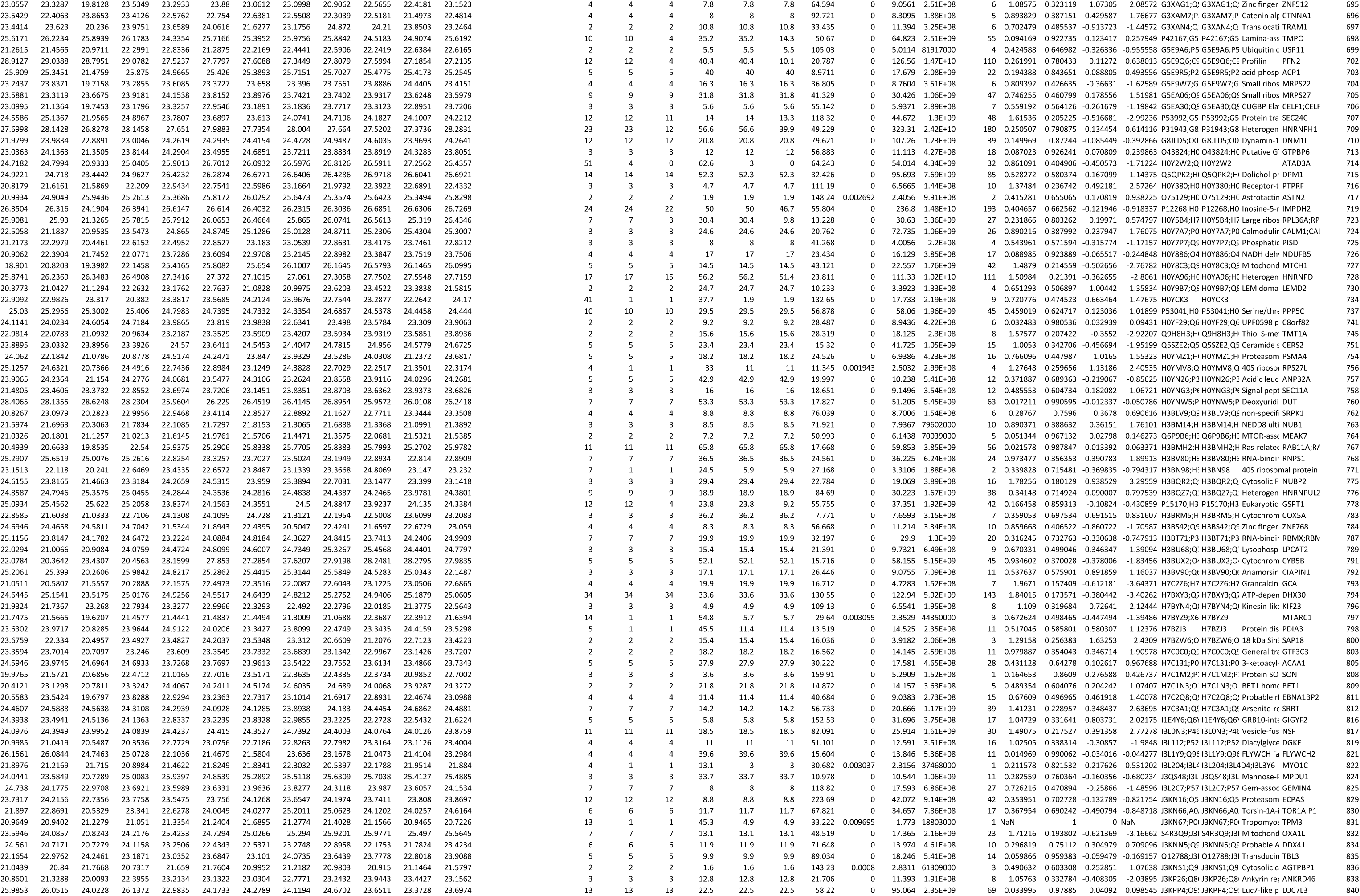
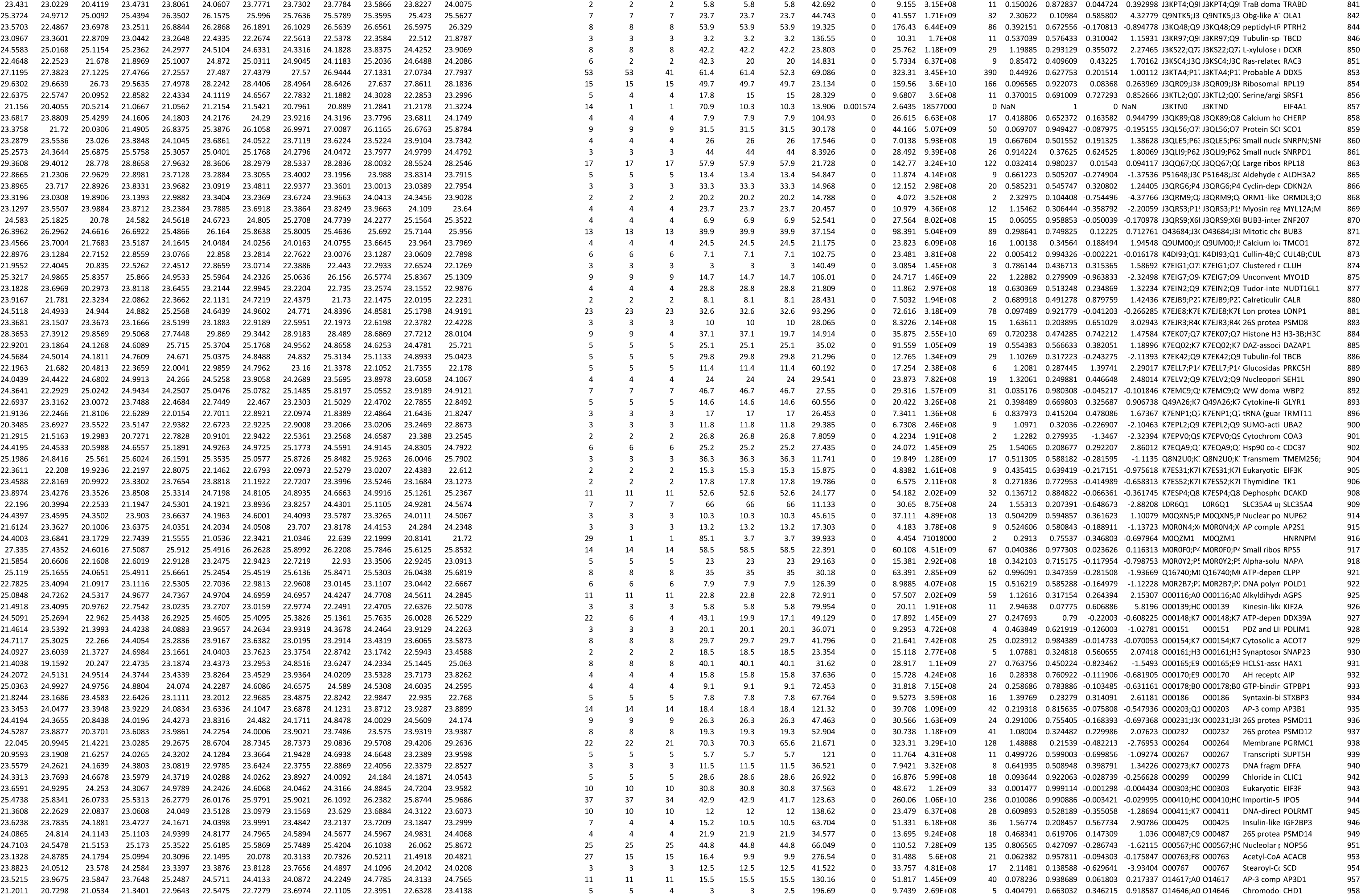
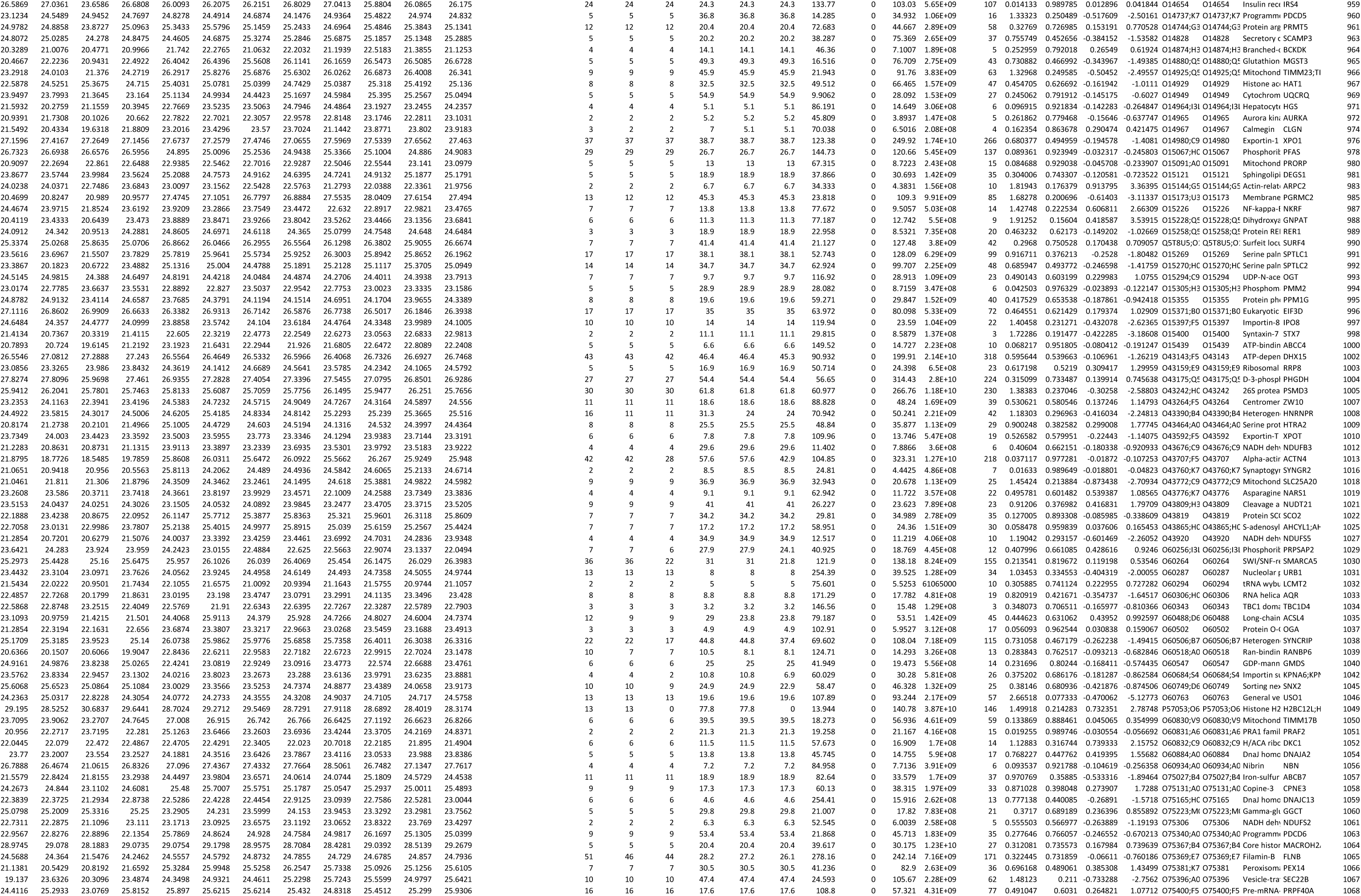
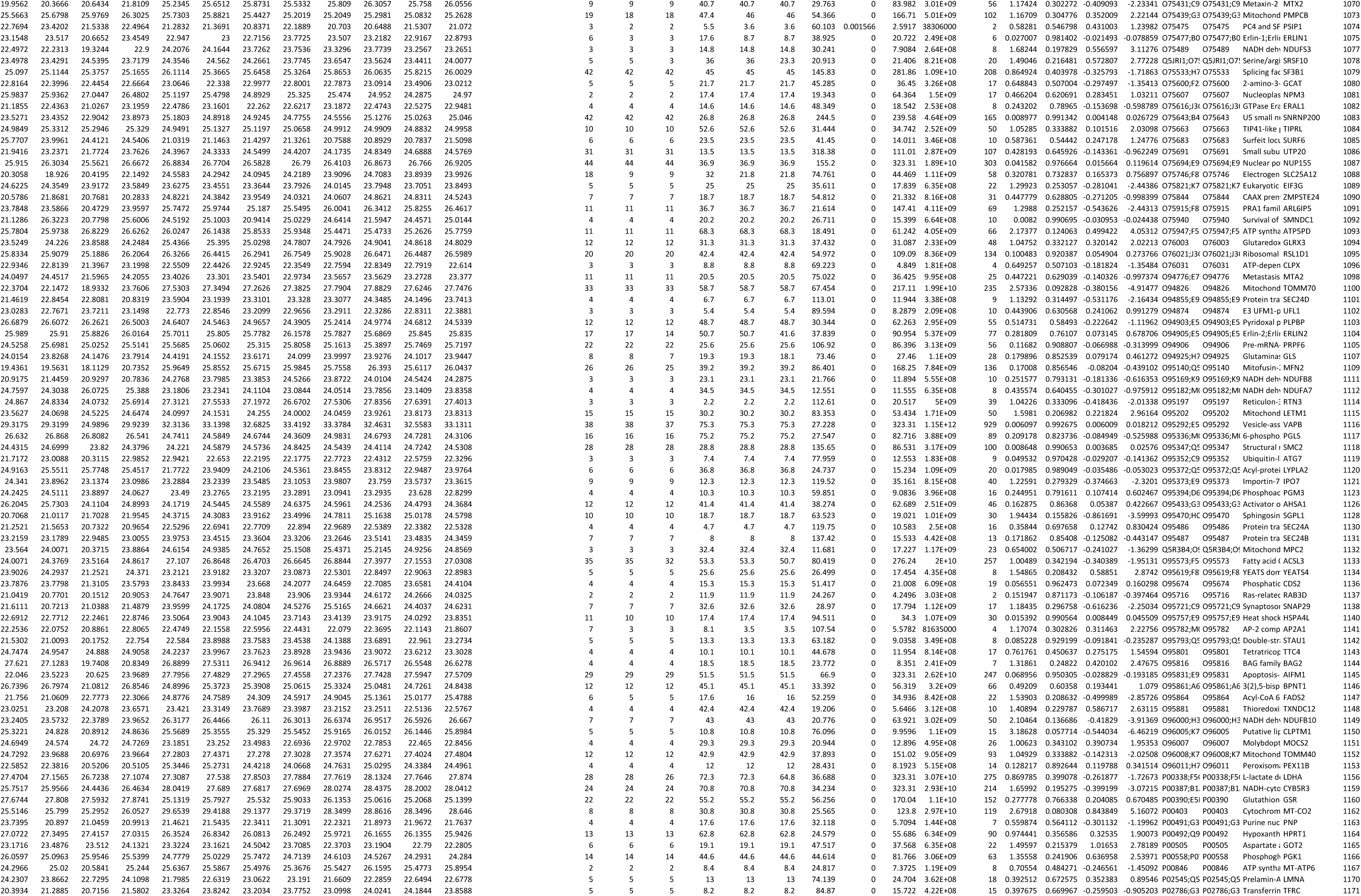
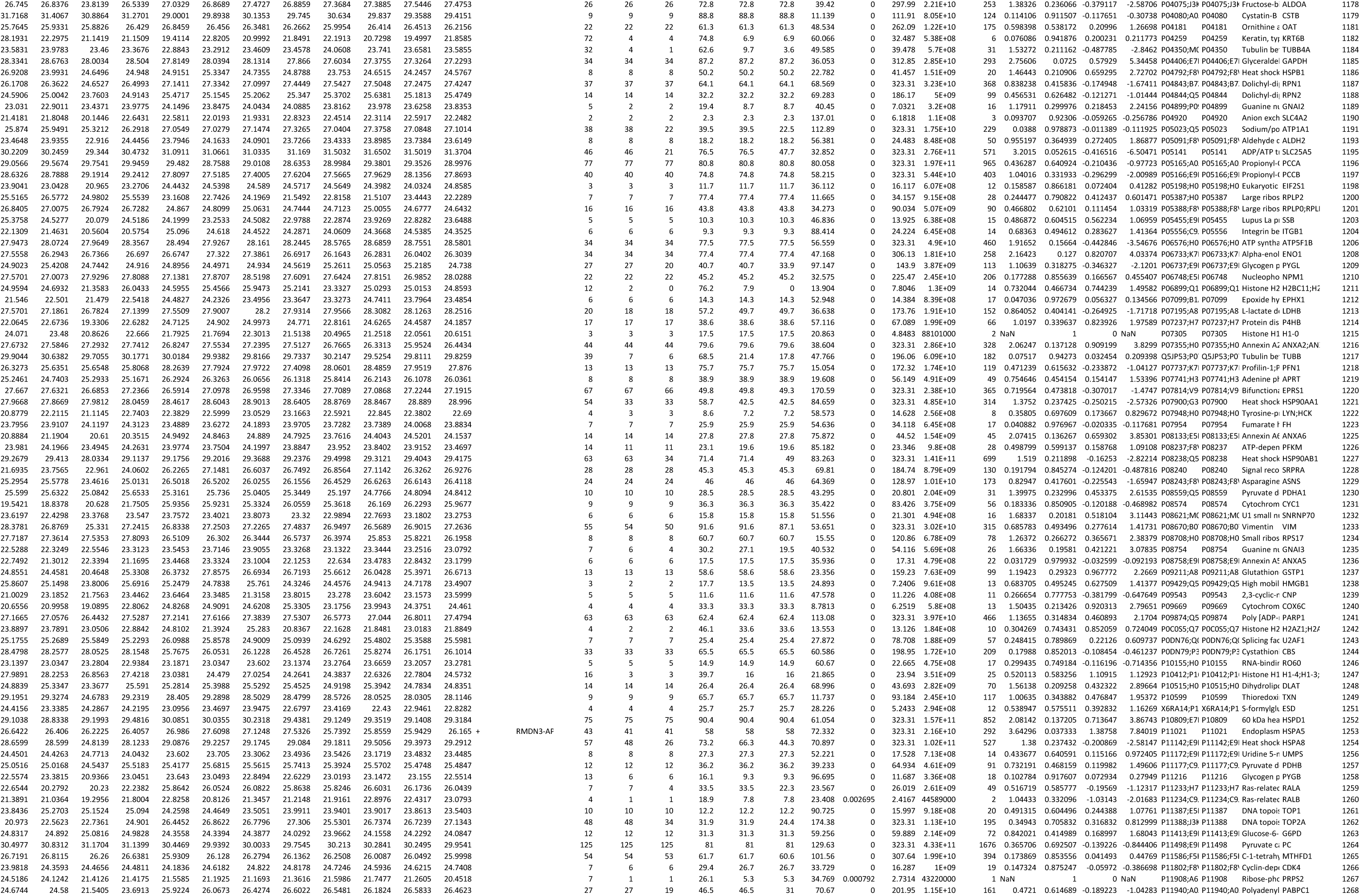
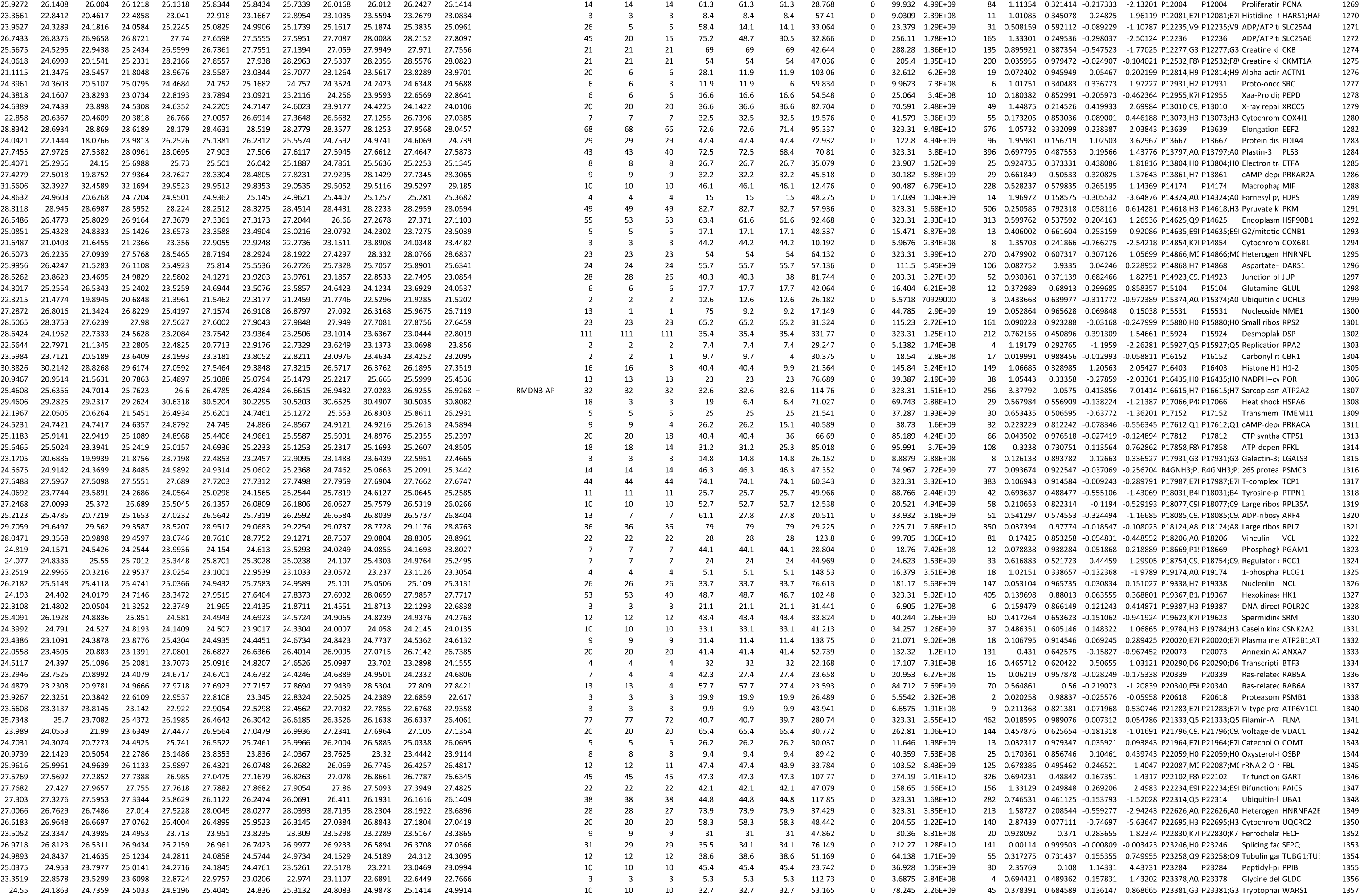
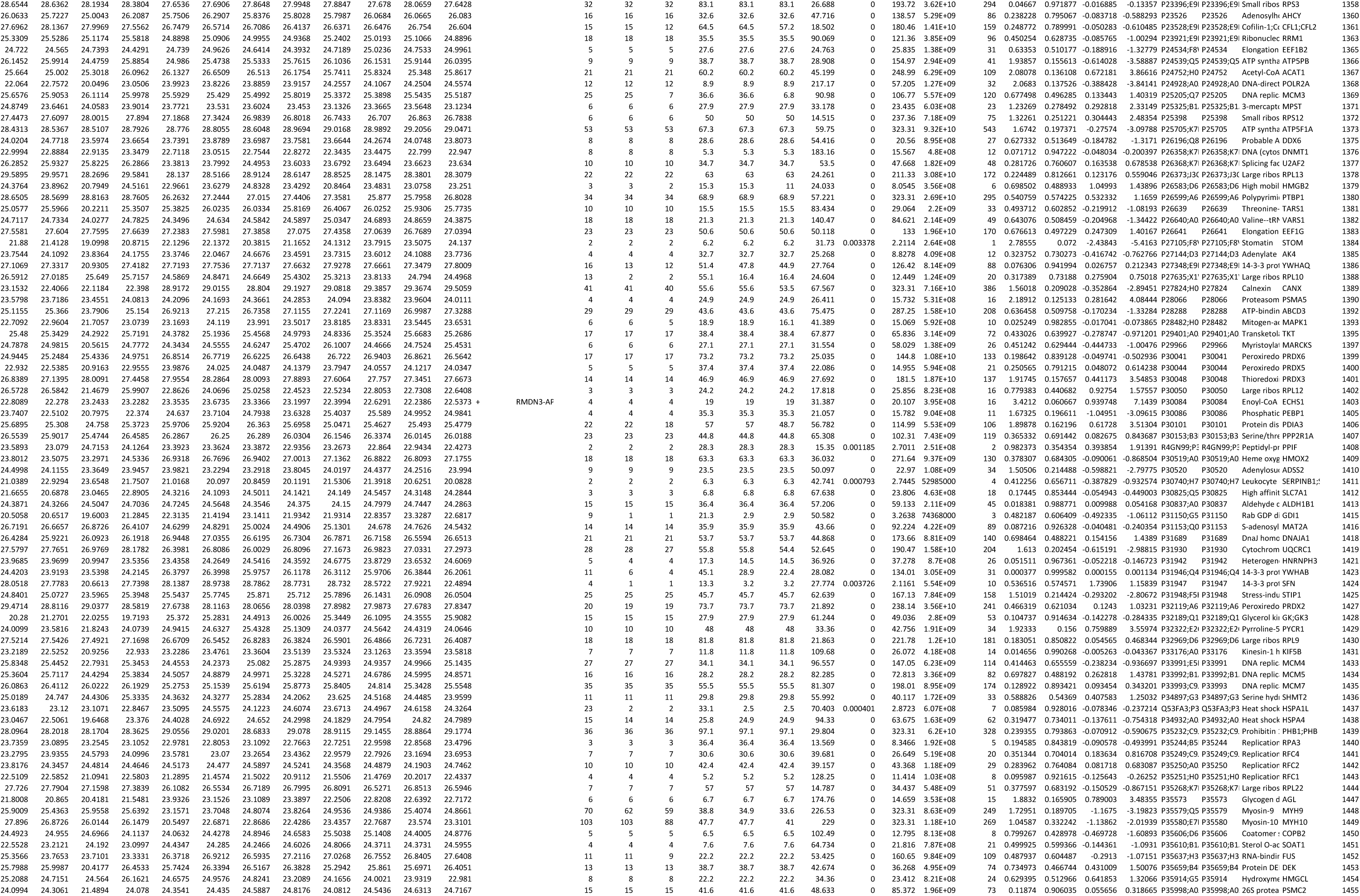
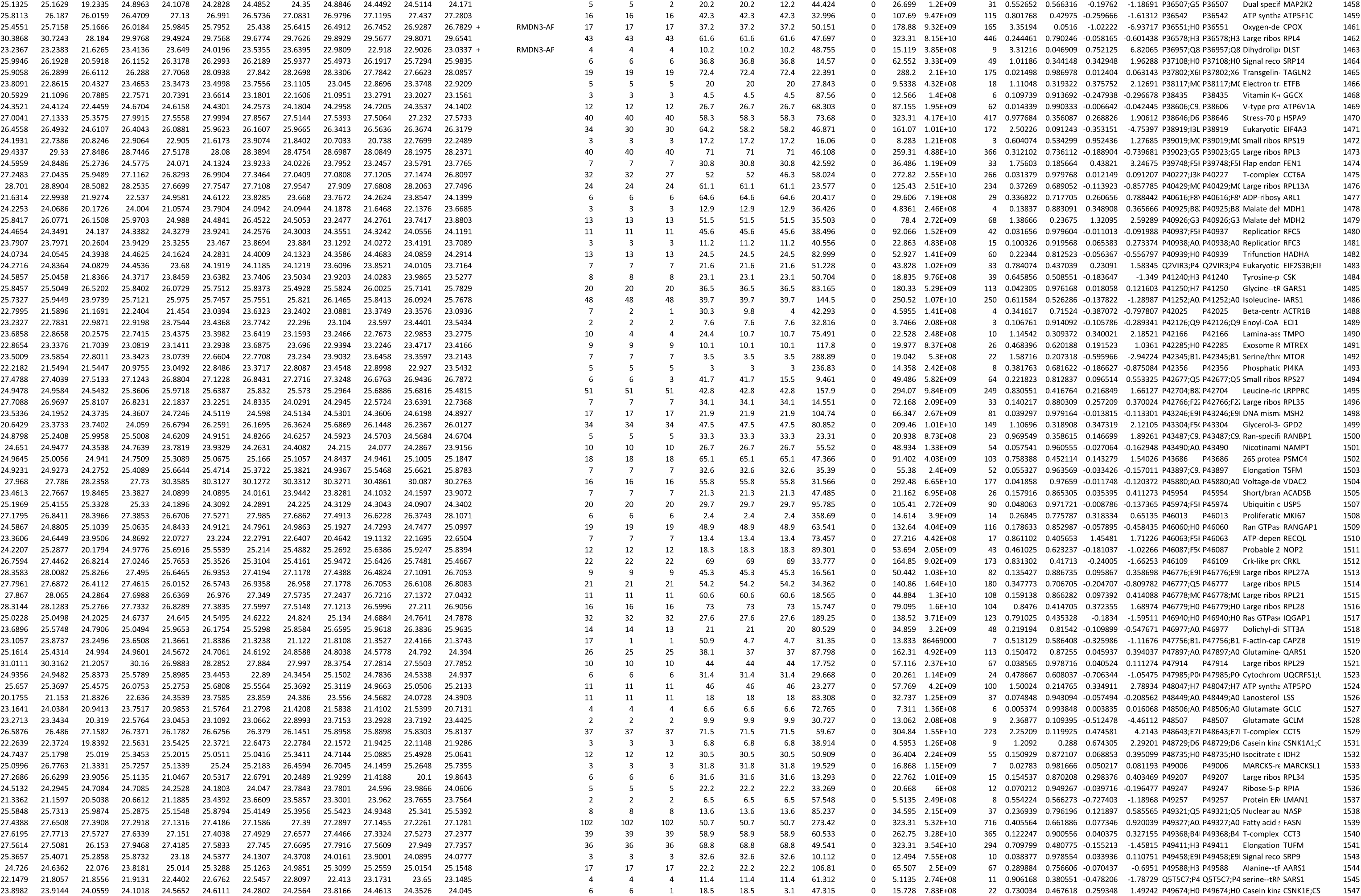
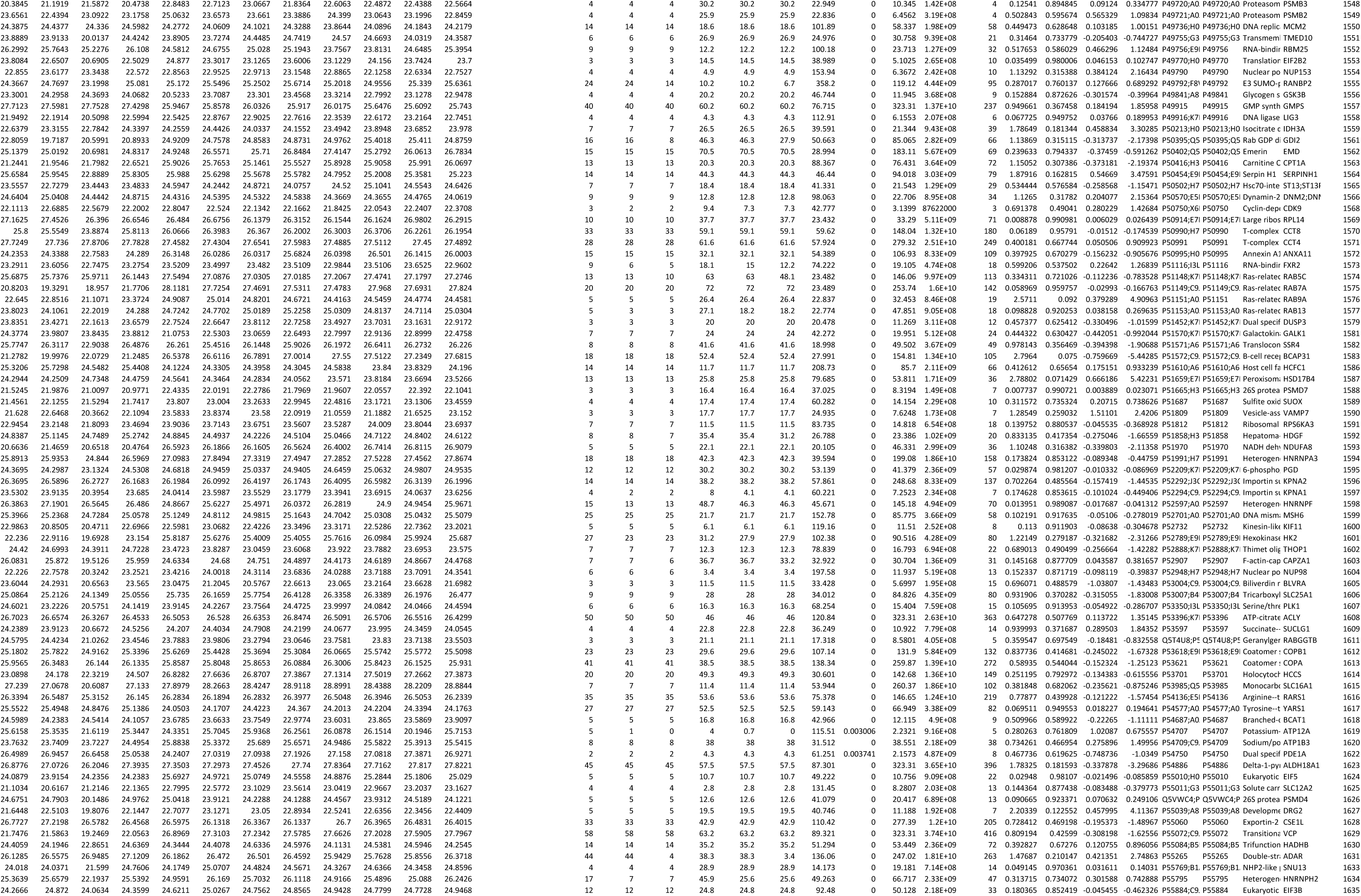
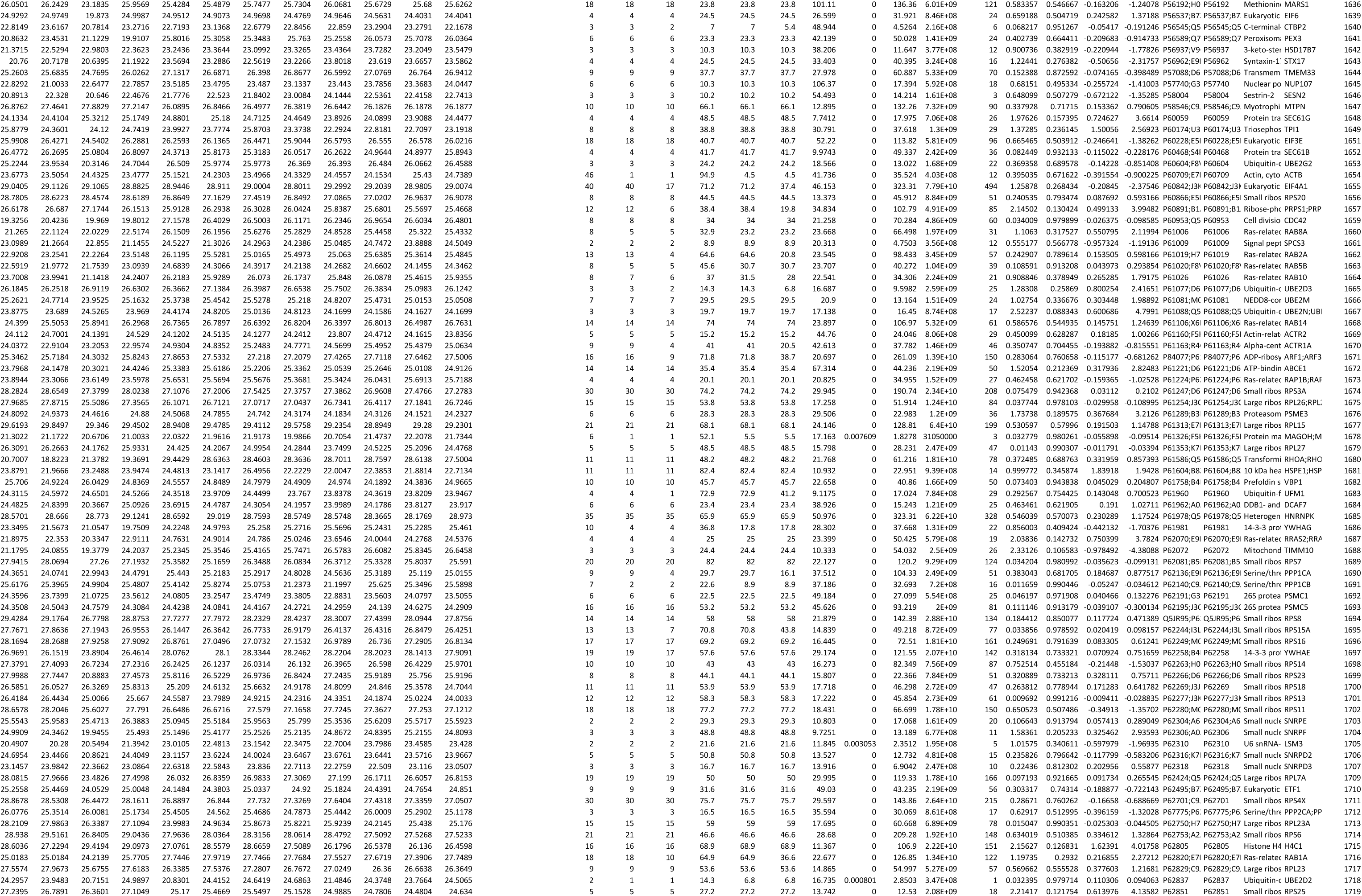
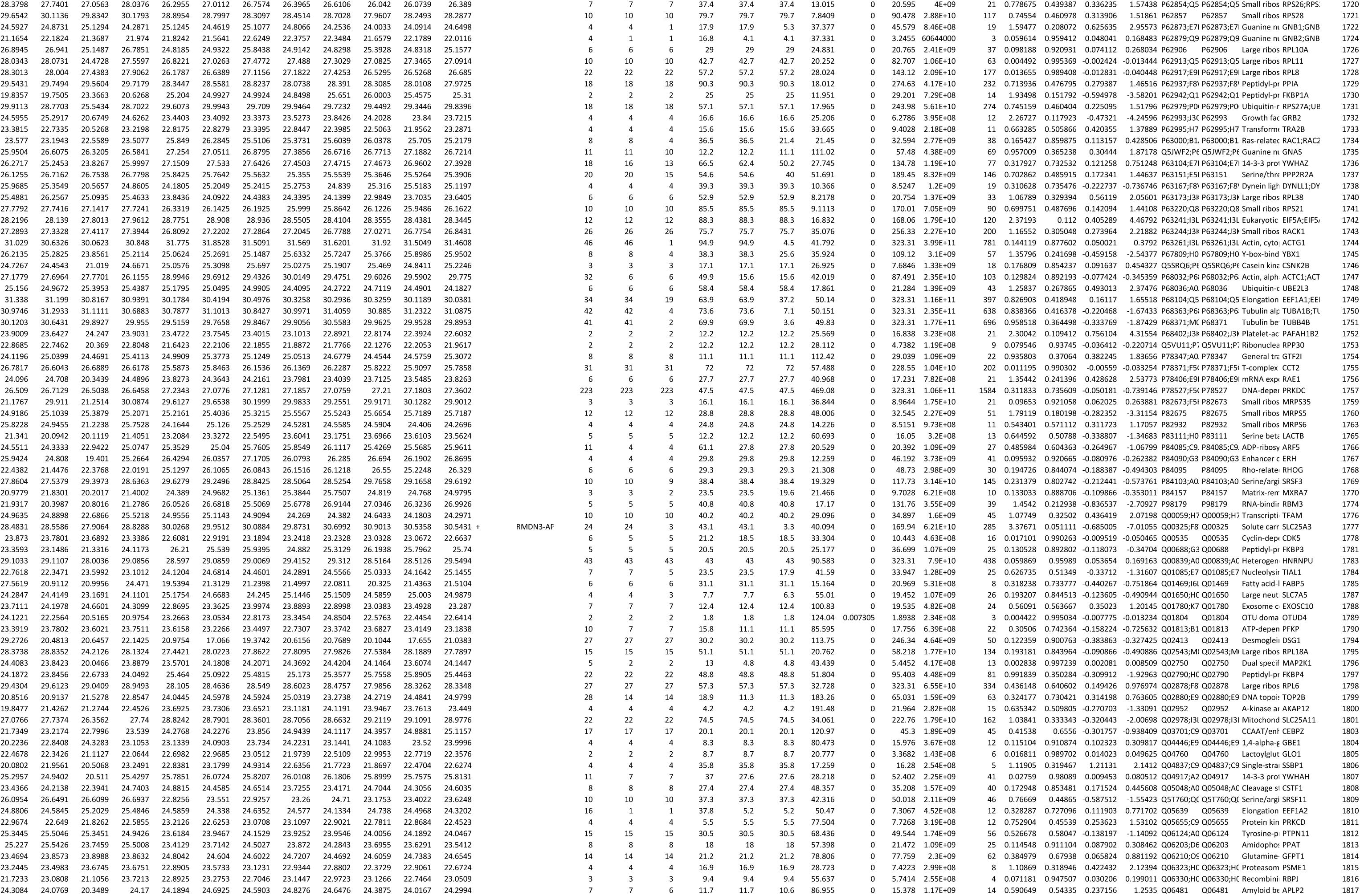
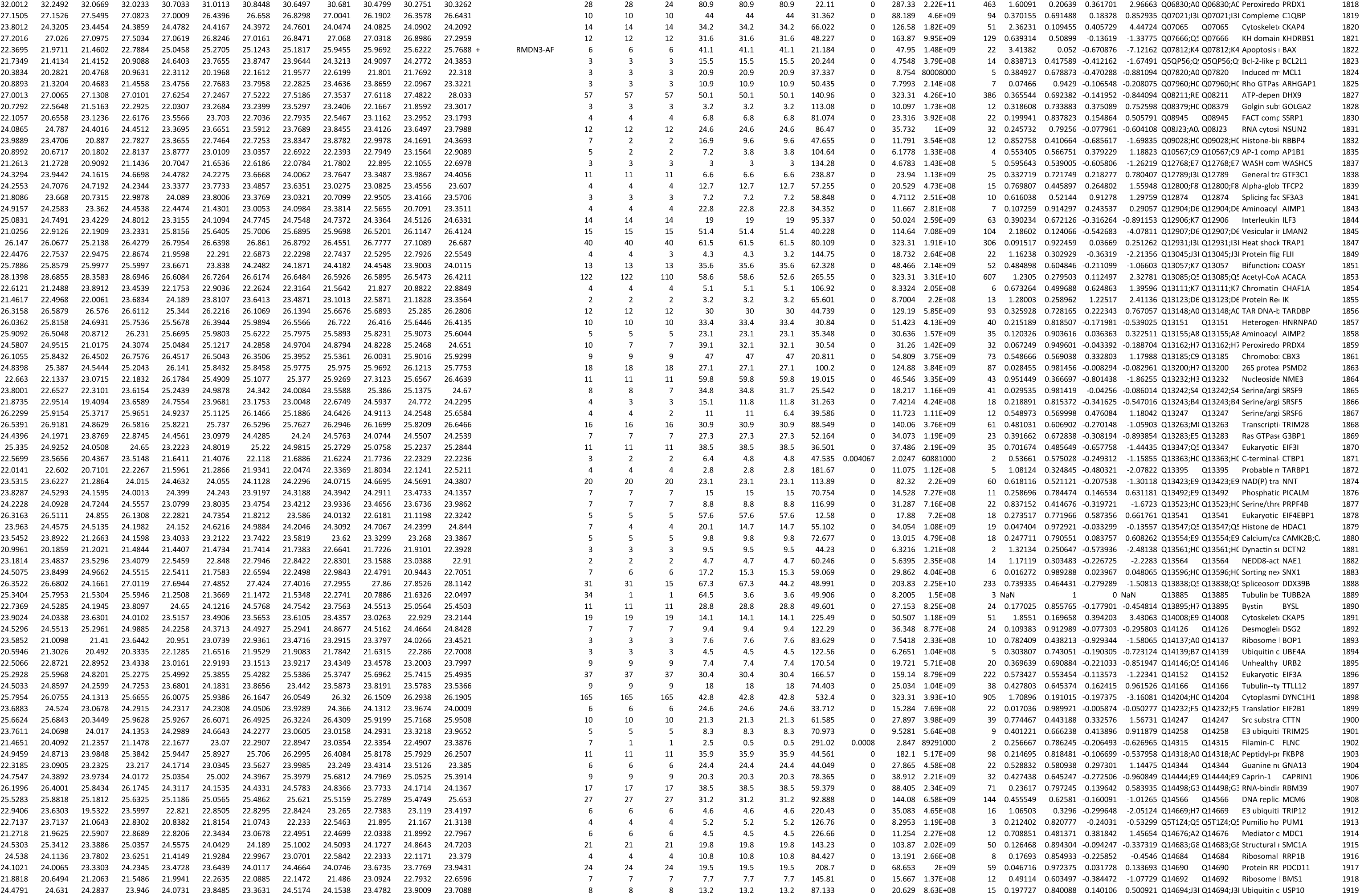
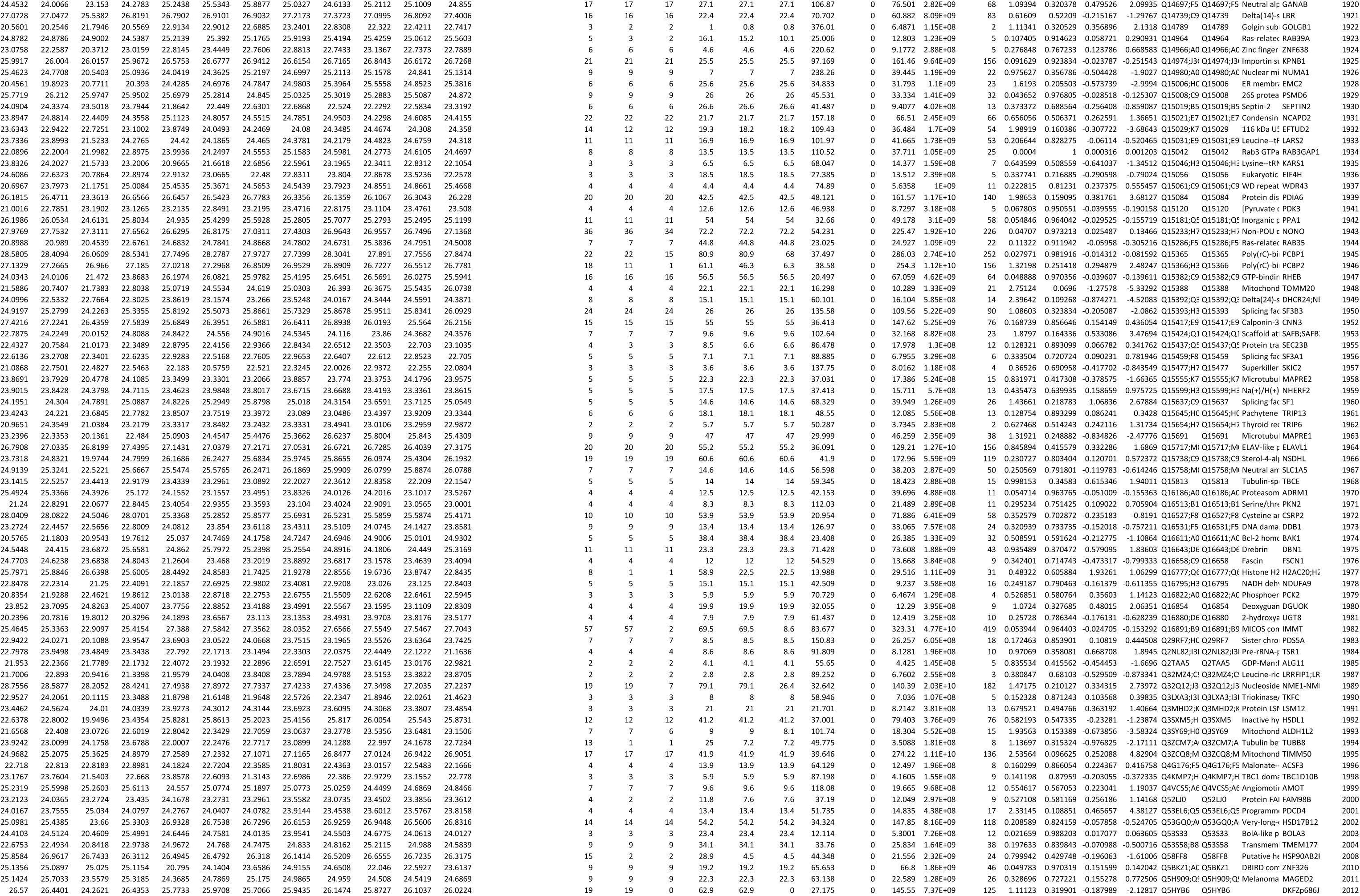
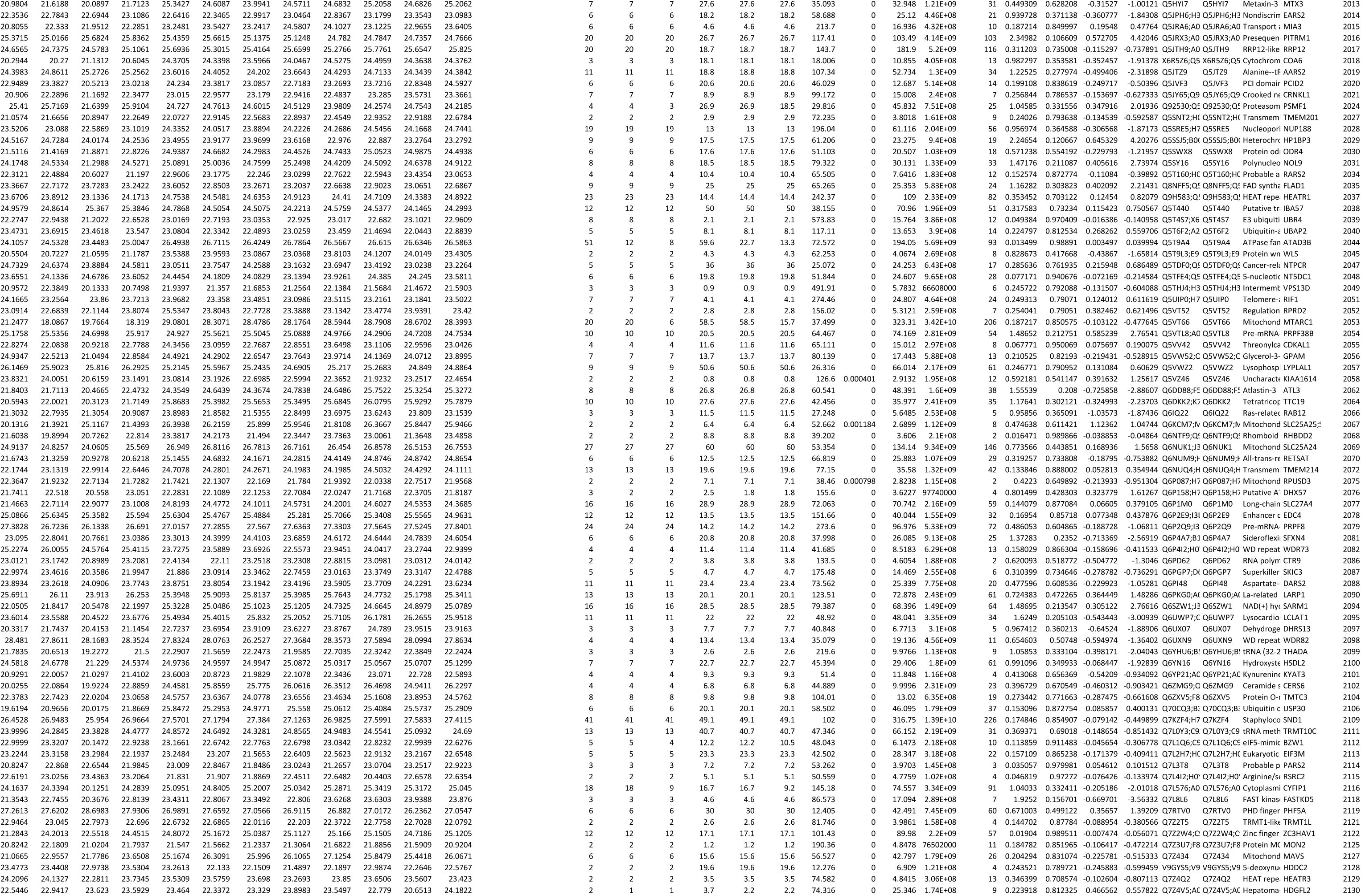
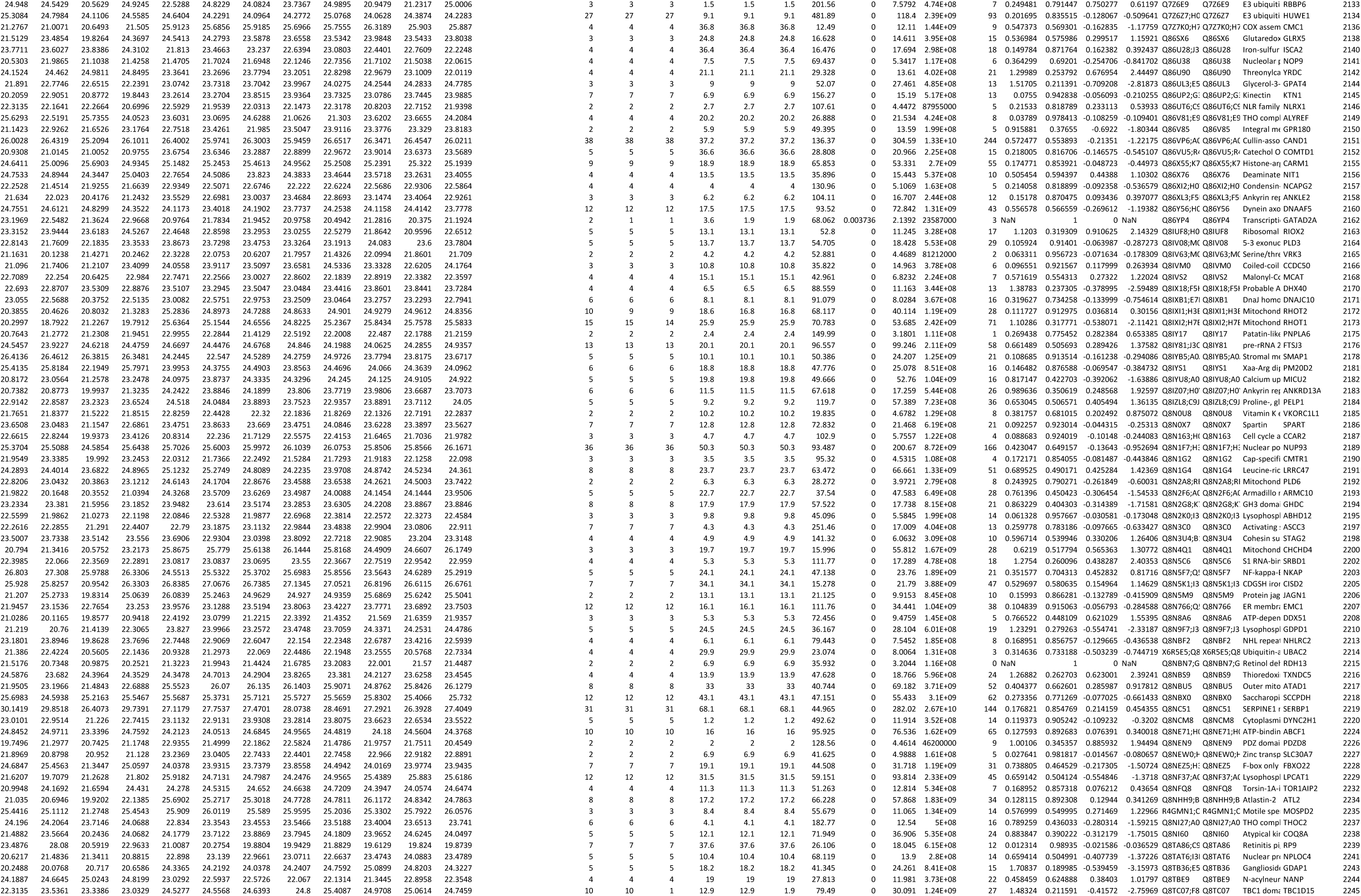
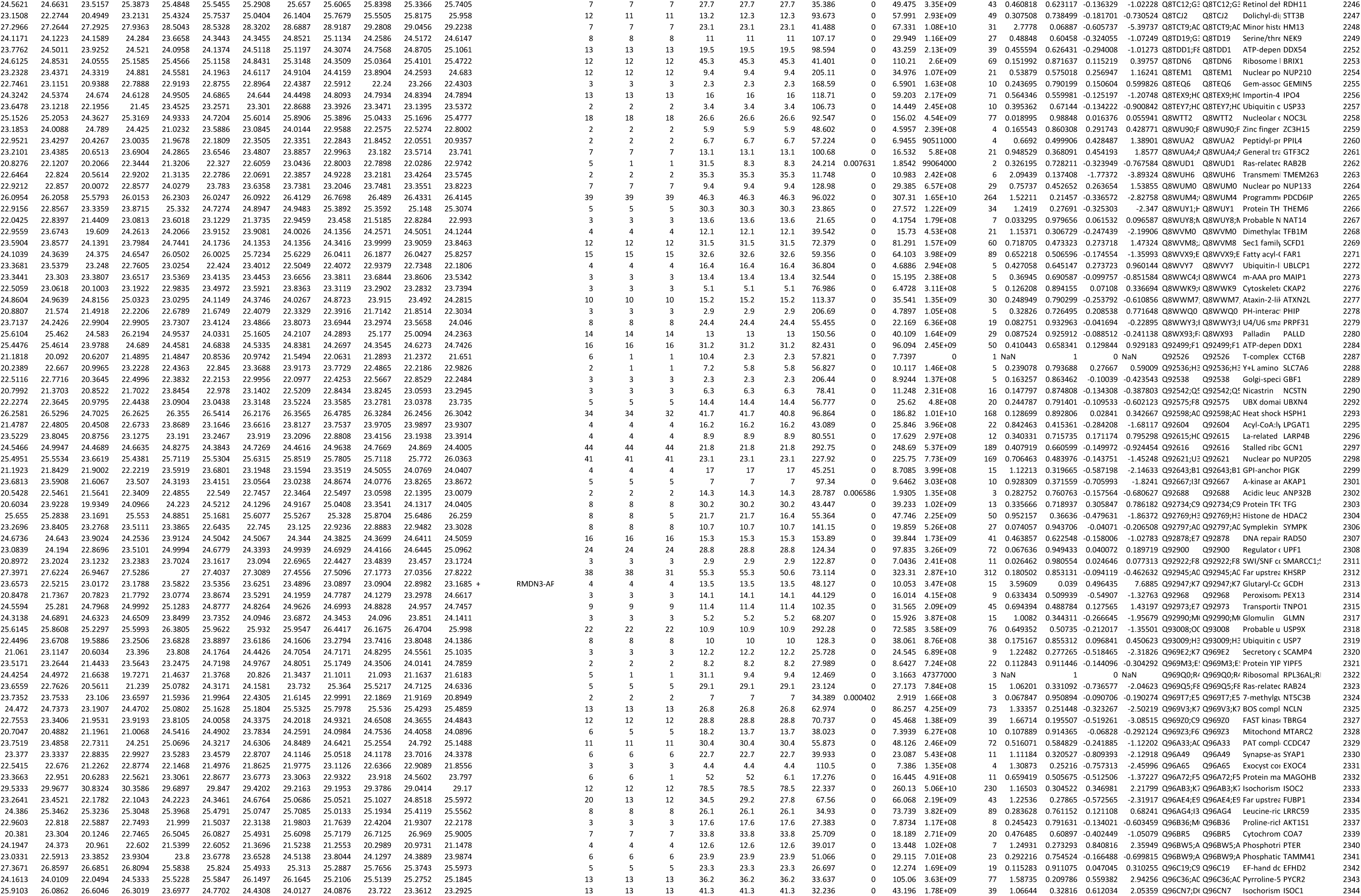
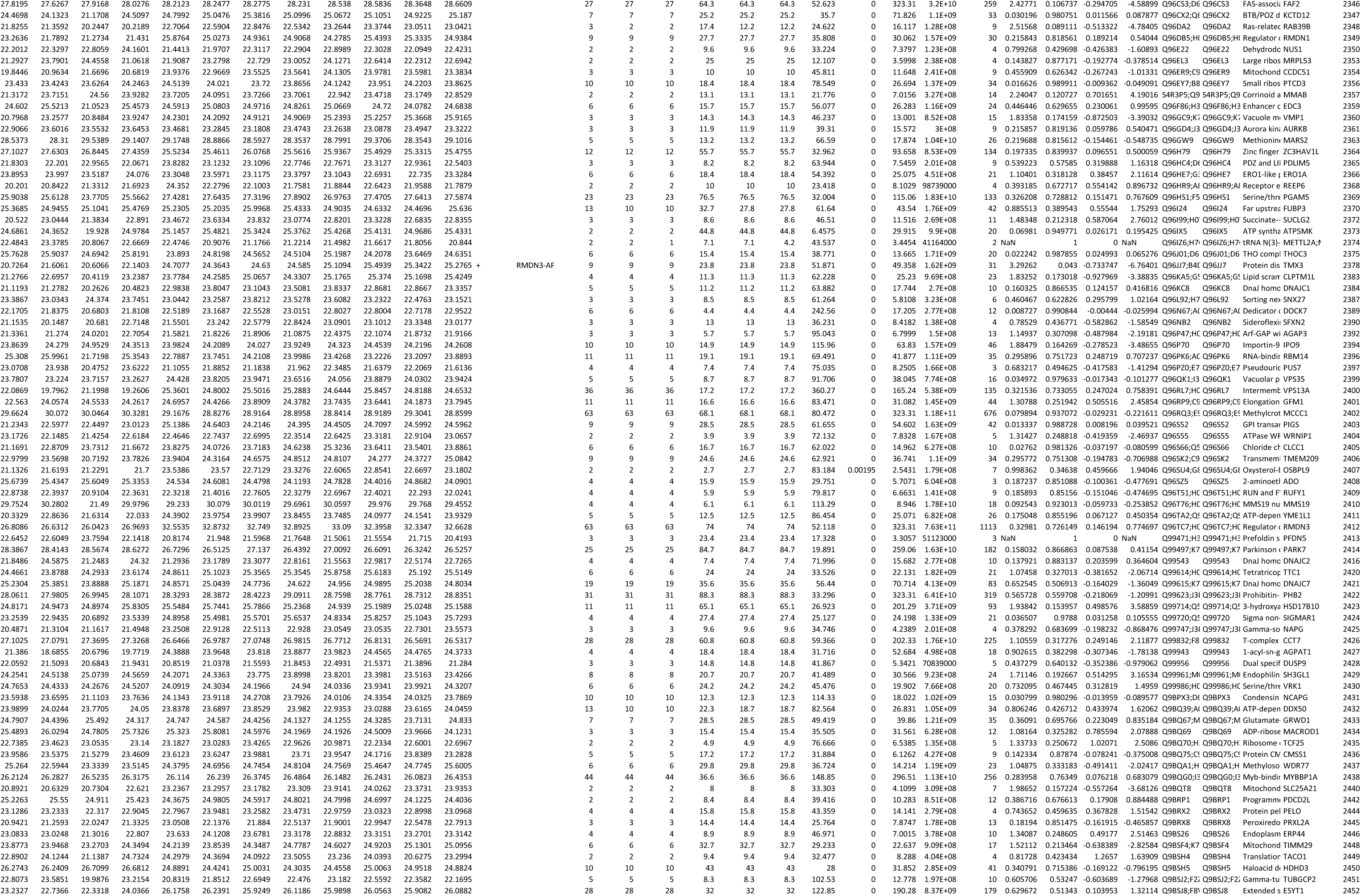
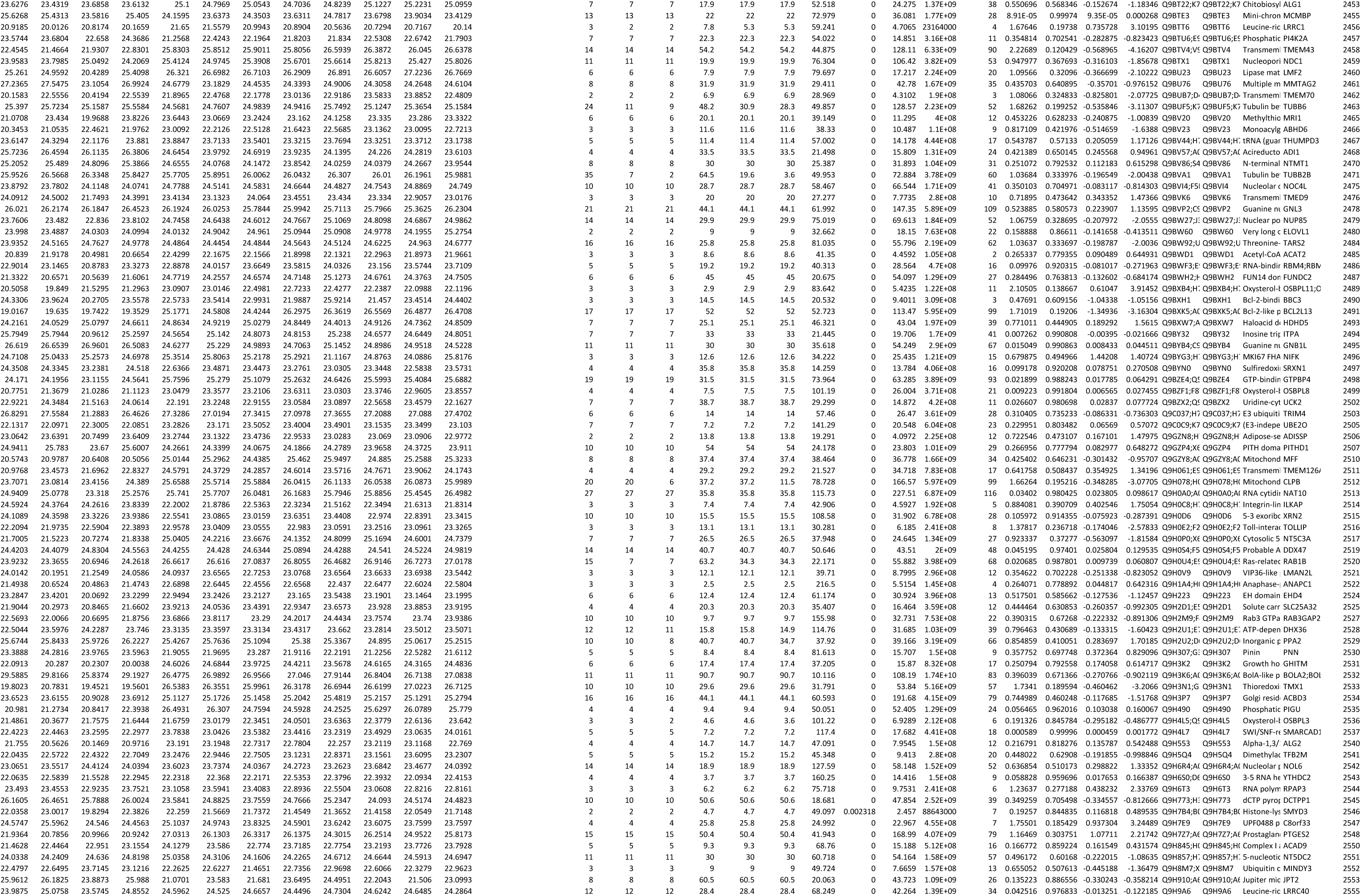
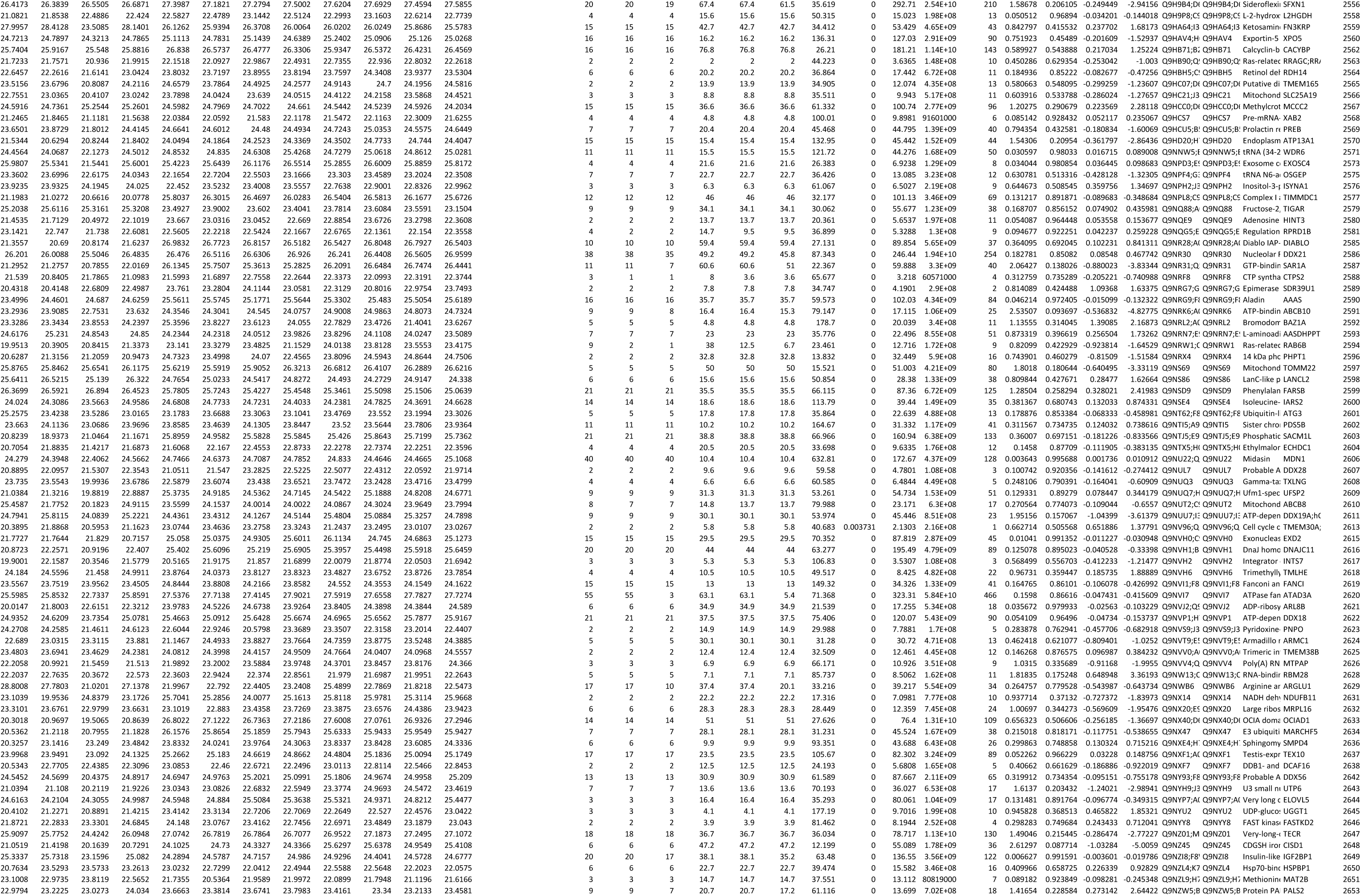
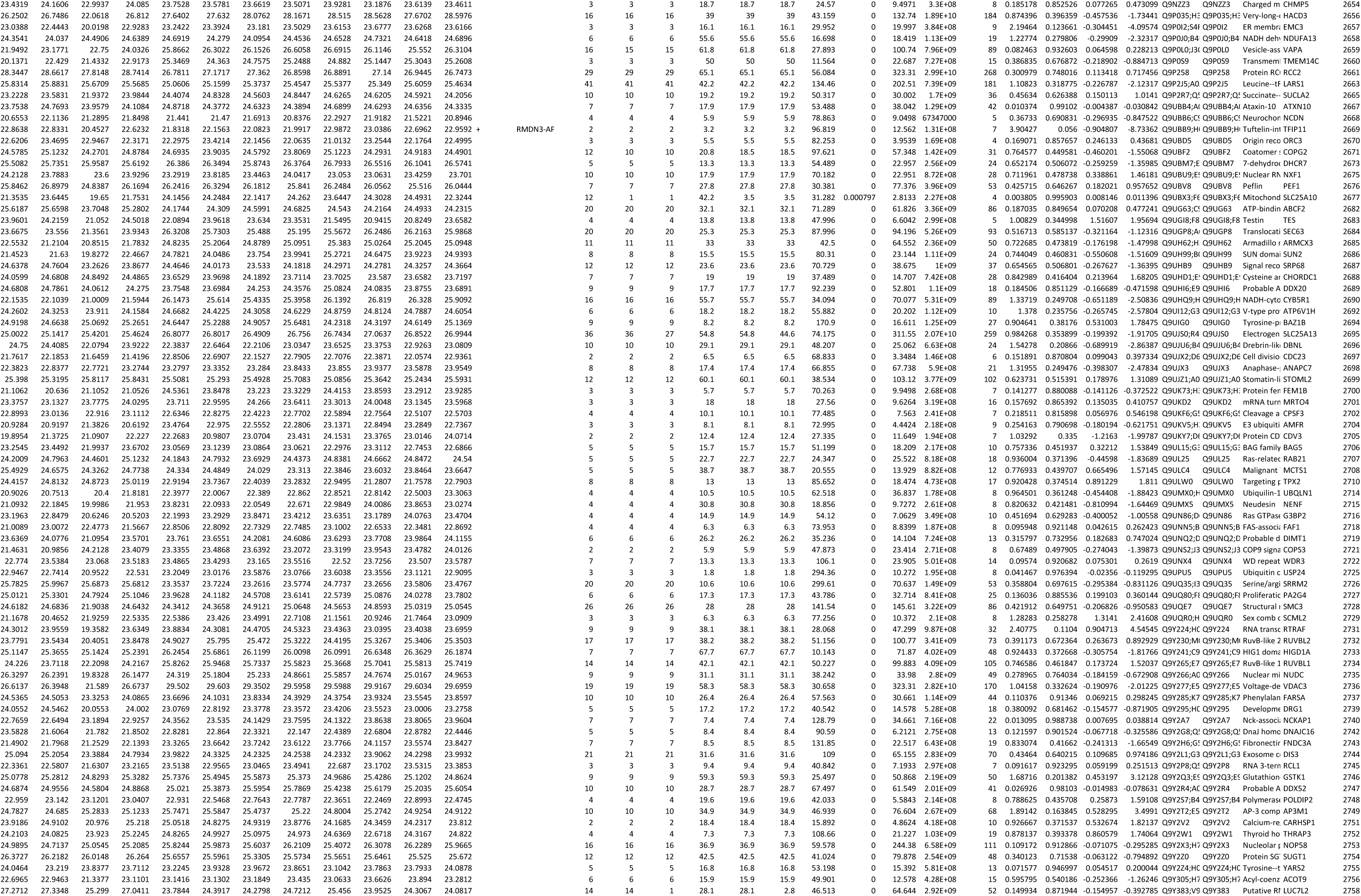
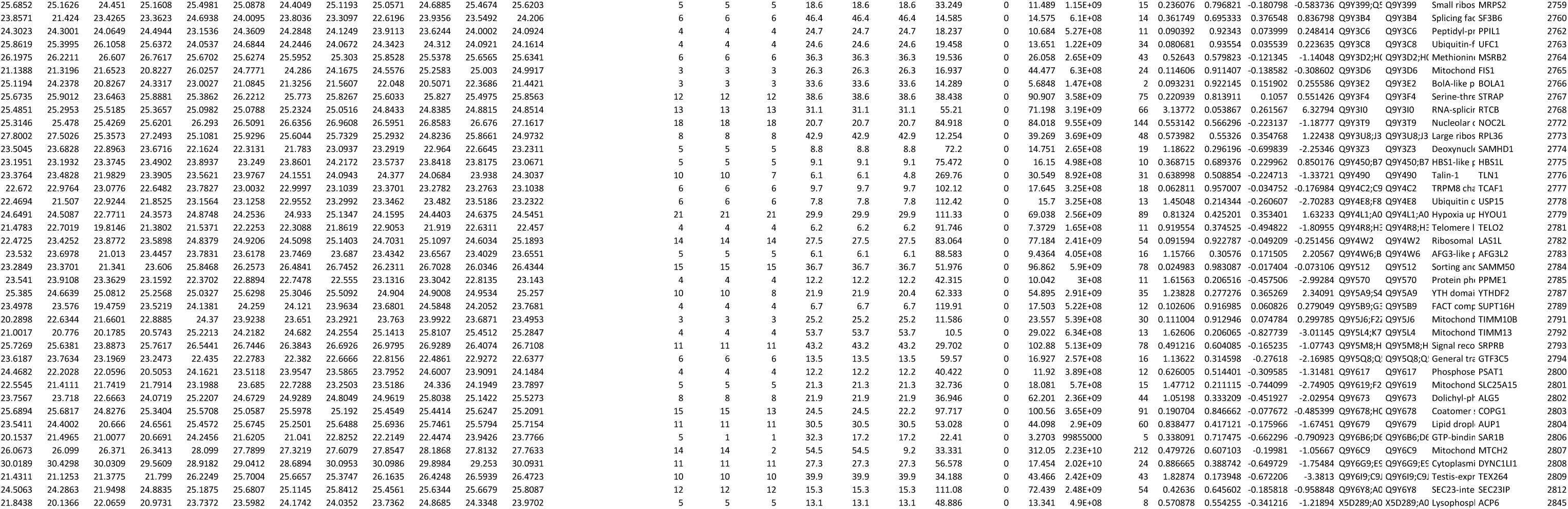

